# Metabolic reprogramming in the spinal cord drives the transition to pain chronicity

**DOI:** 10.1101/2025.01.30.635746

**Authors:** Alex Mabou Tagne, Yannick Fotio, Hye-Lim Lee, Kwang-Mook Jung, Jean Katz, Faizy Ahmed, Johnny Le, Richard Bazinet, Cholsoon Jang, Daniele Piomelli

## Abstract

Acute injuries can progress into painful states that endure long after healing. The mechanisms underlying this transition remain unclear, but metabolic adaptations to the bioenergy demands imposed by injury are plausible contributors. Here we show that peripheral injury activates AKT/mTORC1 in afferent segments of the mouse spinal cord, redirecting local core metabolism toward biomass production while simultaneously suppressing autophagy-mediated biomass reclamation. This metabolic shift supports neuroplasticity, but creates a resource bottleneck that depletes critical spinal cord nutrients. Preventing this depletion with a modified diet normalizes biomass generation and autophagy and halts the transition to chronic pain. This effect, observed across multiple pain models, requires activation of the nutrient sensors, sirtuin-1 and AMPK, as well as restoration of autophagy. The findings identify metabolic reprogramming and consequent autophagy suppression as key drivers of the progression to pain chronicity and highlight nutritional and pharmacological interventions that could prevent this progression after surgery or other physical traumas.

## Introduction

Acutely painful injuries can transition into intractable pain states that persist long after tissue healing is complete^1,2^. Invasive surgeries are a striking example of this progression, with 30-50% of patients who undergo thoracotomy or mastectomy still reporting pain one year after an otherwise successful procedure^3,4^. Similarly, up to 30% of people who experience accidental physical trauma go on to develop persistent neck or back pain^5–7^. Stable neuroplastic modifications associated with central and peripheral sensitization underpin the emergence of chronic pain after injury^8–10^, but the molecular determinants driving these changes are still largely elusive^11,12^. Addressing this knowledge gap is crucial to identify regulatory checkpoints in the transition to pain chronicity, which could be targeted by therapy.

Metabolic adaptations to the energetic challenges posed by injury are plausible contributors to the development of chronic pain. Synaptic transmission in the central nervous system (CNS) consumes a formidable amount of bioenergy, primarily generated through aerobic glycolysis and mitochondrial respiration^13,14^. ATP, the product of these processes, powers action potentials, fuels transmitter release and vesicle recycling, and sustains synapse maintenance and remodeling. When a peripheral organ is damaged, bioenergy demands increase markedly as neurons and glia in afferent segments of the spinal cord must allocate their finite resources to two opposing tasks: supporting enhanced neural activity while simultaneously generating the biomass needed to establish central sensitization. Three neuroplastic modifications associated with the latter phenomenon – neuronal hyperexcitability^8^, long-term heightening of synaptic strength^8^, and accrued dendritic spine dynamics^15,16^ – are, in fact, critically dependent on the production of new proteins and lipids^17–21^. How spinal cord neurons and glia balance these conflicting demands is unclear. However, one possibility is that they reroute their core metabolism toward aerobic glycolysis – a pathway that, while less energy-efficient than mitochondrial respiration, produces essential precursors for the biosynthesis of nucleotides, amino acids, and fatty acids^22^.

Results from animal studies support the possibility that somatic injury reprograms metabolism in the spinal cord. For instance, hind-paw administration of formalin in rats and chronic constriction injury (CCI) of the sciatic nerve in mice promote central sensitization and lasting pain through a mechanism that requires activation of the metabolic controller mammalian target of rapamycin complex 1 (mTORC1) and its upstream regulator AKT^23–25^. Furthermore, hind-paw formalin injection in mice stimulates both the expression of glycolytic enzymes and the accumulation of glycolysis metabolites in lumbar spinal cord segments ipsilateral to the lesion, suggesting that glycolysis might be enhanced^26^. Krebs’ cycle and oxidative phosphorylation components are concomitantly reduced, leading to a pronounced decrease in ATP levels. Importantly, this metabolic shift peaks 4 days after injury and coincides with a decline in the local concentrations of various amino acids, fatty acids, and the fatty acyl derivative palmitoylethanolamide (PEA). PEA is an endogenous agonist of peroxisome proliferator-activated receptor (PPAR)-α^27^, a key transcriptional regulator of mitochondrial respiration^28^. During this critical time window, but not before or after, PPAR-α activation halts the progression to pain chronicity^26^. However, it remains unclear whether the injury-induced metabolic shift and nutrient deficit independently drive this progression or represent an epiphenomenon.

To address this question, in the present study we examined whether preserving normative levels of amino acids, fatty acids, and PEA in the spinal cord prevents the emergence of chronic pain following injury. We found that a modified diet (MD-1) that counters the injury-induced shortfall in nutrients and PEA stops both metabolic reprogramming and chronic pain development following hind-paw formalin injection in male and female mice. Similar effects were observed in three additional models of injury-induced pain – CCI^29^, spared nerve injury (SNI)^30^, and surgical paw incision (SPI)^31^ – but not in the complete Freund’s adjuvant (CFA) model of immune-induced pain. Importantly, a nutrient-matched diet isocaloric with MD-1 but lacking PEA or a diet enriched solely in PEA was not protective, indicating that the effects of MD-1 cannot be attributed to nutrients or PEA alone, but rather to a synergistic interaction between these two factors. In mechanistic investigations, we found that, in mice fed a standard diet, hind-paw damage activates AKT/mTORC1 signaling in afferent segments of the spinal cord, enhancing biomass production and suppressing autophagy. MD-1 normalizes both processes and simultaneously stops the development of chronic pain by engaging the nutrient sensors, sirtuin-1 (SIRT1) and AMP-activated protein kinase (AMPK)^32,33^. Together, these results identify critical injury-induced metabolic alterations in spinal cord that drive the transition to pain chronicity. Importantly, our finding that pharmacological and nutritional interventions that correct these alterations prevent the transition from acute to chronic pain in mice, suggest that this approach could be adopted in the clinic to prevent the transition to chronic pain following invasive surgery or other forms of physical trauma.

## Results

### Peripheral injury promotes nutrient depletion in the spinal cord and MD-1 averts it

Consistent with previous findings^26^, hind-paw formalin injection induced tissue-wide metabolic reprogramming in ipsilateral lumbar (L4-L6) hemicords of male CD1 mice, which peaked four days post-lesion (Supplemental Fig. S1)^26^. Compared to saline injection, formalin administration upregulated the transcription of genes involved in glycolysis, while genes associated with Krebs’ cycle and oxidative phosphorylation were generally, though not uniformly, downregulated (Supplemental Fig. S1A, B). Notable exceptions were succinate dehydrogenase (*Sdhb*, *Sdhd*) and ubiquinol-cytochrome c reductase (*Ucqrq*), whose transcription was elevated (Supplemental Fig. S1B) possibly due to their contribution to other consequences of injury, including suppression of autophagy and stimulation of apoptosis^34,35^ (see below). Furthermore, levels of glycolytic metabolites were higher (Supplemental Fig. S1C) and Krebs’ cycle metabolites were lower relative to vehicle controls (Supplemental Fig. S1D). Prior work has shown that this metabolic shift is associated with a substantial reduction in the local concentrations of 12 amino acids, 2 monounsaturated fatty acids [oleic acid (18:1D^9^) and erucic acid (22:1 D^13^)], and the endogenous PPAR-a agonist PEA^27^. To determine whether this depletion independently contributes to pain chronification, rather than being a byproduct, we formulated a modified diet (MD-1) enriched with the depleted substances (Supplemental Table S1).

Firstly, to assess the diet’s bioavailability, we fed male CD-1 mice MD-1 for 25 days and analyzed blood samples using liquid chromatography/mass spectrometry (LC/MS) (Figure 1A). Compared to mice on a standard diet (SD), mice fed MD-1 showed significantly higher serum concentrations of most supplemented compounds, including 8 amino acids, oleic acid, erucic acid, and PEA (Figure 1B). The concentrations of serine and phenylalanine remained unchanged, likely due to biotransformation, as indicated by higher levels of their catabolites, glycine (from serine) and phenylpyruvate (from phenylalanine) (Supplemental Fig. S2A, B). Elevated serum concentrations of various tryptophan and tyrosine catabolites confirmed that MD-1 components were integrated into core metabolism (Figure 1B, Supplemental Fig. S2C, D). The absence of detectable cysteine, along with unaltered cysteine catabolites, methionine, and methionine catabolite cystathionine (Supplemental Fig. S2E, F), may reflect incorporation into proteins or conversion into compounds not targeted in our analysis. Other serum metabolites influenced by MD-1 exposure are listed in Supplemental Table S2.

**Figure 1.**
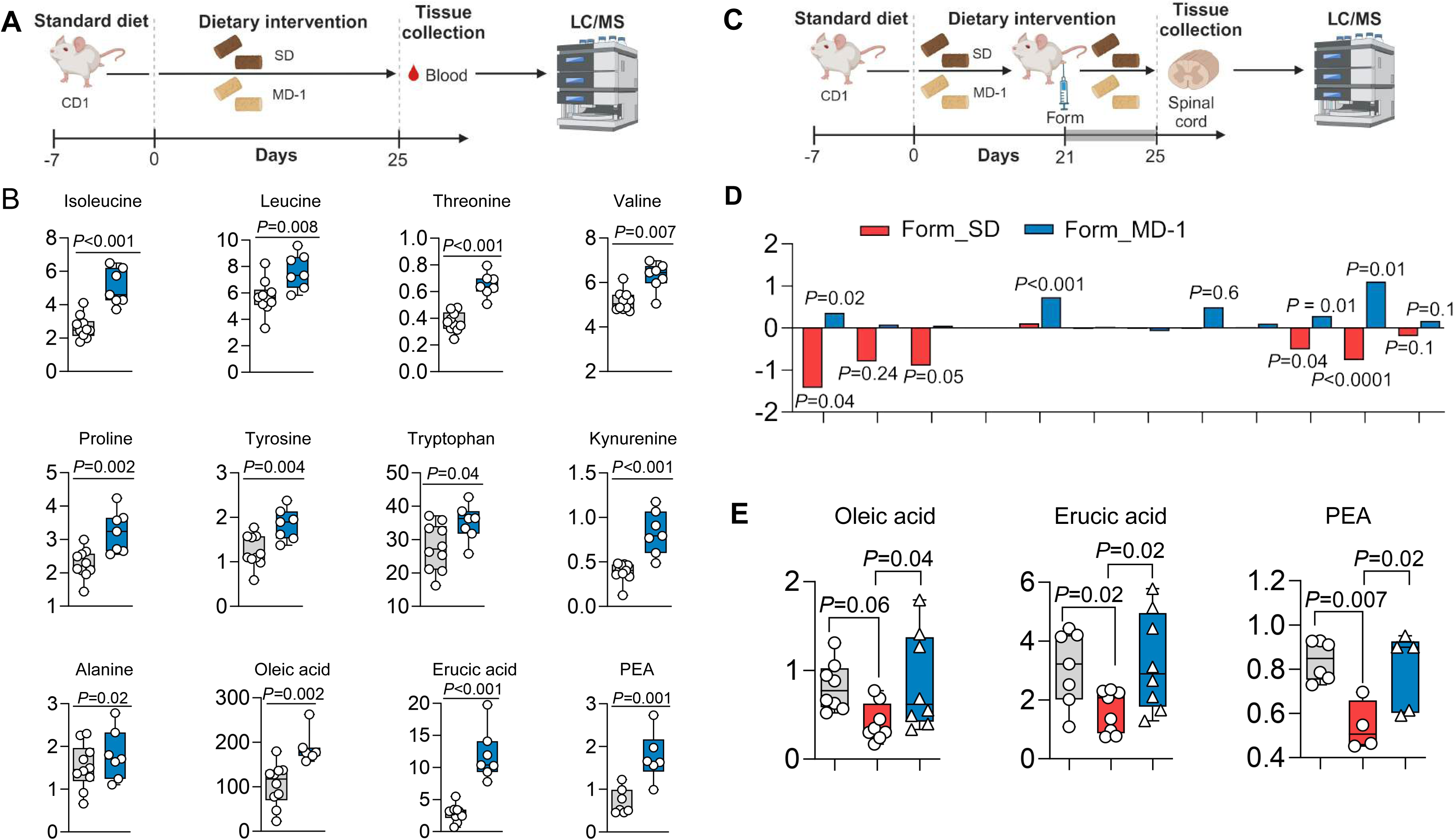
MD-1 is bioavailable and prevents injury-induced nutrient depletion in the spinal cord. (A) Protocol: mice received SD or MD-1 for 25 days and blood was collected for LC/MS analysis. (B) Serum concentrations (ion count) of MD-1 components in mice fed SD (gray boxes) or MD-1 (blue boxes). (C) Protocol: mice received SD or MD-1 for 21 days followed by formalin (Form) or saline (Veh) injection. Four days post-injection, ipsilateral L4-L6 spinal cord segments were collected for LC/MS analysis. (D) Effects of formalin and MD-1 (log_2_ fold changes) on amino acid levels. Red bars: formalin *vs* vehicle in SD-1-fed mice; blue bars; MD-1 *vs* SD feeding in formalin-injected mice. (E) Oleic acid, erucic acid, and PEA content in vehicle-injected mice fed SD (gray boxes) and formalin-injected mice fed SD (red boxes) or MD-1 (blue boxes). (B, D) Student’s *t* test, *n* = 6-10 per group; (E) One-way ANOVA and Šídák’s test, *n* = 7-8 per group.

We next examined whether MD-1 prevents the injury-induced nutrient deficit in ipsilateral lumbar hemicord tissue. CD-1 mice were fed MD-1 or SD for 21 days before receiving hind-paw injections of formalin in saline (1%, 20 mL) or saline alone. This model is commonly used to study acute pain^36^ but also produces a lasting pathological state that mirrors key aspects of severe chronic pain in humans^26^, including bilateral hypersensitivity to mechanical and thermal stimuli^37,38^, heightened anxiety^39^, cognitive impairments^40^, and structural CNS alterations^41^. Four days post-formalin injection, we harvested ipsilateral lumbar hemicords and quantified metabolites by LC/MS (Figure 1C). As previously observed in C57Bl6 mice^26^, formalin-treated, SD-fed mice exhibited lower levels of various amino acids, oleic acid, erucic acid, and PEA, compared to uninjured (vehicle-injected) controls (Figure 1D, E). No such decline was observed, however, in formalin-treated, MD-1-fed mice, where levels of these substances were either stable or slightly elevated compared to uninjured mice (Figure 1D, E). In the latter, MD-1 increased concentrations of PEA and 4 of the 12 amino acids supplemented by the diet (Supplemental Fig. S3). Other metabolomic changes elicited by MD-1 in ipsilateral lumbar hemicords of uninjured mice are detailed in Supplemental Table S3. MD-1’s ability to avert spinal nutrient depletion encouraged us to utilize this diet to investigate the relationship between injury-induced metabolic alterations and chronic pain development.

### MD-1 prevents injury-induced metabolic reprogramming in the spinal cord

Figure 2 illustrates the effects of MD-1, alone or in combination with formalin treatment, on the expression of genes associated with energy metabolism in ipsilateral lumbar hemicords, as assessed by bulk RNA-sequencing. In uninjured mice, MD-1 enhanced the transcription of glycolytic genes, including aldolase (*Aldob*), triosephosphate isomerase (*Tpi1*), and lactate dehydrogenase (*Ldha*, *Ldhb*) (Figure 2A). The diet also increased the expression of oxidative phosphorylation components, such as ATP synthase subunit g (*Atp5l*), cytochrome c oxidase subunit 6C2 (*Cox6c2*), *Sdh (a, b* and *d)*, and *Ucqrq1* (Figure 2B). Additionally, MD-1 induced a slight downward trend in the transcription of Krebs’ cycle-related genes, with citrate synthase (*Cs*) showing borderline statistical significance (P = 0.05) (Figure 2B). However, these transcriptional modifications, along with others reported in Supplemental Table S4, were not accompanied by detectable changes in levels of glycolysis or Krebs’ cycle metabolites (Supplemental Fig. S4). In contrast, MD-1 effectively counteracted all molecular changes caused by formalin injection in ipsilateral lumbar hemicords, blocking the upregulation of glycolytic genes and the downregulation of genes involved in Krebs’ cycle and oxidative phosphorylation (Figure 2C, D and Supplemental Fig. S5; see Supplemental Table S4 for statistical analyses) as well as the shifts in glycolysis and Krebs’ cycle metabolites (Figure 2E). Notably, MD-1 preserved normative ATP levels in formalin-treated mice without altering them in controls (Figure 2F). Changes in other purines, which were variably affected by the lesion, were also blocked by MD-1 (Figure 2G). The results suggest that MD-1 has limited impact on bioenergy production in uninjured mice but averts metabolic reprogramming after injury.

**Figure 2.**
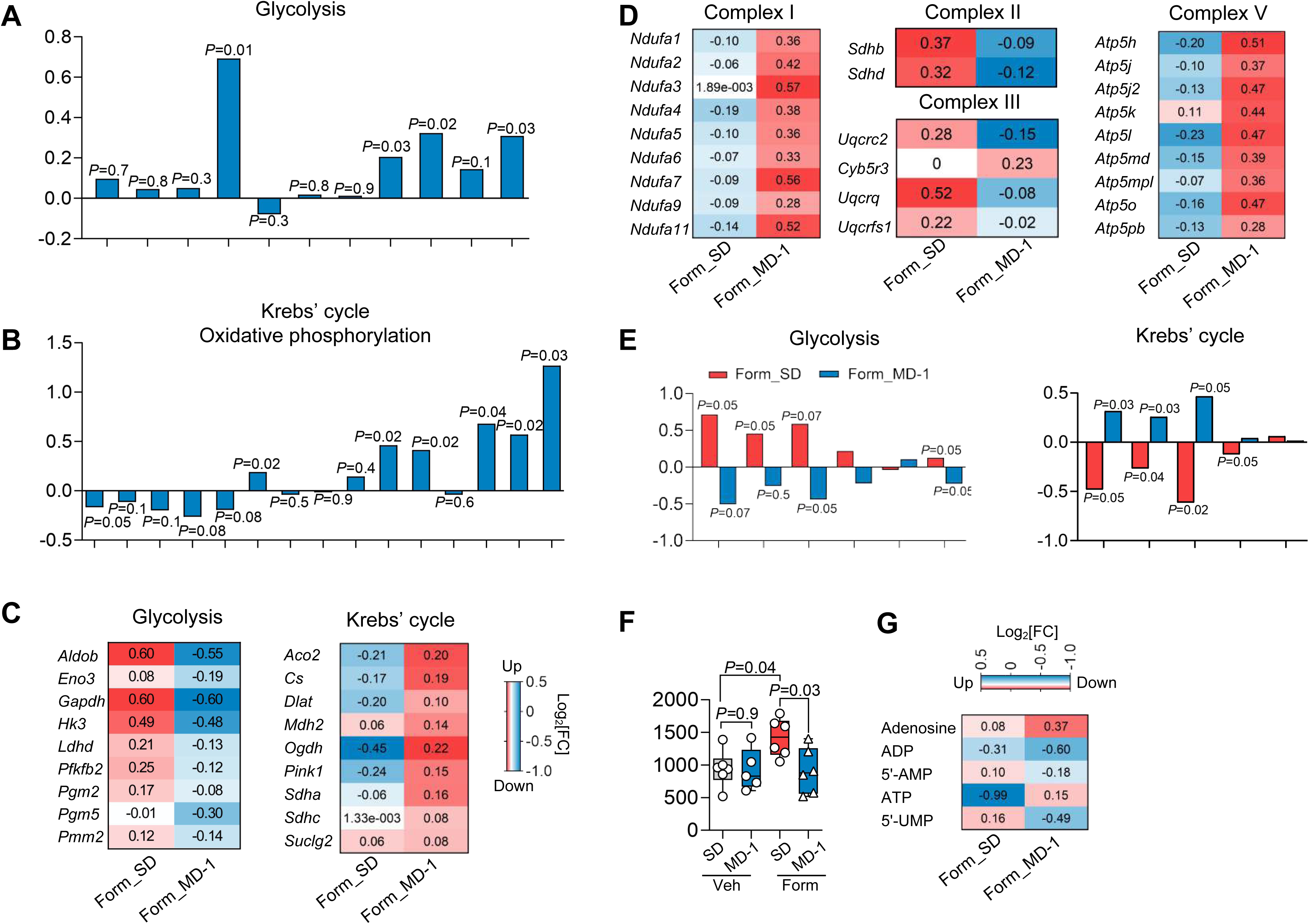
MD-1 prevents injury-induced metabolic reprogramming in the spinal cord. (A, B) Effects of MD-1 on the transcription of genes related to (A) glycolysis and (B) Krebs’ cycle and oxidative phosphorylation. Log_2_ fold changes in MD-1-fed *vs* SD-fed vehicle-injected mice. (C-D) Effects of formalin and MD-1 (log_2_ fold changes) on the transcription of genes related to (C) glycolysis and Krebs’ cycle, and (D) oxidative phosphorylation. Form_SD: formalin *vs* vehicle injection in SD-1-fed mice; Form_ MD-1: MD-1 *vs* SD feeding in formalin-injected mice. (E) Effects of formalin and MD-1 (log_2_ fold changes) on the concentrations of glycolysis (left) and Krebs’ cycle (right) metabolites. Form_SD: formalin *vs* vehicle injection in SD-1-fed mice; Form_ MD-1: MD-1 *vs* SD feeding in formalin-injected mice. Abbreviations: G6P, glucose-6-phosphate; F6P, fructose-6-phosphate; F1,6P, fructose-1-6-bisphosphate; DHAP, dihydroxyacetone phosphate; Pyr, pyruvate; Lac, lactate; Cit, citrate; Iso, isocitrate, Succ, succinate; Fum, fumarate; Mal, malate. (F) AMP/ATP ratio in vehicle (Veh)- or formalin (Form)-injected mice fed SD or MD-1. (G) Effects of formalin and MD-1 (log_2_ fold changes) on nucleoside and nucleotide concentrations. Form_SD: formalin *vs* vehicle injection in SD-1-fed mice; Form_ MD-1: MD-1 *vs* SD feeding in formalin-injected mice. (A, B, E) Multiple unpaired *t* test, *n* = 6-10 per group; (F) One-way ANOVA and Šídák’s test, *n* = 5-6 per group.

The serine/threonine kinase AKT and its downstream target mTORC1 are pivotal regulators of energy metabolism^42,43^. They are also required for the induction of central sensitization and chronic pain following hind-paw formalin injection in rats or sciatic nerve injury in mice^23–25^. In line with this role, formalin administration in SD-fed mice increased AKT and mTOR phosphorylation (activation) in ipsilateral lumbar hemicords, as assessed by Western blot analysis (Supplemental Fig. S6A-C). This activation coincided with enhanced transcription of phosphoinositide-3-kinase catalytic subunit-g (*Pik3cg*) (Supplemental Fig. S6D), which facilitates AKT recruitment^44^, and reduced transcription of regulated in development and DNA damage 1 (*Redd1*), an endogenous AKT/mTORC1 inhibitor^45^, as shown by bulk RNAseq (Supplemental Fig. S6E). Additionally, PPAR-α (*Ppara*) and peroxisome proliferator-activated receptor-g coactivator-1α (*PGC1*a)—key controllers of mitochondrial respiration^46^ that are repressed by mTORC1^47^—were downregulated in formalin-treated mice, compared to uninjured controls (Supplemental Fig. S6F, G), whereas PPAR-γ (*Pparg*) and PPAR-d (*Ppard*) remained unchanged (Supplemental Fig. S6H). Notably, MD-1 prevented these alterations (Supplemental Fig. S6A-H) and decreased baseline AKT phosphorylation in uninjured mice (Supplemental Fig. S6A, B).

To determine whether AKT and mTORC1 are involved in chronic pain development, we treated SD-fed mice with the AKT inhibitor MK-2206 (240 mg/kg, intraperitoneal, IP)^48^ or the mTORC1/2 inhibitor Torin-1 (20 mg/kg, IP)^49^ once daily on days 2, 3, and 4 post-formalin (Supplemental Fig. S7A). Both inhibitors stopped the development of persistent bilateral hypersensitivity (Supplemental Fig. S7B, C), confirming that AKT/mTORC1 signaling is necessary to establish central sensitization and chronic pain^23–25^. These data, along with the known anabolic functions of mTORC1^42,43^, led us to hypothesize that injury-induced mTORC1 activation drives the progression to pain chronicity by affecting biomass production and autophagy.

### Injury enhances biomass production and suppresses autophagy in the spinal cord, and MD-1 prevents these effects

The neuroplastic adaptations that underlie central sensitization depend on the synthesis of synaptic proteins and lipids^17–21^. Accordingly, bulk RNAseq studies showed that the transcription of numerous genes involved in the production of neuroglial biomass was enhanced in ipsilateral lumbar hemicords of SD-fed, formalin-treated mice relative to uninjured controls (Figure 3; see Supplemental Table S5 for statistical analyses). For example, there was an upregulation of genes encoding voltage-gated ion channels (e.g., *Kcna*, *Scn*, *Cacna1*), receptor channels (e.g., *Gria*, *Gabra*, *Trpc*), and motor proteins (e.g., *Dyn1*, *Kif*) (Figure 3A-C) along with genes required for the synthesis of glycerophospholipids, sphingolipids, and cholesterol (Figure 3D-F). These transcriptional modifications aligned with accrued levels of glycerophospholipids (Figure 3G), sphingolipids (Figure 3H, I), and desmosterol (Figure 3J), as determined by metabolomics. Diacylglycerols were concomitantly decreased (Figure 3K), likely due to their use in phospholipid biosynthesis^50^. In contrast, MD-1-fed mice were strikingly resilient to the anabolic stimulation evoked by injury. In formalin-treated mice receiving MD-1, the diet prevented the upregulation of genes encoding ion channels, motor proteins, and lipid synthetic enzymes (Figure 3A-F), and stabilized tissue concentrations of all lipid classes (Figure 3G-K). The results indicate that injury stimulates the production of neuroplasticity-associated proteins and lipids in ipsilateral hemicords, likely under the control of mTORC1^42,43^, and MD-1 blocks this response.

**Figure 3.**
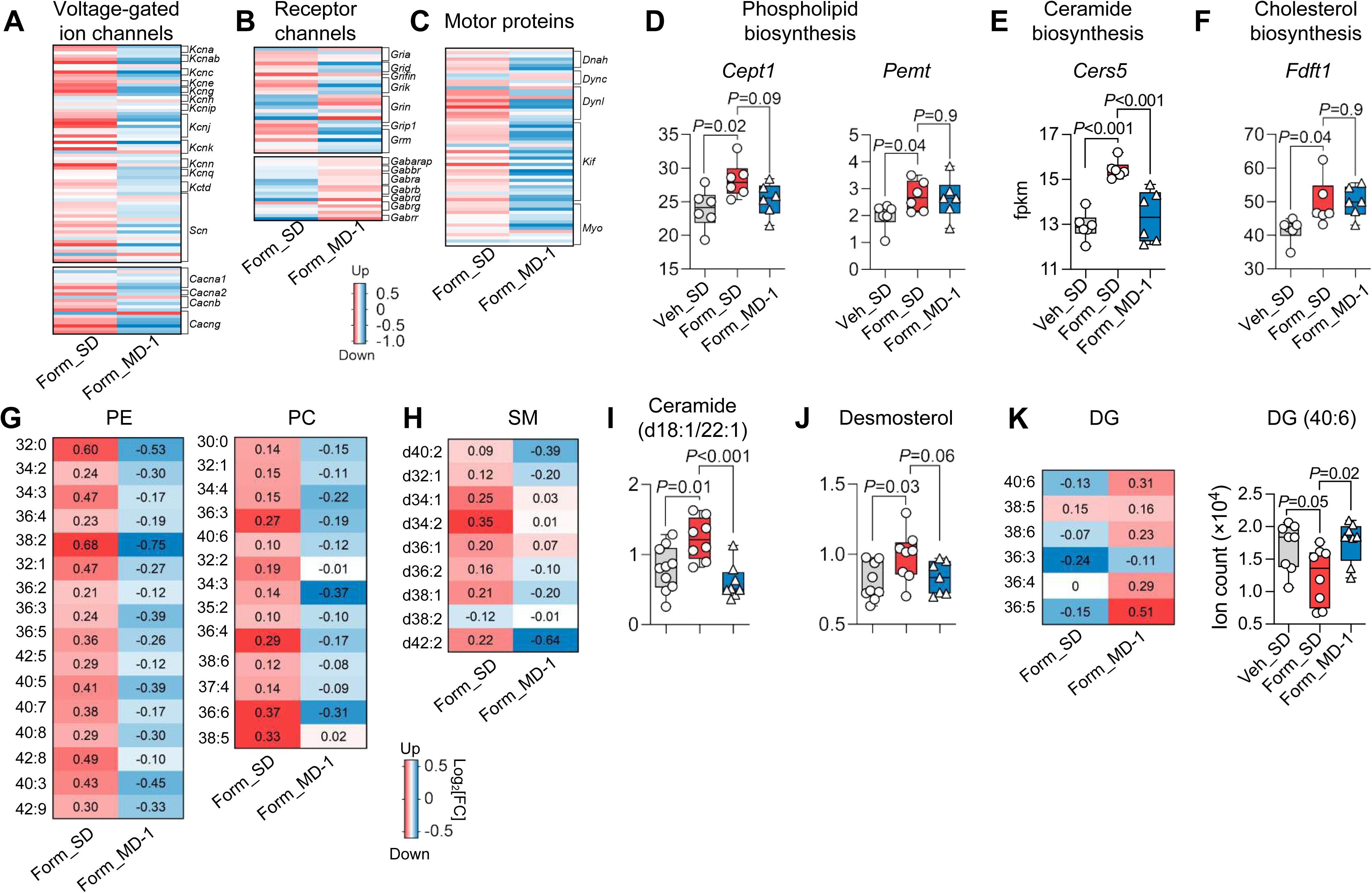
MD-1 prevents injury-induced biomass generation in the spinal cord. (A-C) Effects of formalin and MD-1 (log_2_ fold changes) on the transcription of genes related to (A) voltage-gated ion channels, (B) receptor channels, and (C) motor proteins. Form_SD: formalin *vs* vehicle injection in SD-1-fed mice; Form_MD-1: MD-1 *vs* SD feeding in formalin-injected mice. (D-F) Effects of formalin and MD-1 on the transcription (fpkm) of genes encoding biosynthetic enzymes for (D) glycerophospholipids (choline/ethanolamine phosphotransferase 1, *Cept1*), (E) ceramide (ceramide synthase 5, *Cers5*), and (F) cholesterol (squalene synthase, *Fdft1*). Veh_SD: vehicle-injected mice fed SD; Form_SD: formalin-injected mice fed SD; Form_MD-1: formalin-injected mice fed MD-1. (G, H) Effects of formalin and MD-1 (log_2_ fold changes) on the levels of (G) PE and PC, and (H) sphingomyelins (SM). Form_SD: formalin *vs* vehicle injection in SD-1-fed mice; Form_MD-1: MD-1 *vs* SD feeding in formalin-injected mice. (I, J) Effects of formalin and MD-1 on levels (ion count) of (I) ceramide (d18:1/22:1) and (J) desmosterol. Veh_SD: vehicle-injected mice fed SD; Form_SD: formalin-injected mice fed SD; Form_MD-1: formalin-injected mice fed MD-1. (K) Effects of formalin and MD-1 on DG levels (ion count). Form_SD: formalin *vs* vehicle injection in SD-1-fed mice; Form_ MD-1: MD-1 *vs* SD feeding in formalin-injected mice. (D, I-K) One-way ANOVA and Šídák’s test, *n* = 6-8 per group.

In addition to stimulating biomass production, formalin injection in SD-fed mice reduced the transcription of autophagy regulators unc-51-like autophagy-activating kinase 1 (*Ulk1*) and autophagy-related genes (*Atg*) 7, *Atg9a*, and *Atg2a* in ipsilateral lumbar hemicords, as shown by bulk RNAseq (Figure 4A). Protein levels of ATG5 and microtubule-associated protein 1 light chain 3b (LC3B) were also lower, compared to uninjured mice, as assessed by both immunoblot (Figure 4B and Supplemental Fig. S8) and immunofluorescent analyses, which also identified neurons as one of the cell types involved in this response (Figure 4C-D and Supplemental Fig. S9). MD-1 prevented the effects of injury, normalizing *Ulk1*, *Atg7*, *Atg9a*, and *Atg2a* transcription and restoring LC3B and ATG5 protein levels (Figure 4A-D and Supplemental Figs. S8 and S9). To determine whether autophagy suppression contributes to pain chronification, we administered the autophagy inhibitor 3-methyl adenine (TMA, 30 mg/kg, IP) to SD-fed mice treated with a formalin dose (0.1%, 20 mL) that is insufficient to induce chronic pain^26^ (Figure 4E). TMA administration on days 2-4 after low-dose formalin was followed by robust and sustained bilateral hypersensitivity (Figure 4F, G), indicating that autophagy inhibition enables minor injuries to cause enduring pain. In contrast, injection of low-dose formalin without TMA or saline plus TMA had no such effect (Figure 4F, G).

**Figure 4.**
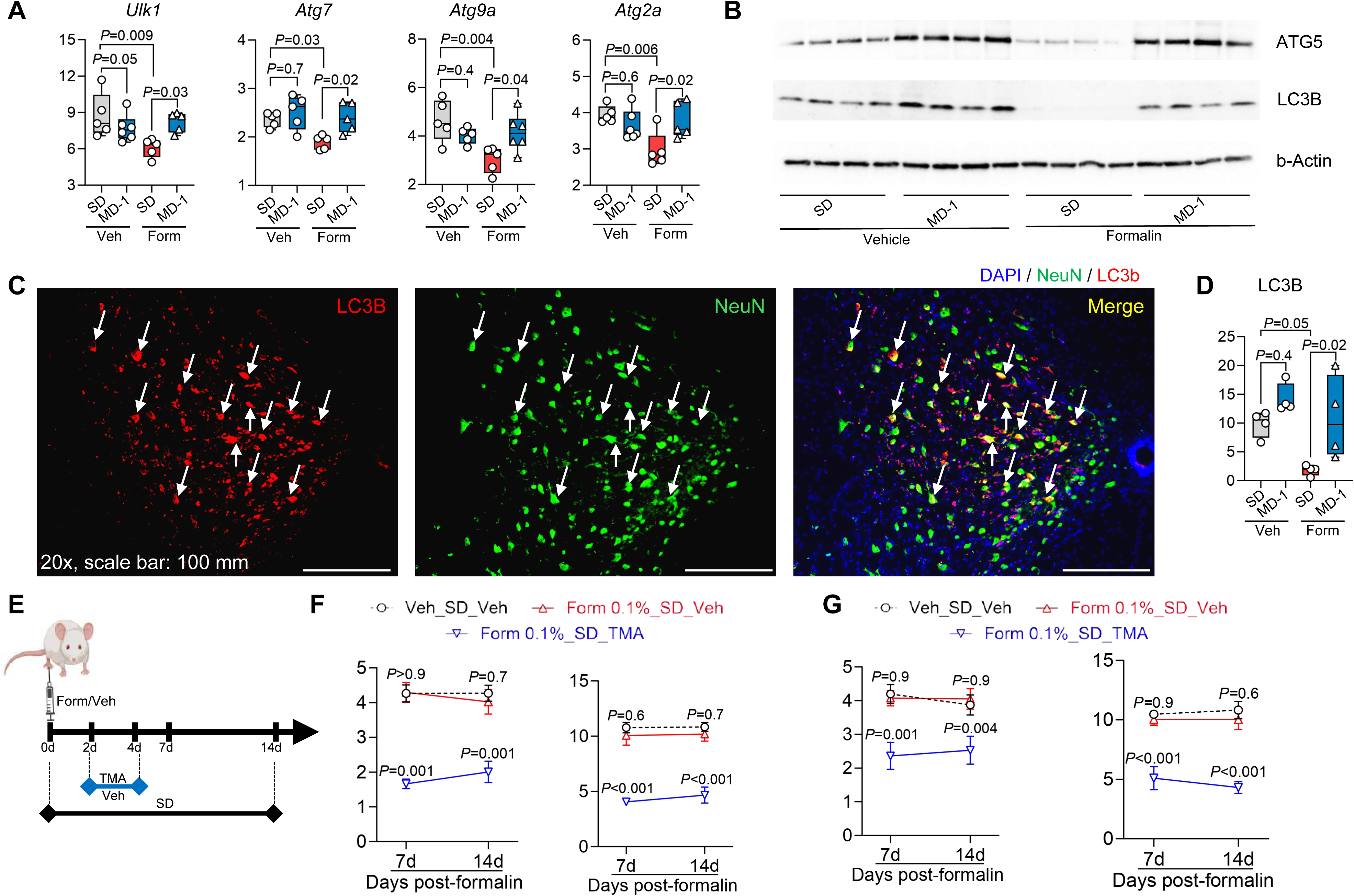
Injury-induced autophagy suppression in spinal cord facilitates the transition to chronic pain and MD-1 prevents autophagy suppression. (A) Transcription (fpkm) of key autophagy regulators in vehicle (Veh)- or formalin (Form)-injected mice fed SD or MD-1. (B) Representative Western blot images showing ATG5 and LC3B content in vehicle- and formalin-injected mice fed either SD or MD-1. β-actin is the loading control. (C) Representative immunofluorescent images for LC3B (red) and neuronal marker NeuN (green) in the L4-L6 spinal cord of formalin-injected mice fed MD-1. Nuclei are stained with DAPI. Arrows indicate LC3B and NeuN colocalization. Magnification: 20x. Scale bar: 100 μm. (D) Quantification of LC3B immunofluorescence in the L4-L6 spinal cord of vehicle (Veh)- or formalin (Form)-injected mice fed SD or MD-1. (E) Protocol: Autophagy inhibitor TMA (30 mg/kg, IP) or its vehicle (Veh) was administered to SD-fed mice on days 2-4 after formalin injection (F, G) Effects of TMA on (F) contralateral or (G) ipsilateral hypersensitivity to mechanical (left) and thermal (right) stimuli. Open circles: vehicle (veh)/SD/veh; red triangles: formalin/SD/veh; blue triangles: formalin/MD-1/veh; open squares, formalin/MD-1/TMA. (A, B, D, F, G) One- or two-way ANOVA followed by Dunnett or Šídák’s test. \**P*<0.05, \*\**P*<0.01, \*\*\**P*<0.001 compared to formalin-SD (*n* = 7-10).

Autophagy suppression promotes apoptosis^51^. Accordingly, formalin injections in SD-fed mice increased the transcription of proapoptotic genes (e.g., *Casp3*, *4*, *8*, and *12*) in L4-L6 spinal cord while concurrently downregulating survival genes such as *Bcl2* and *Mcl1*, compared to uninjured controls (Supplemental Fig. S10A). Immunofluorescent analyses revealed elevated levels of activated caspase-3 immunoreactivity in neurons and other cells of the lumbar spinal cord (Supplemental Fig. S10B). MD-1 did not influence the expression of apoptosis-related genes in uninjured mice, but effectively blocked the changes evoked by formalin (Supplemental Fig. S10A, B). To determine whether apoptosis contributes to pain chronification, we administered the pan-caspase inhibitor emricasan^52^ (3 mg/kg, IP) to SD-fed mice treated with 1% formalin. Emricasan administration on days 2-4 post-formalin normalized cleaved caspase-3 levels in the lumbar spinal cord (Supplemental Fig. S11), indicating that it effectively halted apoptosis. Notably, however, emricasan failed to prevent the transition to persistent hypersensitivity (Supplemental Fig. S12), indicating that apoptosis stimulation—unlike autophagy suppression—is not involved in pain chronification.

Collectively, these findings provide two key insights: (1) peripheral injury suppresses autophagy in spinal cord neurons, and this suppression plays a critical role in the progression to pain chronicity; and (2) autophagy suppression is accompanied by an increase in neuronal apoptosis, which, however, does not contribute to pain chronification.

### Injury alters the systemic metabolome and MD-1 prevents this effect

Spinal cord changes were accompanied by marked alterations in the circulating metabolome (Supplemental Fig. S13A and Supplemental Table S6). On day 4 after formalin injection, serum concentrations of corticosterone and aldosterone were significantly elevated in SD-fed mice, indicating sustained activation of the hypothalamic-pituitary-adrenal axis and renin-angiotensin system (Supplemental Fig. S13B). Additionally, levels of 3-methylglutarylcarnitine, a potential marker of mitochondrial dysfunction^53^, and taurine, a substance with anti-apoptotic properties^54^, were increased (Supplemental Fig. S13C). Serum accumulation of phosphatidylcholine (PC) and sphingomyelin (SM) mirrored similar alterations in the spinal cord (Supplemental Fig. S13D). Also consistent with spinal cord findings, various diacylglycerols and long-chain fatty acids—erucic, nervonic (24:1D^15^), and docosatrienoic (22:3D ^13,16,19^)—were lower in formalin-treated mice (Supplemental Fig. S13E). MD-1 countered these trends, normalizing serum corticosterone, aldosterone, 3-methylglutarylcarnitine, and taurine (Supplemental Fig. S13B, C), while resulting in lower PC and SM levels (Supplemental Fig. S13D), and replenishing diacylglycerols and fatty acids (Supplemental Fig. S13E). Importantly, the effects of MD-1 did not reflect modifications in the intestinal microbiome as by fecal samples analysis we observed that, compared to SD, a 25-day exposure to MD-1 did not influence the Shannon diversity index (Supplemental Fig. S14A) or the relative abundance of intestinal bacterial genera (Supplemental Fig. S14B, C). Thus, chronic pain development is associated with profound changes in the systemic metabolome, which partially reflect those in ipsilateral spinal hemicords but also likely result from other systemic sources, including dorsal root ganglia and other peripheral tissues. These changes are unaffected by gut microbiome composition and are prevented by MD-1.

### MD-1 prevents the transition to chronic pain following peripheral injury

The findings thus far suggest that Akt/mTORC1 activation in the spinal cord stimulates biomass production and autophagy suppression, promoting a localized nutrient deficit that facilitates chronic pain development. To further test this hypothesis, we examined whether MD-1 interrupts this transition. We injected formalin or saline into the hind paw of male SD- or MD-1-fed mice and monitored them for the subsequent three weeks, with access to MD-1 extended for one week after injection (Figure 5A). As expected^36^, in the SD-fed group formalin administration elicited a biphasic nocifensive response (Figure 5B) accompanied by paw inflammation (Supplemental Fig. S15A). This acute reaction was followed by an enduring (>3 months) pathological state that exhibited three hallmarks of severe chronic pain in humans^26^: bilateral hypersensitivity to mechanical and thermal stimuli (contralateral: Figure 5C, D; ipsilateral: Supplemental Fig. S15B)^37,38^, heightened anxiety-like behavior^39^ (Figure 5E and Supplemental Fig. S15C), and long-term memory deficits (Figure 5F)^40^. MD-1 reduced the second, but not the first, phase of the formalin response (Figure 5B), attenuated edema (Supplemental Fig. S15A), and, most crucially, stopped all behavioral signs of chronic pain (Figure 5C-F and Supplemental Fig. S15B, C). Comparable effects were seen in female mice (Supplemental Fig. S16A-E), though MD-1 did not normalize anxiety-like behavior in females (Supplemental Fig. S16F). Due to its slight caloric enrichment (SD = 3.2 kcal/g *vs* MD-1 = 3.7 kcal/g, Supplemental Table S1), male mice fed MD-1 showed a small but statistically detectable difference in body-weight gain compared to SD-fed controls (Supplemental Fig. S17). However, by covariance analysis we found that this difference did not affect pain outcomes (Supplemental Table S7). Additionally, corroborating our behavioral results, bulk RNA-seq analyses showed that the transcription of several proinflammatory genes—including interleukin 1β (*Il1b*), interleukin 6 (*Il6*), and tumor-necrosis factor-α (*Tnf*)^55,56^—was enhanced in ipsilateral lumbar hemicords of SD-fed formalin-treated mice, compared to vehicle-injected controls, but not in mice receiving MD-1 (Supplemental Fig. S18A). Expression of transforming growth factor-β3 (*Tgfb3*), which may have tissue-reparative functions^57^, was lower in injured mice, an effect also blocked by MD-1 (Supplemental Fig. S18B).

**Figure 5.**
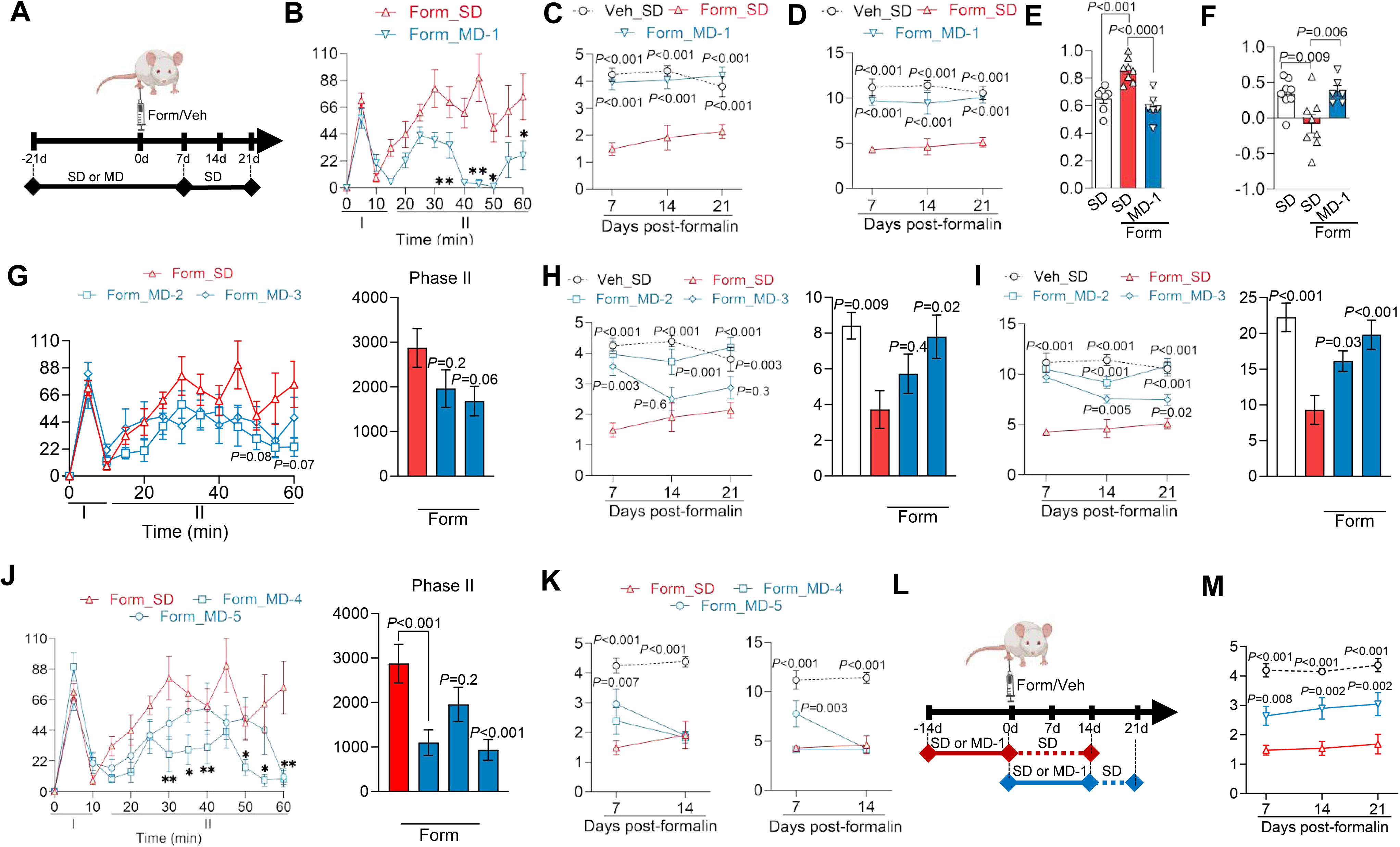
MD-1 prevents the progression to pain chronicity in a dose-, composition-, and time-dependent manner. (A) Mice received SD or a modified diet (MD-1, MD-2, MD-3, MD-4, or MD-5) for 21 days before formalin or saline injection. Access to the modified diets was extended for one week after injection and pain behaviors were monitored for the following 3 weeks. (B) Time-course (min) of the acute nocifensive response to formalin in mice fed SD (red triangles) or MD-1 (blue triangles). The response has two temporally distinct phases (I and II). (C, D) Time-course (days) of contralateral hypersensitivity to (C) mechanical and (D) thermal stimuli in formalin (Form)-injected mice fed SD (red triangles) or MD-1 (blue triangles). Open circles indicate vehicle (Veh)-injected mice fed SD. (E) Anxiety-like behavior (elevated plus-maze) in vehicle-injected mice fed SD (open bar) and formalin-injected mice fed SD (red bar) or MD-1 (blue bar). (F) Long-term memory (24-h novel-object recognition) in vehicle-injected mice fed SD (open bar) and formalin-injected mice fed SD (red bar) or MD-1 (blue bar). (G) Time-course (min) of the nocifensive response to formalin in mice fed SD (red triangles), MD-2 (half-dose MD-1, blue squares), or MD-3 (quarter-dose MD-1, blue diamonds). Right panel: quantification (area under the curve, AUC) for phase II of the formalin response in mice fed SD (red bar), MD-2, or MD-3 (blue bars). (H, I) Time-course (days) of contralateral hypersensitivity to (H) mechanical and (I) thermal stimuli in formalin-injected mice fed SD (red triangles), MD-2 (blue squares), or MD-3 (blue diamonds). Right panels: AUC quantification for the interval between 7 and 21 days. (J) Time-course (min) of the nocifensive response to formalin in mice fed SD (red triangles), MD-4 (blue squares), or MD-5 (blue circles). Right panel: AUC values for phase II of the formalin response in mice fed SD (red bar), MD-4, MD-5, or MD-1 (blue bars). (K) Time-course (days) of contralateral hypersensitivity to (left) mechanical and (right) thermal stimuli in formalin-injected mice fed SD (red triangles), MD-4 (blue squares), or MD-5 (blue circles). (L) Protocol for pre- and post-injury MD-1 administration. (M) Time-course (days) of contralateral mechanical hypersensitivity in formalin-injected mice fed SD (red triangles) or MD-1 (blue triangles) post-injury. Open circles indicate vehicle-injected mice fed SD. (B-K, M) One- or two-way ANOVA followed by Dunnett multiple comparisons test. \**P*<0.05, \*\**P*<0.01, \*\*\**P*<0.001 (formalin vs vehicle).

We next assessed whether dosage, chemical composition, or timing of administration influence MD-1’s efficacy. Firstly, we compared MD-1 with MD-2 and MD-3, two diets containing the same ingredients as MD-1 but at half and one-quarter the dosages, respectively (Supplemental Table S1). We fed mice SD, MD-1, MD-2, or MD-3 for 21 days followed by formalin or saline injection, with access to the diets extended for one week post-injection and then tracked pain-related behaviors over the following three weeks. Unlike MD-1, MD-2 and MD-3 had no effect on formalin-induced acute nociception or inflammation (Figure 5G and Supplemental Fig. S19, B). Additionally, MD-2 blocked the development of persistent hypersensitivity whereas MD-3 produced only transient and partial protection (contralateral: Figure 5H, I; ipsilateral: Supplemental Fig. S19B). The findings, summarized in Supplemental Table S8, suggest that MD-1 prevents chronic pain development in a dose-dependent manner, underscoring the specificity of its action.

We then investigated whether MD-1’s effects could be attributed to its nutrient content, the presence of PEA, or their combination. To address this, we tested two additional diets: MD-4, which matches MD-1 in nutrient composition and energy density but does not contain PEA, and MD-5, an SD supplemented exclusively with PEA (Supplemental Table S1). MD-4 attenuated the second phase of the acute nocifensive response to formalin (Figure 5J) but did not prevent edema formation or persistent hypersensitivity (contralateral: Figure 5K; ipsilateral: Supplemental Fig. S20A, B). In contrast, MD-5 did not affect formalin-induced nociception (Figure 5J) but reduced edema (Supplemental Fig. S18C) and delayed, without stopping, hypersensitivity development (contralateral: Figure 5K; ipsilateral: Supplemental Fig. S18B, C). Interestingly, although neither MD-4 nor MD-5 affected the emergence of hypersensitivity, both diets had distinct effects on the emotional and cognitive sequelae of injury: MD-4 improved memory but failed to alleviate anxiety-like behavior, whereas MD-5 reduced anxiety-like behavior but had only a borderline effect (P=0.05) on memory (Supplemental Fig. S20D, E). The results (Supplemental Table S8) suggest that MD-1’s combination of amino acids, fatty acids, and PEA is essential to provide comprehensive protection against pain chronification after injury.

Finally, we investigated whether the timing of MD-1 exposure—either prior to or following injury—would influence the diet’s efficacy. For this experiment, mice were either fed MD-1 for two weeks leading up to the day of formalin injection, at which point they were switched to an SD, or they began the MD-1 diet on the day of the injection, which was continued for 2 weeks before the mice were returned to SD feeding (Figure 5L). Post-formalin MD-1 exposure partially prevented hypersensitivity (contralateral mechanical: Figure 5M; other responses: Supplemental Fig. S21A-C) and reduced the memory deficit (Supplemental Fig. S21D) but had no effect on anxiety-like behavior (Supplemental Fig. S21E). On the other hand, limiting the mice to pre-formalin MD-1 exposure caused only a modest decrease in ipsilateral mechanical hypersensitivity (Supplemental Fig. S21F) without altering other pain-related responses (Supplemental Fig. S21G-K). The findings indicate that MD-1 administration in the weeks after—but not before— formalin injection provides some protection, reaffirming the centrality of this time window in chronic pain consolidation^26^. Full efficacy is observed, however, only when MD-1 is given both pre- and post-injury, suggesting that preemptive exposure increases systemic stores of depletion-prone nutrients.

### MD-1 prevents pain chronification by activating SIRT1 and AMPK and restoring autophagy

SIRT1 stimulates autophagy *via* both direct and indirect mechanisms^58^, prompting us to test whether this nutrient-sensing NAD^+^-dependent deacetylase^33,59^ could contribute to the protective effects of MD-1. By transcriptomic and immunoblot experiments we confirmed this possibility, finding that expression of SIRT1, but not other sirtuin family members, was elevated in ipsilateral lumbar hemicords of uninjured mice receiving MD-1, compared to SD-fed controls (Figure 6A-C and Supplemental Fig. S22A). Additionally, formalin administration markedly reduced SIRT1 protein levels, but not *Sirt1* transcription (Figure 6A-C), suggesting that injury may stimulate SIRT1 proteolysis^60^. In uninjured mice, MD-1 also enhanced transcription of the ketogenic enzyme, hydroxymethylglutaryl-CoA synthase-2 (*Hmgcs2*) (Figure 6D), which is indirectly regulated by SIRT1^61^. This transcriptional activation correlated with accrued b-hydroxybutyrate levels (Figure 6E). *Hmgcs2* transcription and ketone body production were not affected by formalin treatment (Figure 6D, E).

**Figure 6.**
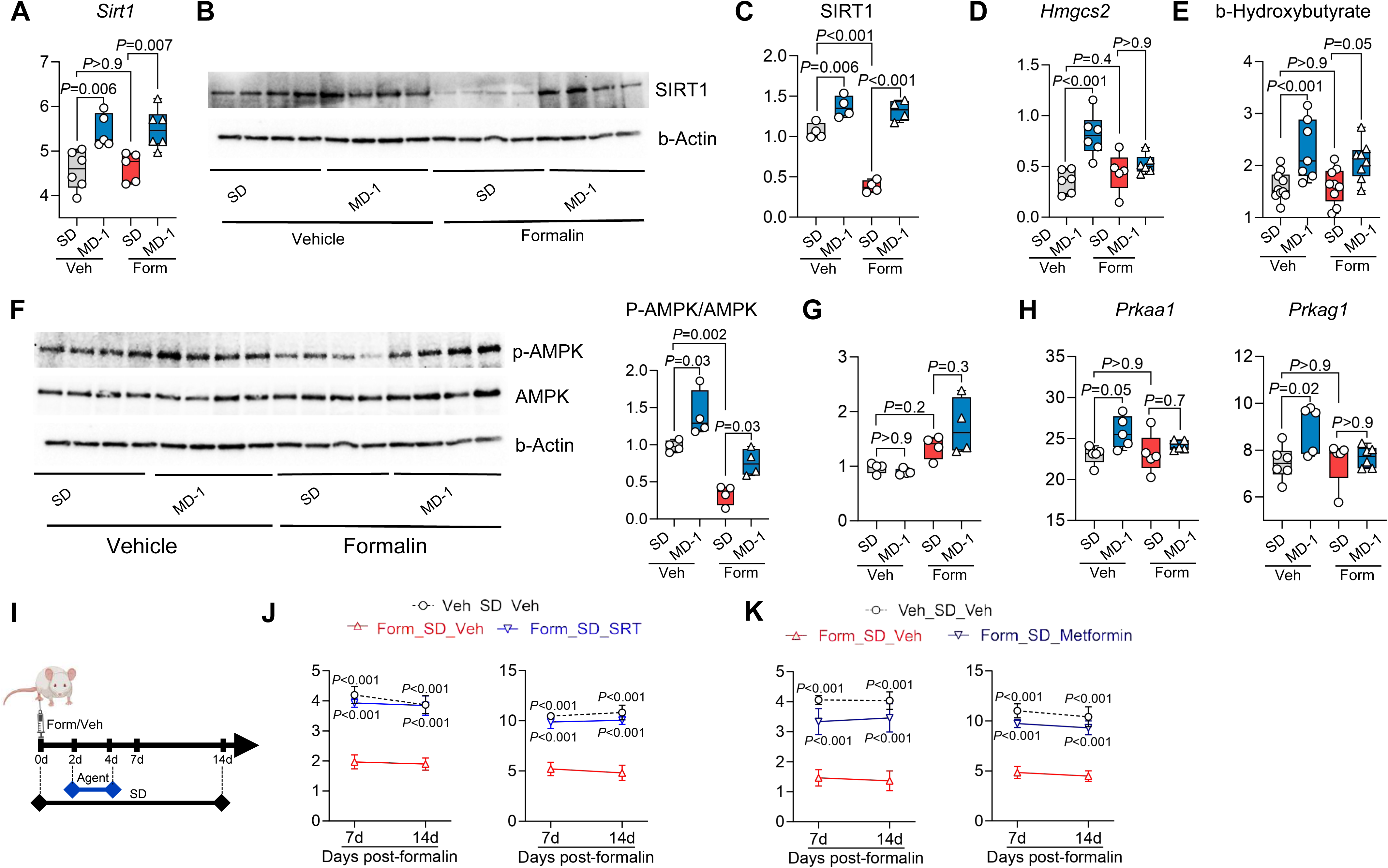
SIRT1 and AMPK mediate the protective effects of MD-1. (A) *Sirt1* gene transcription (fpkm) in the L4-L6 spinal cord of vehicle (Veh)- or formalin (Form)-injected mice fed SD or MD-1. (B) Representative Western blot images showing SIRT1 protein content in vehicle- or formalin-injected mice fed SD or MD-1. β-actin is the loading control. (C) SIRT1 protein quantification (SIRT1/β-actin) in vehicle- or formalin-injected mice fed SD or MD-1. (D) *Hmgcs2 t*ranscription in vehicle (Veh)- or formalin (Form)-injected mice fed SD or MD-1. (E) β-hydroxybutyrate levels in vehicle (Veh)- or formalin (Form)-injected mice fed SD or MD-1. (F) Representative Western blot images showing phosphorylated AMPK (p-AMPK) and total AMPK levels. Right: p-AMPK quantification (p-AMPK/AMPK). (G) AMPK protein quantification (AMPK/β-actin) in vehicle (Veh)- or formalin (Form)-injected mice fed SD or MD-1. (H) Transcription of AMPK subunits, *Prkaa1* and *Prkag1* in vehicle (Veh)- or formalin (Form)-injected mice fed SD or MD-1. (I) Protocol: SIRT1 activator SRT-2104 (100 mg/kg, IP), AMPK activator metformin (200 mg/kg, IP) or their vehicle were administered to SD-fed mice on days 2-4 after formalin injection. (J-K) Effects of (J) SRT-2104 and (K) metformin on contralateral hypersensitivity to mechanical (left) and thermal (right) stimuli. (A, C-E, G, H, J, K) One- or two-way ANOVA with Dunnett or Šídák’s multiple comparisons test. \**P* < 0.05, \*\**P* < 0.01, \*\*\**P* < 0.001 compared to formalin/SD (n = 4-8).

We examined whether AMPK, another sensor of nutrient availability^62^, also contributes to MD-1’s actions. Three observations supported this possibility (Figure 6F): 1) MD-1 increased AMPK phosphorylation (activation) in uninjured mice; 2) formalin injection decreased AMPK phosphorylation; and 3) MD-1 negated formalin’s effect. AMPK protein levels were not altered by either MD-1 or formalin injection (Figure 6G). The transcription of two AMPK subunits (*Prkaa1*, *Prkag1*) was stimulated by MD-1 but was not affected by injury (Figure 6H), while transcription of other subunits (*Prkaa2*, *Prkab2*) was reduced by injury and normalized by MD-1 (Supplemental Fig. S23).

These changes suggest that SIRT1 and AMPK play an important role in the protective effects of MD-1. To assess SIRT1’s function, we administered the SIRT1 activator SRT-2104 (100 mg/kg, IP) on days 2-4 after injection of 1% formalin (Figure 6I). SRT-2104 blocked the development of hypersensitivity (contralateral: Figure 6J; ipsilateral: Supplemental Fig. S24A, B), but failed to prevent paw edema (Supplemental Fig. S24C). Next, we examined the effects of the SIRT1 inhibitor EX-527 (20 or 50 mg/kg, IP) administered on days 2-4 after injection of 0.1% formalin, which does not cause persistent hypersensitivity^25^. Treatment with 50 mg/kg EX-527 was followed by robust and sustained bilateral hypersensitivity (Supplemental Fig. S24D-G), indicating that SIRT1 inhibition facilitates pain chronification. A weaker response was observed at the 20 mg/kg dose (Supplemental Fig. S24D-G).

To evaluate AMPK’s contribution to the protective effects of MD-1, we administered the AMPK activator metformin (200 mg/kg, IP) to SD-fed mice on days 2-4 after injection of 1% formalin (Figure 6I). Unlike SRT-2104, metformin prevented the development of contralateral but not ipsilateral hypersensitivity (contralateral: Figure 6K; ipsilateral: Supplemental Fig. S25A, B). Like SRT-2104, however, metformin did not affect formalin-induced paw edema (Supplemental Fig. S25C).

Given that both SIRT1 and AMPK promote autophagy^33,59,62^, we next examined whether inhibiting this process would negate the protective effects of MD-1. We treated SD- or MD-1-fed mice with either of two autophagy inhibitors—TMA (30 mg/kg, IP) or bafilomycin A1 (1 mg/kg, IP)—on days 2-4 post-formalin and tracked pain-related behaviors over the subsequent two weeks. Both inhibitors abolished MD-1’s effects, demonstrating their dependence on autophagy activation (Supplemental Fig. S26). We interpret these results as indicating that MD-1 prevents the transition to pain chronicity through a mechanism that involves the activation of SIRT1 and AMPK and the consequent restoration of autophagy.

### MD-1 protects mice from post-surgical acute and chronic pain

Thus far, we have utilized MD-1 to investigate the link between injury-induced metabolic alterations in the spinal cord and pain chronification. The diet’s marked impact on formalin-induced pain encouraged us to explore the generalizability of its effects and their potential clinical value. To this end, we assessed MD-1’s efficacy in four models of chronic pain: three involving damage to either nerve (CCI, SNI)^29,30^ or paw tissue (SPI)^31^, and one involving immune stimulation (CFA)^63^. We performed CCI or SNI in male SD- or MD-1-fed mice and monitored pain-related responses for the following 21 days, extending access to MD-1 for an additional week post-surgery (Figure 7A). In SD-fed mice, CCI and SNI caused lasting (>3 weeks) ipsilateral hypersensitivity, while MD-1-exposed mice were substantially protected (Figure 7B-E). Moreover, SNI mice exhibited long-term memory deficits, which were also prevented by MD-1 (Supplemental Fig. S27). Similarly, SD-fed mice subjected to SPI (Figure 7F) developed ipsilateral hypersensitivity, although of shorter duration (<2 weeks) than CCI or SNI (Figure 7G, H). This painful state was markedly attenuated by MD-1 exposure (Figure 7G, H). MD-1 also normalized the nocifensive response to prostaglandin E_2_ (PGE_2_) administered on post-surgical day 14 (Figure 7I), indicating that nociceptive priming^64^ was blocked. Interestingly, paw healing was accelerated in mice receiving MD-1 (Figure 7J), suggesting an overall strengthening of the animals’ resilience to tissue damage. In contrast, MD-1 did not influence CFA-induced pain (Figure 7K), producing only a small attenuation on day 7 after CFA administration (Figure 7L, M). To explore possible cellular substrates underpinning MD-1’s effects, we quantified by immunofluorescence glial fibrillary acidic protein (GFAP), ionized calcium-binding adapter molecule 1 (IBA-1), and SIRT1 in lumbar spinal cord of mice fed either SD or MD-1 for three weeks. These analyses revealed that exposure to MD-1 normalized GFAP immunoreactivity, which was enhanced by formalin (Supplemental Fig. S28), but produced only a modest, not statistically significant effect on IBA-1 immunoreactivity (Supplemental Fig. S29). Additionally, MD-1 increased overall immunoreactive SIRT-1 levels in the lumbar spinal cord of both formalin- and vehicle-treated mice (Supplemental Fig. S30A, B) and normalized them in neurons and other cells following formalin injection (Supplemental Fig. S30B, C). Thus, MD-1 prevents persistent painful states induced by somatic injury, but not by immune triggers, through a mechanism that involves spinal cord neurons and astrocytes.

**Figure 7.**
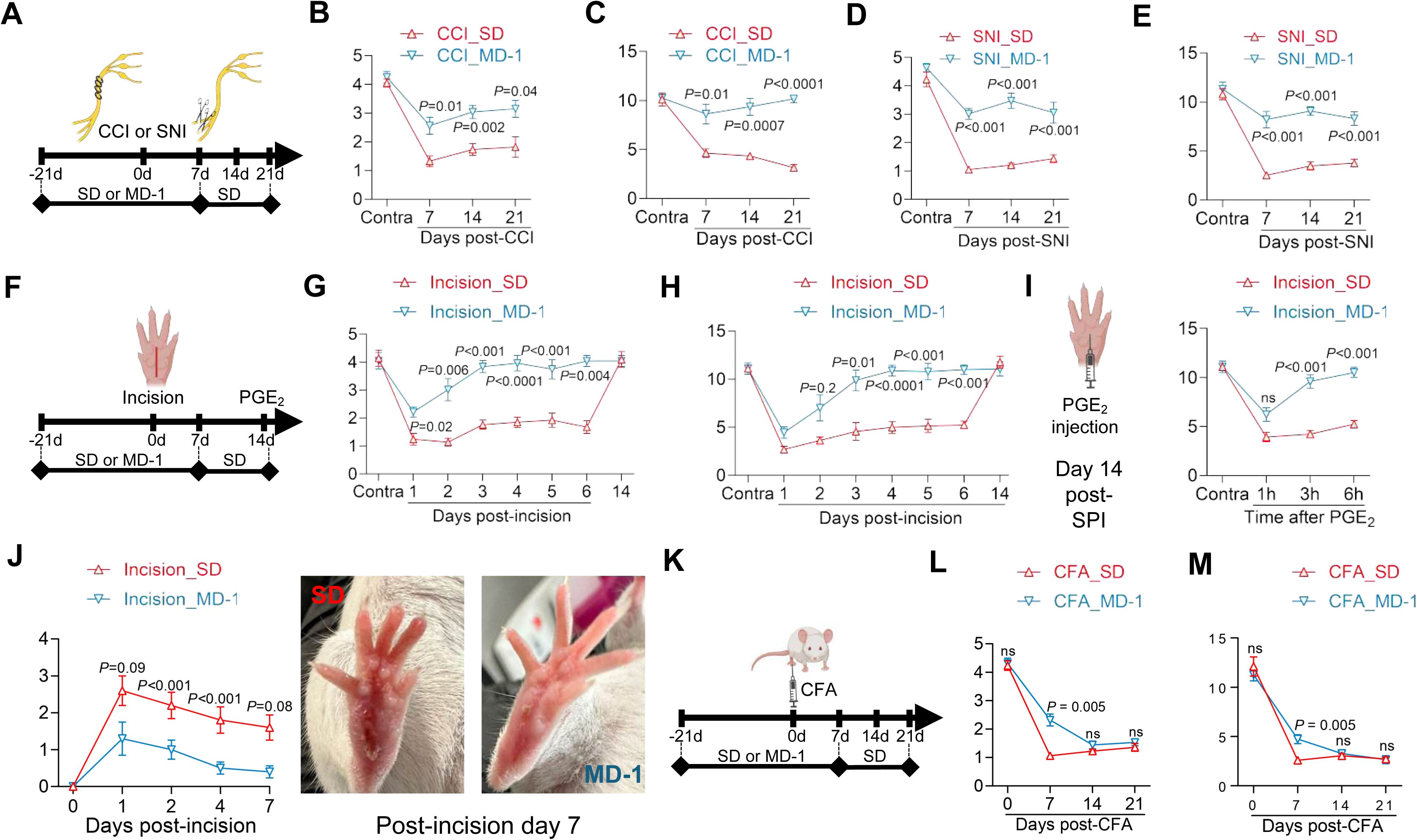
MD-1 prevents the development of acute and chronic pain after surgery. (A) CCI and SNI models: mice were fed SD or MD-1 for 21 days before surgeries. MD-1 access was extended for one week post-surgery and nocifensive behavior was monitored for the following 2 weeks. (B, C) Time-course of CCI-induced ipsilateral hypersensitivity (B, mechanical; C thermal) in mice fed SD (red triangles) or MD-1 (blue triangles). (D, E) Time-course of SNI-induced ipsilateral hypersensitivity (D, mechanical; E thermal) in mice fed SD (red triangles) or MD-1 (blue triangles). (F) SPI model: mice were fed SD or MD-1 for 21 days before SPI. MD-1 access was extended for one week after SPI and behavior was monitored for 2 more weeks. On day 14 post-SPI, the mice were given PGE_2_ (100 ng/20 mL, SC) in the lesioned paw and nocifensive behavior was monitored for 6 hours. (G, H) Time-course of SPI-induced ipsilateral hypersensitivity (G, mechanical; H thermal) in mice fed SD (red triangles) or MD-1 (blue triangles). (I) PGE_2_-induced ipsilateral thermal hypersensitivity in mice fed SD or MD-1. (J) Time-course (days) of first-intention wound healing after SPI in mice fed SD or MD-1. Right: representative images of incised paws from mice fed SD (left) and MD-1 (right). (K) CFA model: mice were fed SD or MD-1 for 21 days before hind-paw CFA injection. MD-1 access was extended for one week after injection and behavior was tracked for 3 weeks. (L, M) Time-course of CFA-induced ipsilateral hypersensitivity (L, mechanical; M thermal) in mice fed SD (red triangles) or MD-1 (blue triangles). (B-E, G-I, J, K-M) Two-way ANOVA with Dunnett’s multiple comparisons test (\**P* < 0.05, \*\**P* < 0.01, \*\*\**P* < 0.001 vs. SD group; n = 8-10 per group).

## Discussion

The results offer several novel insights into the molecular mechanisms underlying the progression to pain chronicity. First, we found that peripheral injury activates AKT/mTORC1 in afferent segments of the spinal cord, stimulating local biomass production, suppressing autophagy-mediated biomass reclamation, and depleting vulnerable pools of amino acids and fatty acids. This metabolic crisis is exacerbated by concomitant activation of the PEA-degrading enzyme N-acylethanolamine acid amidase^26,65^, which lowers PEA-mediated signaling at PPAR-α^66,67^, aggravating mitochondrial dysfunction^68^ and promoting neuroinflammation^69^. Second, experiments with the modified diet MD-1 reveal that this combined depletion of nutrients and PEA is not merely a byproduct of injury, but a driving force behind chronic pain. Third, we show that the nutrient sensor SIRT1 controls pain processing in the spinal cord and—along with AMPK, which was previously implicated in chronic pain^70,71^—mediates the protective effects of MD-1. Fourth, we show that autophagy is a critical regulator of spinal pain processing and that deficits in this process enable minor injuries to cause enduring pain. Lastly, we demonstrate that the stimulation of apoptosis by injury, which was previously reported^72^, is not directly involved in pain chronification. Collectively, these findings identify critical metabolic alterations in the spinal cord that drive the transition to pain chronicity and suggest pharmacological and nutritional strategies to halt this transition.

Previous work has shown that peripheral tissue damage induces central sensitization and persistent pain by recruiting AKT and its downstream target, mTORC1^23–25^. Our results outline the molecular cascade linking AKT/mTORC1 activation to chronic pain development (Supplemental Fig. S25). They demonstrate that AKT/mTORC1 signaling promotes the transcriptional upregulation of glycolysis-related genes, increasing the levels of glycolytic metabolites, which provide fuel for protein and lipid biosynthesis in support of neuroplasticity^17–21^. Concomitantly, SIRT1 and AMPK activities are downregulated, autophagy is suppressed, and critical amino acids, fatty acids, and PEA become depleted. Two key observations underscore the functional impact of this depletion: firstly, correcting the nutrient/PEA deficit with MD-1 stops the molecular cascade initiated by injury and interrupts pain chronification; secondly, inhibiting autophagy negates the protective effects of MD-1 and allows minor injuries to transition into lasting pain. These results emphasize the central role of core metabolism in chronic pain development, while raising several critical questions for future research, including: Which extracellular signals initiate injury-induced metabolic reprogramming? Which cell types release these signals, and which ones respond? And how does metabolic reprogramming reshape neuroglial structure in the spinal cord, to consolidate a painful state? Answering these questions will illuminate critical mechanistic aspects of the transition to pain chronicity and facilitate therapeutic discovery.

We designed MD-1 to augment the daily intake of specific amino acids, fatty acids, and PEA, which are depleted in the spinal cord following peripheral tissue damage. Pharmacokinetic studies show that MD-1 is systemically bioavailable and prevents this depletion. Replenishing the reduced nutrients normalizes energy metabolism and biomass generation, reinstates autophagy, and stops chronic pain development. Mechanistic investigations reveal three significant features of these effects. First, the effects exhibit dose- and time-dependence, indicating that MD-1 interferes with specific steps of the molecular cascade triggered by injury. Second, the effects rely on a synergistic interaction between MD-1’s constituents: neither an isocaloric diet enriched solely with amino acids and fatty acids (MD-4) nor one supplemented exclusively with PEA (MD-5) provides more than partial and transient protection. Interestingly, at doses significantly exceeding those in MD-1, PEA, like other PPAR-a agonists, can circumvent the need for nutrient supplementation, directly inhibiting pain chronification and underscoring the pivotal regulatory function of PPAR-a in this process^26^. Third, the effects of MD-1 involve multiple cell types in the spinal cord, including neurons and astrocytes, while microglia appear to play a minor role. Lastly, MD-1 enhances SIRT1 expression and AMPK activity – both of which are suppressed in the spinal cord of injured mice – while pharmacological activation of SIRT1 or AMPK replicates most, though not all, of MD-1’s effects. These findings highlight the integral role of SIRT1 and AMPK in modulating injury responses and mediating the therapeutic benefits of MD-1.

The best-understood function of SIRT1 and AMPK is to align core metabolism with energy availability^32,33,62,73^. Notably, the most distinctive effects of these nutrient sensors are reminiscent of those produced by MD-1. For example, in liver and skeletal muscle, SIRT1 engages PPAR-a and PGC1a, among other targets, to suppress glycolysis and lipid biosynthesis and promote fatty acid oxidation and ketogenesis^74–77^. Additionally, by interacting with mTORC1 and autophagosome component LC3B, SIRT1 represses global protein translation and enhances autophagy^78^. Similarly, AMPK downregulates anabolic processes and upregulates autophagy^62^. AMPK also forms a positive feedback loop with SIRT1 in which AMPK recruits SIRT1 by upregulating the NAD^+^-synthetic enzyme nicotinamide phosphoribosyltransferase^79^, while SIRT1 recruits AMPK by deacetylating its upstream kinase, liver kinase B1^80^. Our studies show that activation of SIRT1 and AMPK—achieved pharmacologically (using SRT2104 and metformin, respectively) or nutritionally (using MD-1)—effectively prevents pain chronification following injury. Interestingly, SIRT1 activation may also account for two effects of MD-1 that appear unrelated to the correction of injury-induced alterations in the spinal cord, namely, the diet’s ability to attenuate the acute nocifensive and inflammatory reaction to formalin and accelerate wound healing^81,82^. These findings highlight a system-wide impact of MD-1 and suggest its potential applicability to a broad range of diseases, including age-related pathologies where SIRT1 and AMPK downregulation is implicated^83–85^. Further studies are needed to explore these possibilities and clarify the precise mechanism through which SIRT1 and AMPK modulate the transition to chronic pain.

We conducted our mechanistic experiments in mice subjected to hind-paw formalin injection. We chose this model for three reasons. Firstly, while typically employed to study acute nociception^36^, it also produces a lasting phenotype that replicates key features of severe chronic pain in humans—such as contralateral sensitization and structural reorganization of the forebrain^26^— features not fully replicated by other models^29–31,63^. Secondly, the graded nature of the formalin response provides a valuable framework for investigating factors that either facilitate or repress pain chronification. Here, we leveraged this property to evaluate the roles of autophagy suppression and SIRT1 or AMPK activation. Thirdly, prior studies have identified a critical window for pain chronification in this model^26^, which we selected for our transcriptomic and metabolomic analyses. To evaluate the generalizability and translational relevance of MD-1’s effects, we tested the diet in four additional pain models: SNI, which involves partial sciatic nerve transection^29^; CCI, mimicking nerve compression and inflammation via sciatic nerve ligation^30^; SPI, inducing inflammation and sensitization through paw incision^31^; and CFA, characterized by immune-driven pain without direct nerve injury^63^. MD-1 demonstrated robust efficacy across all models except CFA, reinforcing the external validity of our findings and identifying post-surgical pain as a plausible clinical indication for this dietary intervention. The lack of effect in the CFA model confirms the existence of mechanistic differences between injury-induced and immune-driven pain^86^, warranting further investigation.

In conclusion, our findings indicate that injury induces, via AKT/mTORC1 activation, metabolic reprogramming, biomass production, and autophagy suppression. These processes converge to cause a metabolic crisis that depletes key nutrients and PEA in the spinal cord and drives the transition to chronic pain, most likely by disrupting normal neuroplasticity. MD-1’s efficacy in halting this transition underscores the critical role played by core metabolism in this process. By replenishing amino acids, fatty acids, and PEA, MD-1 enhances SIRT1 and AMPK activity, re-establishes autophagy, and prevents pain chronification across multiple mouse models. Thus, the results not only reveal novel mechanistic factors driving the progression to pain chronicity but also provide a basis for the clinical application of nutritional interventions targeting this progression after invasive surgeries^3,4^ or accidental physical trauma^5–7^.

### Limitations of the study

This study has two main limitations. Firstly, it was conducted in mice and its results cannot be directly extrapolated to humans due to marked differences in the way these species respond to metabolic challenges (*e.g.,* exercise)^87^. Nevertheless, the findings offer valuable insights to guide future clinical research: for example, MD-1’s efficacy in mouse models of post-surgical pain, combined with its lack of interference with wound healing, highlights its potential utility in perioperative analgesia and postsurgical pain prevention. Moreover, the partial overlap between metabolomic changes in mouse serum and molecular events in the spinal cord suggests a promising strategy for identifying serum biomarkers of chronic pain progression in humans. Secondly, the study primarily focused on the spinal cord, leaving other potential target organs unexamined. Future investigations should explore the involvement of additional sites, including first-order nociceptors, immune cells, and various metabolic organs, which may contribute to the systemic response to the bioenergy challenges posed by tissue damage.

## STAR ✪ Methods

### Resource availability

#### Lead contact

Further information and requests for resources and reagents should be directed to and will be fulfilled by the Lead Contact, Dr. Daniele Piomelli.

#### Materials availability

This study did not generate new unique reagents.

### Experimental models and subject details

#### Animals

We used male and female CD-1 mice (7 weeks of age upon arrival; Charles River, Wilmington, MA). The mice were housed in single-sex groups of 4-5 per cage under a 12-hour light/dark cycle (lights on from 06:30 to 18:30) at controlled temperature (20 ± 2°C) and relative humidity (55-60%). They had *ad libitum* access to food and water. Upon arrival, the mice were acclimated to the animal facility for one week and subsequently fed either a control standard diet (SD) or a modified diet (MD) for varying amounts of time. The study followed ethical guidelines for laboratory animal care set by the National Institutes of Health (NIH) and the International Association for the Study of Pain (IASP). Experimental procedures were approved by the Animal Care and Use Committee of the University of California, Irvine (AUP-23-082).

### Method details

#### Chemicals

We purchased isoleucine, valine, tyrosine, methionine, and complete Freund’s adjuvant from Sigma Aldrich (Saint Louis, MO). A proprietary water-soluble PEA formulation (Levagen^+®^) was a kind gift of Gencor Pacific (Austin, TX). Metformin, [^2^H_4_]-PEA, alanine, proline, serine, threonine, cysteine, tryptophan, phenylalanine, and leucine were from Cayman Chemicals (Ann Arbor, MI). Oleic acid and erucic acid were from Nu-Chek Prep (Elysian, MN). Torin-1, MK-2206, SRT-2104, 3-methyladenine, emricasan, EX-527 and bafilomycin A1 were from MedChemExpress (Monmouth Junction, NJ). All other chemicals were obtained from Sigma Aldrich (Saint Louis, MO) or Honeywell (Muskegon, MI, USA) and were of the highest available grade.

#### Modified diets

MD-1 and other experimental diets were formulated by coarsely grinding a standard mouse diet (SD; Envigo 2020x) using a commercial mixer. The resulting fine powder was enriched with laboratory-grade compounds specifically selected to address the depletion of key substances in the ipsilateral L4-L6 spinal cord observed on days 3–4 following hind-paw formalin injection (Supplemental Table S1). This enriched mixture was thoroughly blended with distilled water for 45 minutes to ensure uniformity. The prepared mixture was shaped into pellets using a pastry bag and extruded onto trays, followed by dehydration at room temperature for two days. The dried pellets were then stored in hermetically sealed containers, wrapped in tin foil to protect them from light and moisture. All diets were used within 30 days of preparation to ensure freshness and stability.

#### Drug administration

Drug solutions were freshly prepared prior to each use and administered via intraperitoneal (IP) or subcutaneous (SC) injection, depending on the experimental protocol. Metformin was dissolved in sterile saline. Torin-1, MK-2206, SRT-2104, emricasan, EX-527, TMA, and bafilomycin A1 were dissolved in a vehicle consisting of polyethylene glycol 400/Tween 80/saline (15:15:70, vol).

#### Pain models

##### Formalin

We injected a diluted formalin solution (1% or 0.1% in sterile saline, 20 µL) or saline into the plantar surface of the right hind paw, as described^88^. Following injections, the mice were immediately transferred to a transparent observation chamber where nocifensive behaviors (time spent licking or biting the injected paw and number of paw shakings) were videorecorded for 60 min to be later quantified by an observer blinded to experimental conditions. Mechanical hypersensitivity, heat hypersensitivity, and paw edema were measured at various time points in both formalin-injected and noninjected hind paws, as detailed under *Behavioral tests*.

##### Spared nerve injury (SNI)

We used a protocol described previously^89,90^. Briefly, the mice were anesthetized with 2–3% isoflurane in O_2_ delivered via a face mask. The right common sciatic nerve was exposed under aseptic conditions by blunt dissection at the level of its trifurcation into sural, tibial, and common peroneal nerves. The common peroneal and tibial branches of the sciatic nerve were tightly ligated with a 5.0 silk suture and transected distally, while the sural nerve was left intact. The wound was closed with a single muscle suture and skin clips. In sham-operated animals, the sciatic nerve was exposed but not transected.

##### Chronic constriction injury (CCI)

CCI of the sciatic nerve was carried out as described^88^. Briefly, the mice were anesthetized with 2–3% isoflurane in O_2_ delivered via a face mask. The right common sciatic nerve was exposed under aseptic conditions at the level of the middle thigh by blunt dissection. Proximal to the trifurcation, the nerve was cleaned from surrounding connective tissue, and three chromic catgut ligatures (4-0, Ethicon, Somerville, USA) were loosely tied around it at 1-mm intervals. The wound was closed with a single muscle suture and skin clips. In sham-operated animals, the sciatic nerve was exposed but not tied.

##### Surgical paw incision (SPI)

We performed SPI as described^91,92^. Briefly, the mice were anesthetized with 2–3% isoflurane in O_2_ delivered via a face mask. A 0.5-cm longitudinal incision was made under aseptic conditions through the skin and fascia of the plantar aspect of the right hind paw using a scalpel blade. The incision started 0.2 cm from the proximal edge of the heel and extended distally. The plantaris muscle was elevated with curved forceps and incised longitudinally, leaving the muscle origin and insertion intact. After hemostasis, the skin was sutured with a 6-0 nylon on a FS-2 needle (Ethicon, USA) and an antibiotic ointment was applied. Unwounded mice underwent a sham procedure consisting of anesthesia, antiseptic preparation, and ointment application. Following surgery, mice were returned to their home cages and continued receiving either SD or MD-1. Wound healing was monitored as described^93^, assigning a 1-point score to each of the following parameters: heat hypersensitivity, mechanical hypersensitivity, redness, visible edema, presence of pus, wound closure, and scar formation.

##### Hyperalgesic priming (HP)

HP was induced using the paw surgical incision protocol described above. Following surgery, mice were returned to their home cages. On day 14 post-surgical paw incision, prostaglandin E₂ (PGE₂; 100 ng in 20 µL) was administered subcutaneously into the lesioned paw. Nocifensive behavior (thermal hypersensitivity) was monitored for 6 hours post-injection.

##### Complete Freund’s adjuvant (CFA)-induced inflammation

We injected CFA (1 mg-mL^−1^, 20 µL) into the plantar surface of the right hind paw of slightly restrained mice^94^. Nocifensive behavior (thermal and mechanical hypersensitivity) was assessed before the injection and on days 7, 14 and 21 post-injection.

#### Behavioral tests

##### Mechanical hypersensitivity

Mechanical hypersensitivity was evaluated using a dynamic plantar aesthesiometer (Ugo Basile, Italy)^95^. After a 45-min habituation period in transparent cages positioned on a wire mesh surface, a mechanical stimulus was applied to the plantar surface of both hind paws by an automated steel filament exerting a force increasing from 0 to 5 g over 10 s. Withdrawal threshold was defined as the force (in grams) at which mice withdrew their paws from the mechanical stimulus. Three measurements were taken at intervals of 3 min and averaged.

##### Heat hypersensitivity

Sensitivity to heat was measured using a Hargreaves plantar test apparatus (San Diego Instruments, San Diego, USA) as described^95,96^. After a 45-min habituation period, the plantar surface of both hind paws was exposed to a beam of radiant heat through the glass floor. The cutoff time was set at 15 s. The stimulation was repeated three times with an interval of 2 min between stimuli, and latencies (in seconds) to withdraw the paw were recorded and averaged.

##### Tail-flick

Tail-flick assays were conducted following an established protocol^95^. Mice were gently restrained in a soft tissue pocket made of pet-training pad (Glad^TM^), and the distal 1/3 of each mouse’s tail was immersed in a hot water bath maintained at 54°C. The latency to withdraw the tail from the bath was recorded (in seconds). Measurements were performed twice, separated by a 5-min interval between trials, and the results were averaged. A 10-s cut-off time was implemented to prevent tissue damage.

##### Paw edema

Paw edema was measured with a digital caliper (Fisher Scientific, USA) and is expressed as the difference (Δ thickness, mm) between ipsilateral and contralateral paws.

##### Elevated plus maze

The test was performed under low ambient lighting (open arms: 160–180 lux and closed arms: 40–50 lux in accordance with an established protocol^97^. Briefly, each mouse was placed on the central platform of the maze, facing closed open arms, and the trial was recorded for 5 min using the Debut video capture software (NCH Software, Canberra, Australia). A blinded observer measured the time spent in the open and closed arms, as well as the number of entries into each arm type. The anxiety index was calculated using the formula: 1 - [(time spent in open arms/total time) + (open arm entries/total entries)] / 2^98^.

##### Novel object recognition

The test was conducted over 3 days^98,99^. On day 1, the mice were acclimated to the empty arena for 10 min. On day 2, they were reintroduced to the arena, which now contained two identical objects, and were left there for 10 min. On day 3, one of the objects was substituted with a new object of different shape, color, and texture. Mice were given another 10-min session to explore the arena, during which an observer blinded to experimental conditions recorded the total time spent exploring each object (i.e., nosing and sniffing at a distance ≤ 2 cm). The discrimination index was computed as: [(time of novel object exploration) – (time of familiar object exploration)]/total exploration time.

#### Tissue collection

Mice were deeply anesthetized with isoflurane. Blood was collected via cardiac puncture using syringes either rinsed with ethylenediaminetetraacetic acid (EDTA) or left unrinsed, and transferred into 1 mL polypropylene tubes containing either spray-coated potassium-EDTA (for plasma collection) or no anticoagulant (for serum collection). The blood was centrifuged at 1,450 × *g* for 15 min at 4°C. The resulting supernatant (serum or plasma) was carefully transferred into polypropylene tubes, immediately frozen, and stored at −80°C. Spinal cords were harvested by gentle hydraulic extrusion onto an ice-cold glass plate. The ipsilateral L4-L6 lumbar segments were dissected, snap-frozen on dry ice, and stored at −80°C until further analysis.

#### RNA sequencing and bioinformatics analysis

We extracted total RNA from L4-L6 spinal cord segments with the RNeasy Mini Kit (Qiagen) following manufacturer’s instructions. Samples with RNA integrity number (RIN) ≥ 8.5 were used for library construction. cDNA synthesis, amplification, library construction, and sequencing were performed at Novogene (Beijing, China) using the Illumina NovaSeq platform with paired-end 150–base pair sequencing strategy. Downstream bioinformatic analyses were performed using a combination of programs including STAR, HTseq, Cufflink and Novogene’s wrapped scripts, and alignments were parsed using STAR. Principal component analysis (PCA) and comparative analyses of differentially expressed genes (DEGs) were performed using the DESeq2/edgeR package and a model based on negative binomial distribution. Resulting *P* values were adjusted using the Benjamini and Hochberg’s approach for controlling false discovery rate (adjusted *P* values, *Padj*). Comparative analysis of DEGs was carried out between two test groups. Changes displaying *Padj* < 0.05 were considered significant. DEG distribution was assessed using Volcano plots showing statistical significance (*Padj*) vs magnitude of change (fold change). DEGs were annotated using the Database for Annotation, Visualization and Integrated Discovery (DAVID) database, PANTHER gene ontology (GO) knowledge base, and the Kyoto Encyclopedia of Genes and Genomes (KEGG) pathway database, which was implemented using the ShinyGO 0.80 bioinformatics platform. GO terms with *Padj* < 0.05 were considered significantly enriched in DEGs.

#### Metabolomic analyses

L4-L6 spinal cord segments were harvested and snap frozen on dry ice. Samples were pulverized to a homogeneous powder using a Cryomill (Retsch, Newtown, PA). An ice-cold mixture of methanol:acetonitrile:water (40:40:20, vol; 0.5-0.6 mL) was added to ∼10 mg of the powdered samples to make 25 mg/mL suspensions, which were centrifuged at 16,000 x *g* for 10 min at 4°C. For serum, 5 µL were diluted 30-fold with the same ice-cold mixture of methanol:acetonitrile:water and centrifuged under the same conditions. Supernatants (3 µL) from spinal cord and serum samples were analyzed as described ^100^. Briefly, a quadrupole-orbitrap mass spectrometer (Q Exactive Plus, ThermoFisher Scientific) operated in negative or positive ionization mode was coupled to a Vanquish Ultra High-Performance LC system (Thermo Fisher Scientific) with electrospray ionization. Scan range was *m/z* 70-1000, scanning frequency was 2 Hz and resolution was 140,000. LC separations were conducted using a XBridge BEH Amide column (2.1 mm x 150 mm^2^, 2.5 µm particle size, 130Å pore size pore size; Waters Corporation) with a gradient consisting of solvent A (20 mM ammonium acetate, 20 mM ammonium hydroxide in 95:5 water:acetonitrile, pH 9.45) and solvent B (acetonitrile). The flow rate was 0.150 mL/min. The gradient was: 0 min,85% B; 2 min, 85% B; 3 min, 80% B; 5 min, 80% B; 6 min, 75% B; 7 min, 75% B; 8 min, 70% B; 9 min, 70% B; 10 min, 50% B; 12 min, 50% B; 13 min, 25% B; 16 min, 25% B; 18 min, 0% B; 23 min, 0% B; 24 min, 85% B; 30 min, 85% B. Autosampler temperature was 5 °C. Data were analyzed using the MAVEN software (Build # 682), Compound Discoverer software (Thermofisher Scientific), and R software. To control for instrument variability, an internal control [^13^C_5_, ^15^N]-valine, was spiked in the extraction solvent.

#### PEA analysis

##### PEA extraction

We extracted PEA from plasma and spinal cord samples as described^101^. Briefly, plasma (0.1 mL) was transferred into 8-mL glass vials. Proteins were precipitated by the addition of 0.45 mL ice-cold acetonitrile containing 1% formic acid and 0.05 mL internal standard [^2^H_4_]-PEA. The mixture was stirred vigorously for 30 s and centrifuged at 1450 x *g* at 4°C for 15 min. Spinal tissue samples (∼20 mg each) were transferred into 2 mL Precellys CK-14 soft tissue tubes (Bertin, Rockville, MD) and homogenized in 0.5 mL of ice-cold acetonitrile containing 1% formic acid and the internal standard listed above. Supernatants from plasma or spinal tissue samples were loaded onto Enhanced Matrix Removal (EMR)-Lipid cartridges (Agilent Technologies, Santa Clara, CA) and eluted under positive pressure (3-5 mmHg, 1 drop/5 sec). Residual pellets from plasma and spinal tissue were rinsed with water/acetonitrile (1:4 vol/vol; 0.2 mL), stirred for 30 s, and centrifuged at 1450 x *g* at 4°C for 15 min. The supernatants were transferred onto EMR cartridges, eluted (1 drop/sec), and pooled with the first eluate. The cartridges were rinsed with water/acetonitrile (1:4 vol/vol; 0.2 mL), and pressure was gradually increased to 10 mmHg for maximal analyte recovery. Eluates were dried under a gentle stream of N_2_ (2 mmHg for 1 hour), reconstituted in methanol (0.1 mL) and transferred to deactivated glass inserts (0.2 mL) placed inside amber glass vials (2 mL, Agilent Technologies) for liquid chromatography/tandem mass spectrometry (LC-MS/MS) analyses.

##### PEA quantification

We quantified PEA using a 1260 series LC system (Agilent Technologies, Santa Clara, CA) coupled to a 6460C triple-quadrupole mass spectrometry detector (MSD; Agilent). An Eclipse PAH column (1.8 µm, 2.1 × 50 mm; Agilent Technologies) was eluted with a mobile phase consisting of 0.1% formic acid in water as solvent A and 0.1% formic acid in methanol as solvent B. A linear gradient was used: 75.0% B at 0 time to 80.0% B in 5.0 min, change to 95% B at 5.01 min continuing to 6.0 min, change to 75% B at 6.01 min, and hold till 8.0 min for column re-equilibration and stop time. The column temperature was maintained at 45°C and the autosampler temperature at 10°C. The injection volume was 2 μL, the flow rate was 0.3 mL/min, and the total analysis time was 15.5 min. To prevent carryover, the injection needle was washed three times in the autosampler port for 30 s before each injection, using a wash solution consisting of 10% acetone in water/methanol/isopropanol/acetonitrile (1:1:1:1, vol). The MSD was operated in the positive electrospray ionization (ESI) mode. PEA was quantified by multiple reaction monitoring (MRM) of the following transitions: PEA = *m/z* 300.3 > 62.2, [^2^H_4_]-PEA = *m/z* 304.3 > 66.2. The lowest limit of detection (LOD) was 0.5 ng/mL; lowest limit of quantification (LOQ) was 1 ng/mL. Capillary and nozzle voltages were 3,000 and 1,900 V, respectively. The drying gas temperature was 300°C with a flow of 5 mL/min. The sheath gas temperature was 300°C with a flow of 12 mL/h. Nebulizer pressure was set at 40 psi. We used MassHunter software version B.08.00 (Agilent Technologies) for instrument control, data acquisition, and analysis.

#### Protein analyses

##### Western blot

Western blot analyses were performed as described^102^. L4–L6 lumbar spinal cord segments were dissected, snap-frozen in liquid nitrogen, and stored at −80°C. For protein extraction, tissues were homogenized on ice in RIPA lysis buffer (Thermo Fisher Scientific, Waltham, MA) containing a protease and phosphatase inhibitor cocktail (Thermo Fisher Scientific). Homogenates were centrifuged at 13,000 × g for 10 min at 4°C, and the supernatants were collected as total protein extracts. Protein concentrations were determined using the Pierce BCA Protein Assay Kit (Thermo Fisher Scientific). Equal amounts of protein (20–40 µg) were mixed with 4× Laemmli sample buffer (Bio-Rad, Hercules, CA) containing β-mercaptoethanol, heated at 95°C for 5 min, and separated on 4%-12% SDS-PAGE gels. Proteins were transferred to nitrocellulose membranes at 100 V for 90 min in transfer buffer containing 20% methanol. Membranes were blocked in 5% skim milk in Tris-buffered saline with 0.1% Tween-20 (TBST, pH 7.4) for 1 hour at room temperature. Membranes were then incubated overnight at 4°C with primary antibodies listed above. Following incubation with primary antibodies, membranes were washed three times with TBST and incubated for 1 hour at room temperature with species-specific HRP-conjugated secondary antibodies (1:5000, Cell Signaling Technology). Membranes were washed again with TBST and developed using an ECL detection kit (Thermo Fisher Scientific). Blots were visualized using Image Lab 6.1 software, and densitometric analysis of band intensities was performed using ImageJ software (National Institutes of Health). Protein expression levels were normalized to β-actin and expressed as fold changes relative to control.

##### Immunofluorescence

Mice were deeply anesthetized and transcardially perfused with ice-cold phosphate-buffered saline (PBS), followed by 4% paraformaldehyde (PFA) in PBS. Lumbar spinal cords (L4–L6 segments) were dissected, post-fixed in 4% PFA for 4–6 hours at 4°C, and cryoprotected overnight in 30% sucrose in PBS (pH 7.4) at 4°C. Tissues were embedded in optimal cutting temperature (OCT) compound (Tissue-Tek®, Sakura Finetek, Torrance, CA) and flash-frozen in cold isopentane. Cryosections (10 µm thickness) were prepared using a cryostat and mounted on Superfrost Plus slides (Cat. #1255015, Thermo Fisher Scientific). Sections were rinsed with 0.1 M phosphate buffer (PB) (76.43 mM Na₂HPO₄, 23.73 mM NaH₂PO₄, distilled water, pH 7.4), permeabilized with 0.3% Triton X-100 in PB, and blocked with 3% normal horse serum (NHS) in PB for 1 hour at room temperature. Immunostaining was performed by incubating sections overnight at 4°C with primary antibodies diluted in staining buffer containing 3% NHS and 0.3% Triton X-100 in 0.1 M PB as follows: rabbit anti-LC3B (1:1000), rabbit anti-Caspase-3 (1:1000), rabbit anti-SIRT1 (1:1000), mouse anti-GFAP (1:400), mouse anti-IBA1 (1:1000), mouse anti-NeuN (1:400). After three 10-minute washes in 0.1 M PB, sections were incubated at room temperature for 1 hour in the dark with Alexa Fluor 568-conjugated goat anti-rabbit secondary antibody (1:1000) diluted in PB containing 3% NHS and 0.3% Triton X-100. Sections were mounted using VECTASHIELD® Antifade Mounting Medium with DAPI (Cat. #H1200, Vector Laboratories, Newark, CA) and sealed with coverslips. Images were captured at 10x, 20x, or 40x magnification using a Keyence BZ-X810 fluorescence microscope. LC3B-, SIRT1-, GFAP-, IBA1- or Caspase-3-positive puncta were analyzed using ImageJ software, and neuronal localization was confirmed by co-staining with mouse anti-NeuN antibody (1:400). Quantification of LC3B- and Caspase-3-positive puncta was performed in at least three randomly selected sections.

#### Gut microbiome analysis

##### Fecal sample preparation

Mixed fecal droppings (∼2ml in volume) were collected from 4 mice in each cage (*n* = 4 cages per group). The samples were placed into 15-mL tubes and immediately stored at -80°C. They were thawed once for DNA extraction and 16S rRNA sequencing.

##### Preparation of DNA and 16S library construction

Extraction of DNA from frozen stool samples was performed using Zymo Research Quick-DNA Fecal/Soil Microbe 96 Magbead Kit according to manufacturer’s instructions. Approximately 180-200 mg of stool sample was used for the DNA extraction. The resulting DNA was measured by Qubit and 5 ng was used as input for library construction. The library preparation was performed according to the Illumina 16S Metagenomic Sequencing Library Preparation protocol. More specifically, the protocol is a two-step PCR that begins with the primer pair sequences for the V3 and V4 region with partial Illumina adapter handles to generate a single amplicon of approximately ∼460 bp [16S Amplicon Forward (V3 region): 5’-TCGTCGGCAGCGTCAGATGTGTATAAGAGACAGCCTACG GGNGGCWGCAG-3, and 16S Amplicon Reverse (V4 region): 5’-GTCTCGTGGGCTCGGAGATGTGTATAAGAGACAGGACTACHVGGGTATCTAA TCC-3’]. The second step PCR completes the Illumina adapter and adds P5 and P7 indexes using primers for from the dual index kit for Nextera XT library construction. The resulting libraries were assayed for quantity using Qubit and for quality using the Agilent Bioanalyzer 2100 DNA HS chip. The libraries were normalized and then multiplexed together. The multiplexed library pool was quantified using qPCR and sequenced on Illumina Miseq 2X300bp run.

##### Analysis

We imported 978,449 demultiplexed Illumina Miseq sequence reads into QIIME2 version 2022.2 ^103^ (https://qiime2.org). Quality checking, denoising, and merging of paired-end reads were performed using DADA2 via the q2-dada2 plugin^104^. We picked operational taxonomic units (OTUs) at a 100% identity level (amplicon sequence variants) using UCLUST via QIIME2. We assigned taxonomy to the OTUs using the q2-feature-classifier, classify-sklearn naïve Bayes taxonomy classifier against the SILVA database 138 against the OTUs reference sequences^105,106^. The QIIME2-created OTU table as well as the taxonomy table and metadata were transferred into R for statistical analysis (R version 4.2.2). We rarefied the OTU table via randomized sampling without replacement with 300 iterations at 22,000 sequences per sample using the “EcolUtils” package (R core Team, 2018, https://www.r-project.org/; Salazar, G. 2020. EcolUtils: Utilities for community ecology analysis. https://github.com/GuillemSalazar/EcolUtils). We determined the effect of MD-1 on microbial composition was determined using Permutational multivariate analysis of variance (PERMANOVA) on a Bray Curtis dissimilarity matrix generated from the rarefied OTU table using the adonis2 function of the vegan package (version 2.6-4) in R. We performed a Shapiro-Wilk test to check for normality distribution of residuals for the Shannon diversity. Since the distribution was normal, an ANOVA was used to check for significance of any of the factors for alpha diversity.

### Statistical analyses

Statistical analyses were conducted using GraphPad Prism version 10.2 (La Jolla, CA). Results are presented as means ± SEM of n experiments. Statistical significance was set at *P* < 0.05 and assessed using unpaired, two-tailed Student’s *t* test or analysis of variance (ANOVA) (one-way or two-way) followed by post hoc tests, as appropriate. Analysis of transcriptomics and microbiome data were conducted as described in previous sections.

## Acknowledgments

This work was supported by the National Institute on Drug Abuse (DA055578, to D.P.). Additional funding was provided by grant AT012658 (to Y.F.), Pew Foundation, R01-AA029124 and R21-AA030358 (to CJ), and F31DK134173 (to JL). The authors thank the researchers of the Genomics High-Throughput Facility (GHTF) and Microbiome Center in the University of California, Irvine, for assistance with microbiome analysis; and Dr. Randy Levinson for critical reading of the manuscript.

## Author contributions

D.P. and A.M.T. designed the studies; A.M.T, Y.F, H.L., J.K., J.L. performed the experiments with assistance from K.M.J, C.J. and F.A. D.P. ideated the project, supervised it, and wrote the manuscript with assistance from all other authors.

## Data and code availability

- Transcriptomic, and metabolomic data have been deposited in DRYAD and are publicly available as of the date of publication. Accession numbers are listed in the <u>key resources table</u>. <u>Data S1</u> refers to unprocessed data underlying the display items in the manuscript, related to all main and supplemental figures.
- This paper does not report original code.
- Any additional information required to reanalyze the data reported in this paper is available from the lead contact upon request.

## Key resources table

| REAGENT or RESOURCE | SOURCE | IDENTIFIER |
| --- | --- | --- |
| **Antibodies** |  |  |
| Rabbit polyclonal anti-p-AKT (Ser473) | Cell Signaling Technology | 9271 |
| Rabbit polyclonal anti-AKT | Cell Signaling Technology | 9272 |
| Rabbit monoclonal anti-p-mTOR (Ser2448) | Cell Signaling Technology | 5536 |
| Rabbit polyclonal anti-mTOR | Cell Signaling Technology | 2972 |
| Rabbit monoclonal anti-p-AMPKa (Thr172) | Cell Signaling Technology | 2535 |
| Rabbit polyclonal anti-AMPKa | Cell Signaling Technology | 5831 |
| Rabbit polyclonal anti-LC3B | Cell Signaling Technology | 2775 |
| Rabbit polyclonal anti-Caspase-3 (Asp175) | Cell Signaling Technology | 9661 |
| Mouse anti-Iba1/AIF1 | Millipore Sigma | MABN92 |
| Rabbit monoclonal anti-SIRT1 (D1D7) | Cell Signaling Technology | 9475 |
| Mouse monoclonal anti-GFAP (GA5) | Cell Signaling Technology | 3670 |
| Mouse anti-NeuN (E4M5P) | Cell Signaling Technology | 94403 |
| Alexa Fluor 568-conjugated donkey anti-rabbit | Invitrogen | A78946 |
| Alexa Fluor 488-conjugated goat anti-mouse | Invitrogen | A11029 |
| Mouse monoclonal anti-SIRT1 | Abcam | ab110304 |
| **Chemicals, peptides, and recombinant proteins** |  |  |
| Isoflurane | Pivotal veterinary | 21295098 |
| EDTA (0.5M), pH 8.0 | Thermo Fisher Scientific | AM9260G |
| Acetonitrile | Thermo Fisher Scientific | T001014000 |
| Methanol | Thermo Fisher Scientific | T001024000 |
| Water, HPLC grade | Thermo Fisher Scientific | WSSK-4 |
| Formic acid | Thermo Fisher Scientific | A117-50 |
| Invitrogen TRIzol | Thermo Fisher Scientific | 15596026 |
| Isoleucine | Sigma Aldrich | I2752 |
| Valine | Sigma Aldrich | V0500 |
| Tyrosine | Sigma Aldrich | T3754 |
| Methionine | Sigma Aldrich | M9625 |
| Complete Freund's Adjuvant (CFA) | Sigma Aldrich | F5881 |
| PEA formulation (Levagen<sup>®</sup>+) ) | Gencor Pacific | PML0010815 |
| Metformin | Cayman Chemicals | 13118 |
| Palmitoylethanolamide-d4 ([<sup>2</sup>H<sub>4</sub>]-PEA) | Cayman Chemicals | 10007824 |
| Alanine | Cayman Chemicals | 29757 |
| Proline | Cayman Chemicals | 30772 |
| Serine | Cayman Chemicals | 33561 |
| Threonine | Cayman Chemicals | 30194 |
| Cysteine | Cayman Chemicals | 33309 |
| Tryptophan | Cayman Chemicals | 29600 |
| Phenylalanine | Cayman Chemicals | 31498 |
| Leucine | Cayman Chemicals | 34342 |
| Oleic acid | Nu-Chek Prep | U-46-A |
| Erucic acid | Nu-Chek Prep | U-79-A |
| Torin-1 | MedChemExpress | HY-13003 |
| MK-2206 | MedChemExpress | HY-10358 |
| SRT-2104 | MedChemExpress | HY-15262 |
| 3-methyladenine (TMA) | MedChemExpress | HY-19312 |
| Bafilomycin A1 (bafA1) | MedChemExpress | HY-100558 |
| Emricasan | MedChemExpress | HY-10396 |
| EX-527 | MedChemExpress | HY-15452 |
| Formic acid | Thermo Fisher Scientific | 28905 |
| Antifade Mounting Medium with DAPI | Vector Laboratories | H1200 |
| Protease inhibitor | Sigma-Aldrich | P8340 |
| **Critical Commercial Assays** |  |  |
| Quick-DNA Fecal/Soil Microbe 96 Magbead Kit | Zymo Research | D6012-FM |
| Pierce BCA Protein Assay Kit | Thermo Fisher Scientific | 23227 |
| RNeasy Mini Kit | Qiagen | 74104 |
| **Deposited data** |  |  |
| Transcriptomic and Metabolomic dataset | Deposited to DRYAD | https://doi.org/10.7286/1KD8R |
| **Experimental models: Organisms/strains** |  |  |
| Mouse: *CD-1* |  |  |
| **Software and algorithms** |  |  |
| ImageJ | https://imagej.nih.gov/ij/download | N/A |
|  | ml |  |
| GraphPad Prism | GraphPad Prism | Version 10.2.3 |
| MassHunter | Agilent Technologies | N/A |
| STRING | https://string-db.org/ | N/A |

### Supplemental Figures and Tables

**Supplemental Fig. S1.**
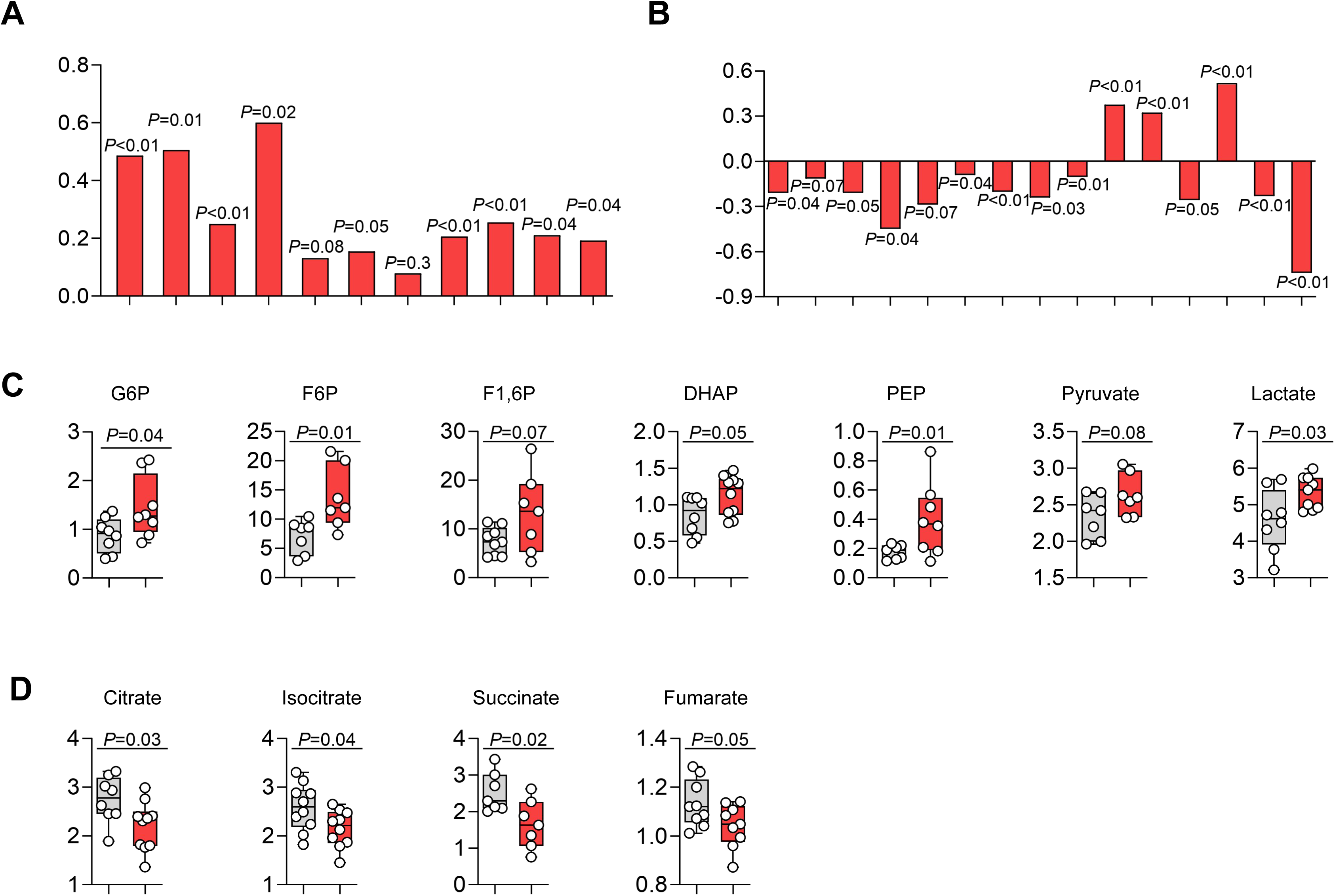
(Related to Figure 1) Effects of formalin injection on gene transcription and metabolite levels in ipsilateral L4-L6 spinal cord of SD-fed mice. (A, B) Transcription of genes involved in (A) glycolysis and (B) Krebs’s cycle and oxidative phosphorylation. Data are expressed as log_2_ changes (formalin vs vehicle) (*n* = 6; multiple unpaired *t* test). (C, D) Concentrations of (C) glycolysis and (D) Krebs’s cycle metabolites. Data are expressed as ion counts (mean ± SEM; *n* = 7–10 per group; Student’s *t* test).

**Supplemental Fig. S2.**
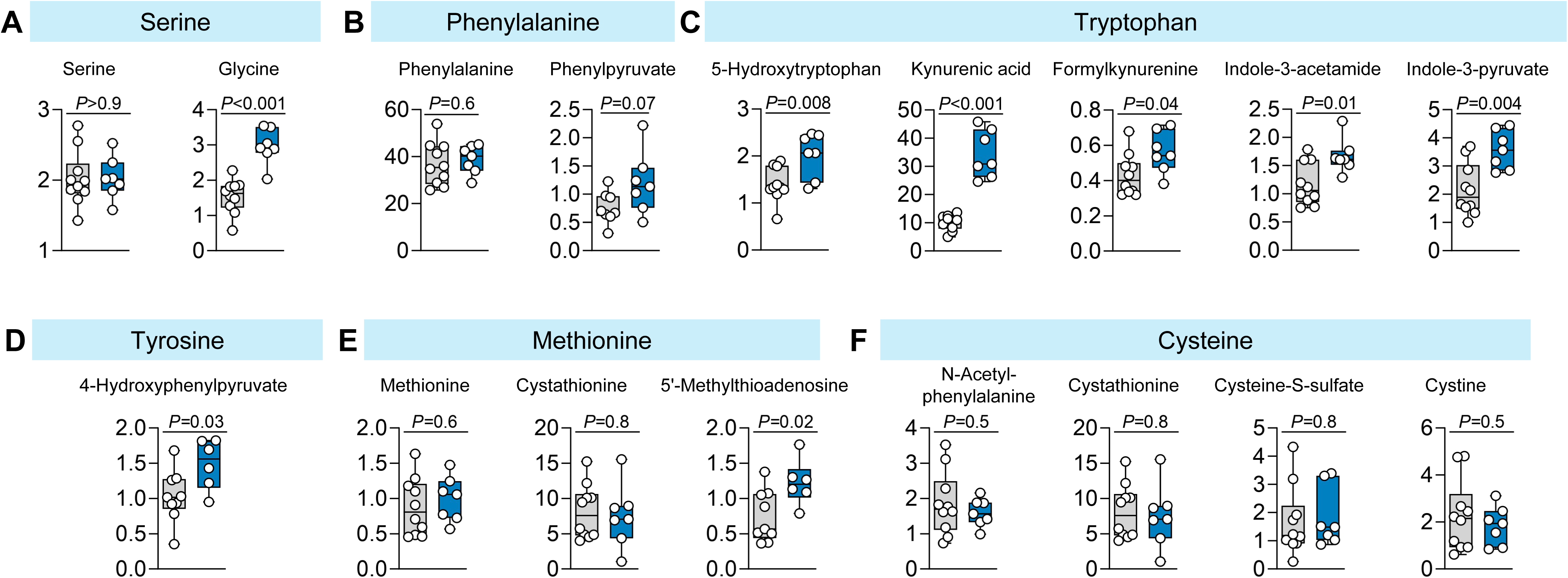
(Related to Figure 1B) Serum concentrations of amino acids and amino-acid metabolites in vehicle-injected mice fed SD (gray boxes; n = 9-10) or MD-1 (blue boxes; n = 6-7) for 25 days. Data are expressed as ion counts (mean ± SEM; Student’s *t* test).

**Supplemental Fig. S3.**
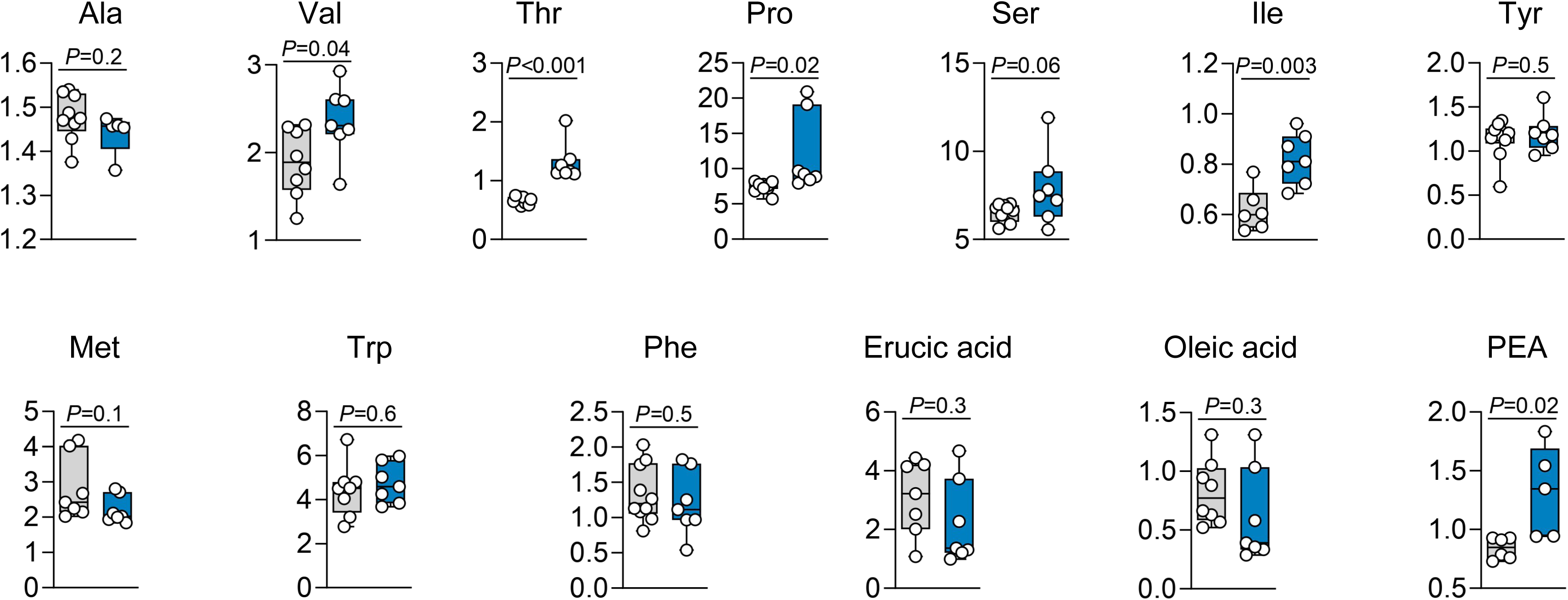
(Related to Figure 1D, E) Concentrations of MD-1 components in ipsilateral L4-L6 spinal cord of vehicle-injected mice fed SD (gray boxes; *n* = 7-10) or MD-1 (blue boxes; *n* = 5-7) for 25 days. Data are expressed as ion counts (mean ± SEM; Student’s *t* test).

**Supplemental Fig. S4.**
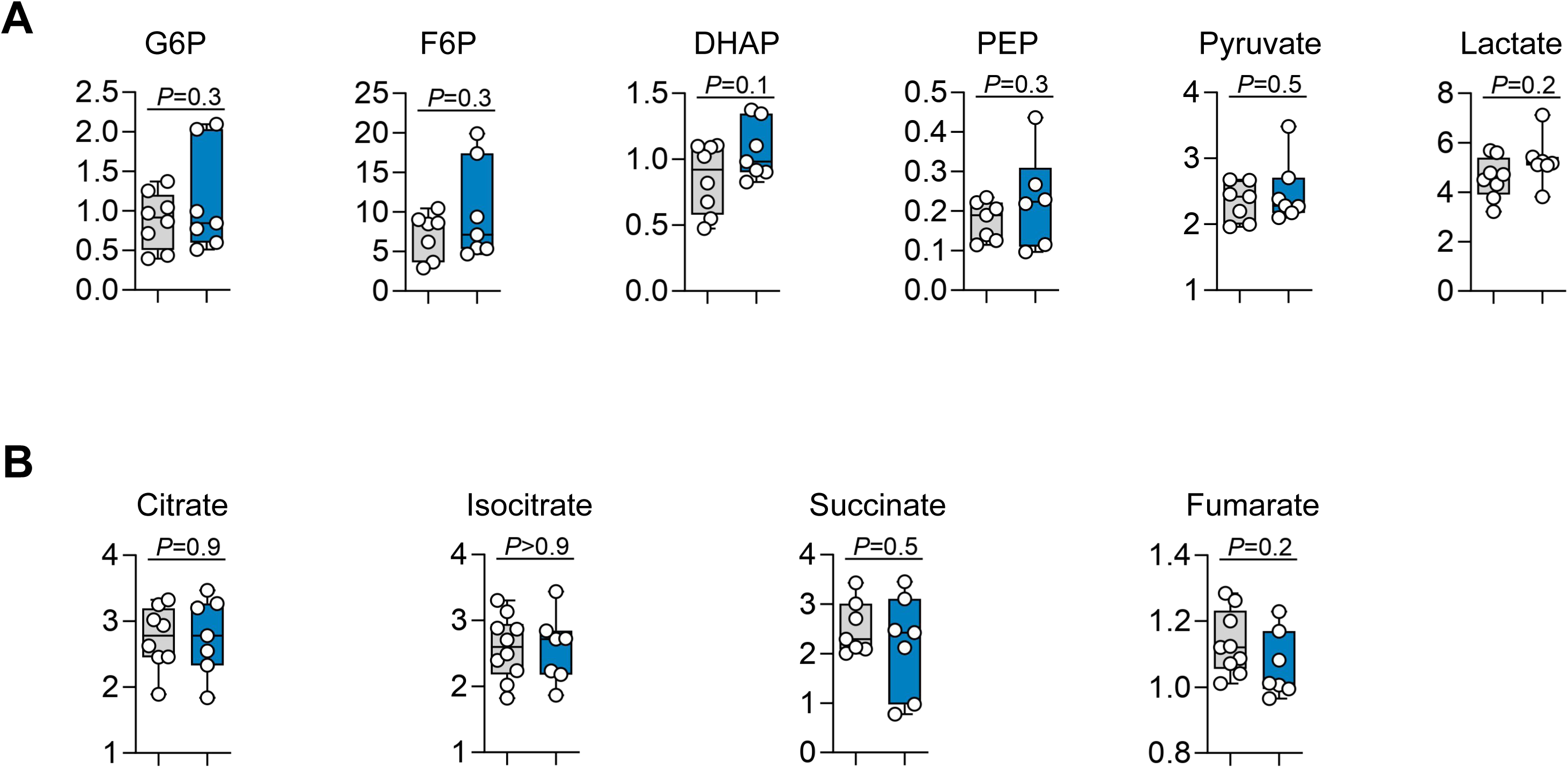
(Related to Figure 2) Concentrations of (A) glycolysis and (B) Krebs’ cycle metabolites in ipsilateral L4-L6 spinal cord of vehicle-injected mice fed SD (gray boxes) or MD-1 (blue boxes). Data are expressed as ion counts (mean ± SEM; *n* = 7-10 per group; Student’s *t* test).

**Supplemental Fig. S5.**
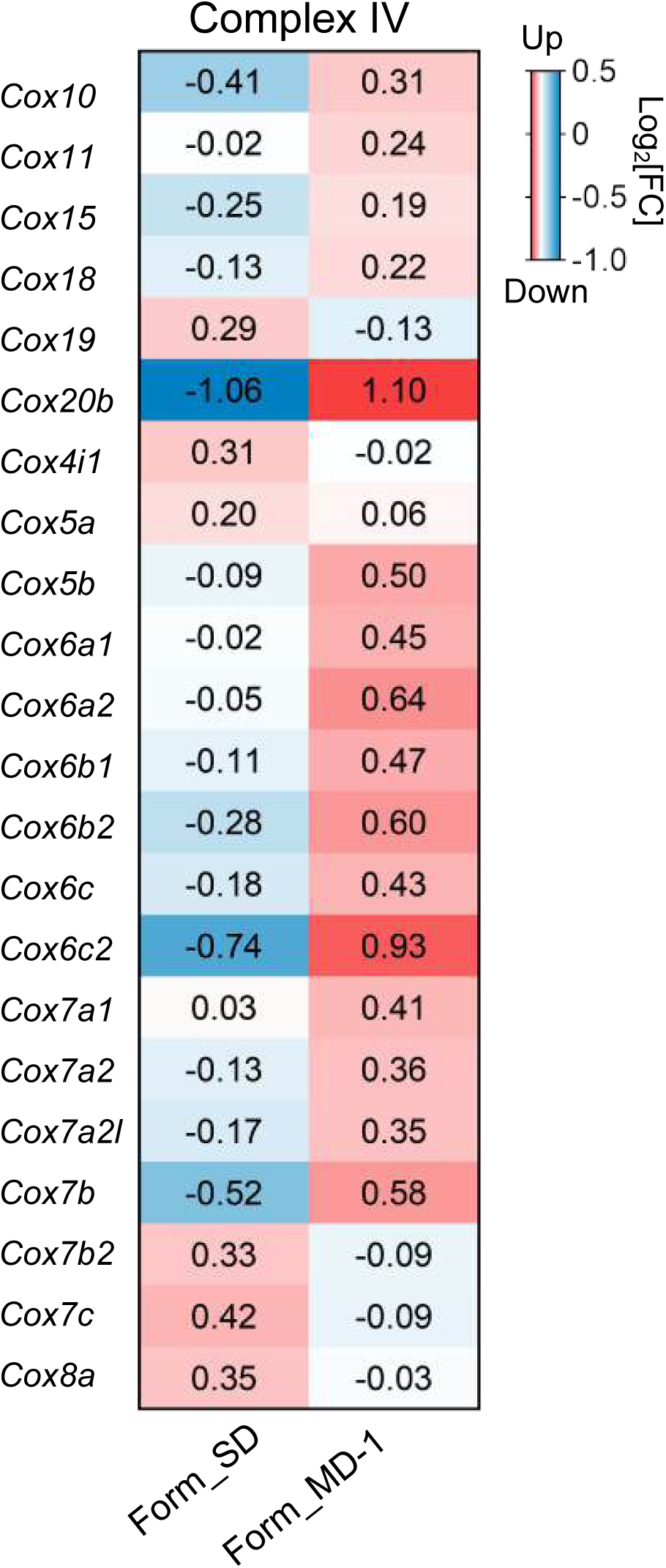
(Related to Figure 2D) Effects of formalin injection on Complex IV transcription in ipsilateral L4-L6 spinal cord of vehicle-injected mice fed SD or MD-1. Red: increase; blue: decrease. Data are expressed as log_2_ changes (formalin *vs* vehicle; n = 6 per group).

**Supplemental Fig. S6.**
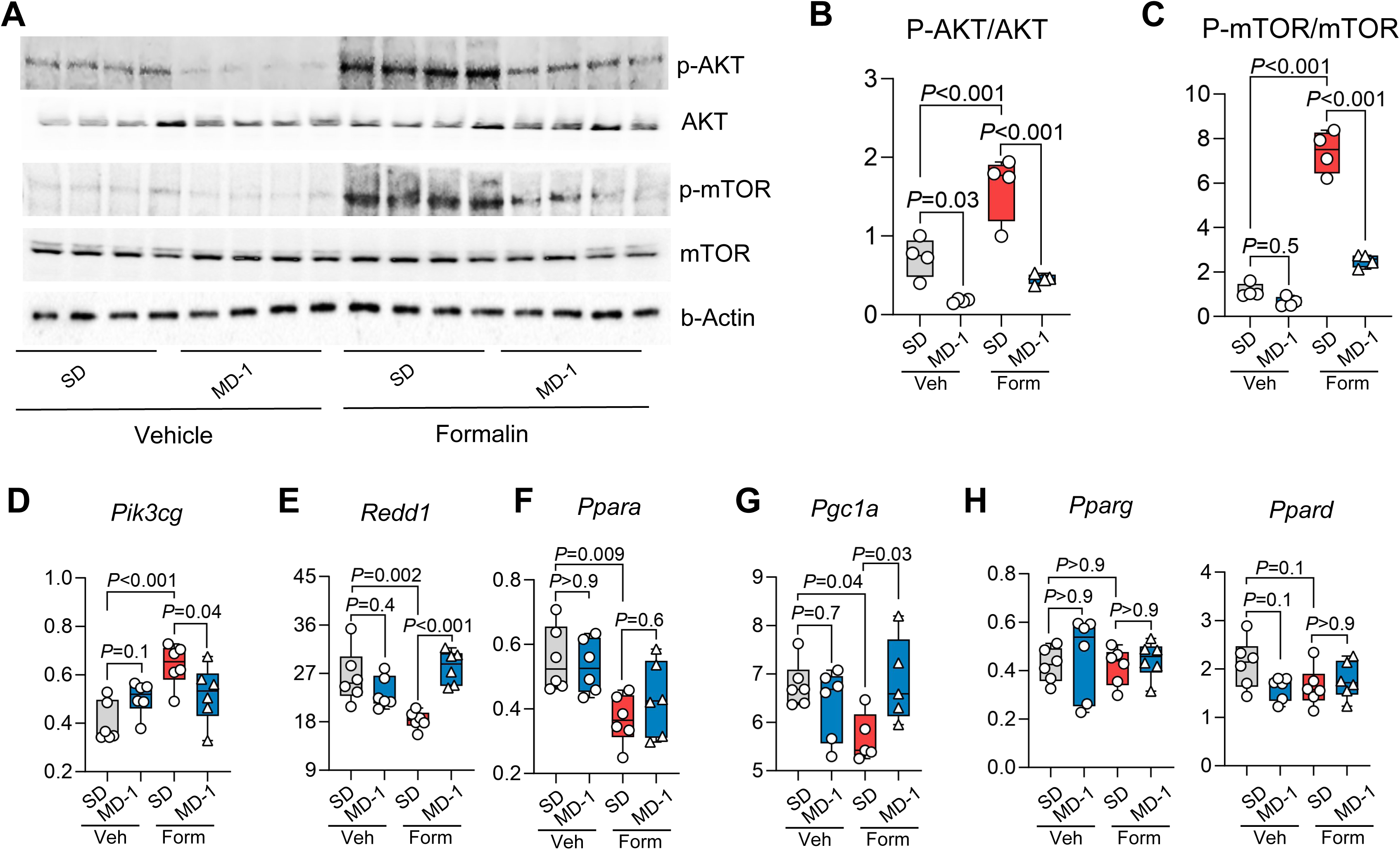
(Related to Figure 3) Formalin injection activates AKT/mTORC1 in ipsilateral L4-L6 spinal cord and MD-1 counters this activation. (A) Representative Western blot images showing levels of phospho-AKT (p-AKT), AKT, phospho-mTOR (p-mTOR), and mTOR in vehicle- or formalin-injected mice fed SD or MD-1. β-actin is the loading control. (B, C) Quantification of phospho-AKT (p-AKT/AKT) and phospho-mTOR (p-mTOR/mTOR) in vehicle (Veh)- or formalin (Form)-injected mice fed SD or MD-1. (D-H) Transcription of (D) *Pik3cg*, (E) *Redd1*, (F) *Ppara*, (G) *Ppargc1a* (*Pgc1a*), and (H) *Pparg* and *Ppard*. Data are expressed as mean ± SEM (n = 4-6 per group); one-way ANOVA with post hoc Šídák’s test. \**P*<0.05, \*\**P*<0.01, \*\*\**P*<0.001 compared to formalin/SD.

**Supplemental Fig. S7.**
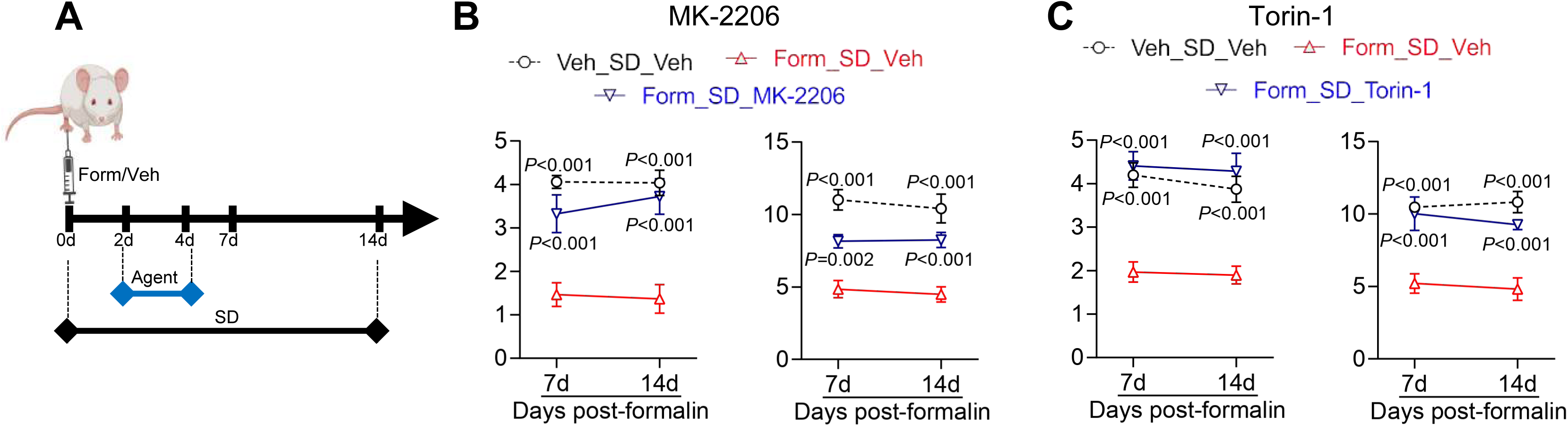
(Related to Figure 3) Effects of post-injury administration of AKT and mTOR inhibitors in formalin-injected mice. (A) Protocol: SD-fed mice were treated with AKT inhibitor MK-2206 (240 mg/kg, IP) or mTOR inhibitor Torin-1 (20 mg/kg, IP) on days 2-4 post-injection. Nocifensive behavior was monitored for the following two weeks. (B, C) Effects of (B) MK-2206 or (C) Torin-1 on contralateral hypersensitivity to mechanical (left) and thermal (right) stimuli. Data are expressed as mean ± SEM (n = 8-10 per group); two-way ANOVA with post hoc Šídák’s test. \**P*<0.05, \*\**P*<0.01, \*\*\**P*<0.001 compared to Form-SD.

**Supplemental Fig. S8.**
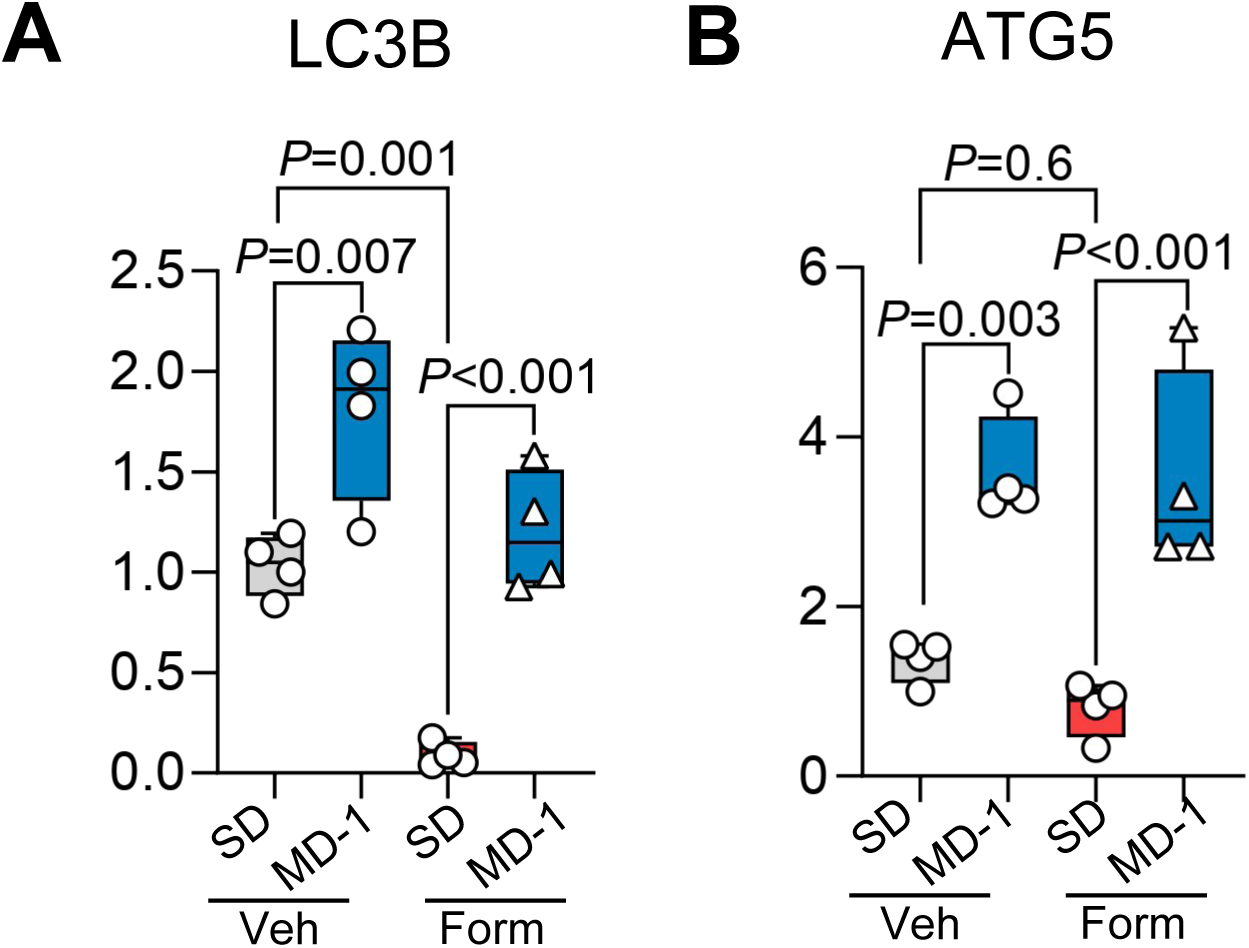
(Related to Figure 4B) Densitometry quantification of (A) LC3B and (B) ATG5 protein levels, normalized to β-actin. Data are expressed as mean ± SEM (*n* = 4 per group); one-way ANOVA with post hoc Šídák’s test.

**Supplemental Fig. S9.**
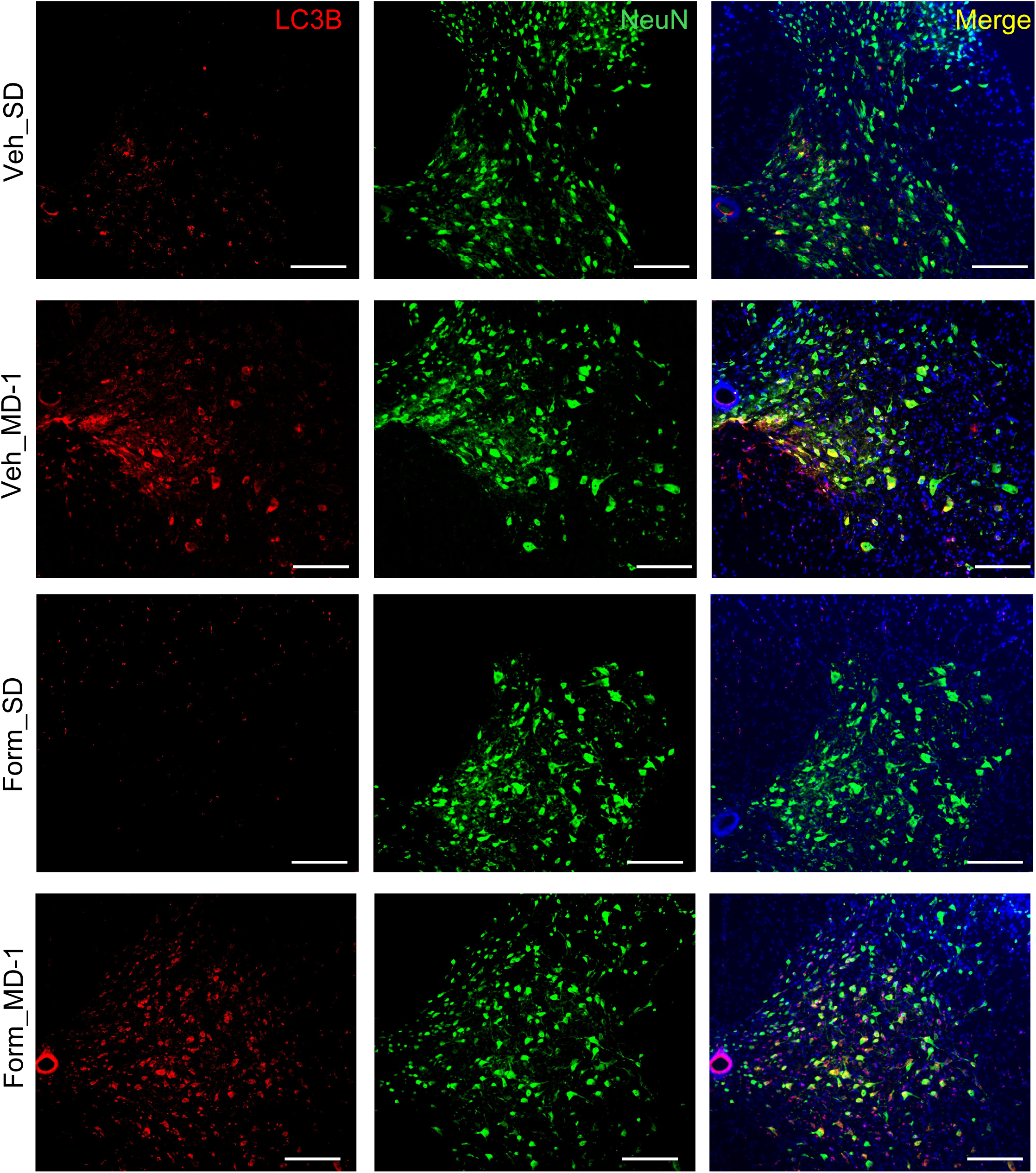
(Related to Figure 4C, D) Representative immunofluorescent images for LC3B (red) and neuronal marker NeuN (green) in L4-L6 spinal cord of vehicle- or formalin-injected mice fed either SD or MD-1. Nuclei are stained with DAPI. Magnification: 10x. Scale bar: 100 μm

**Supplemental Fig. S10.**
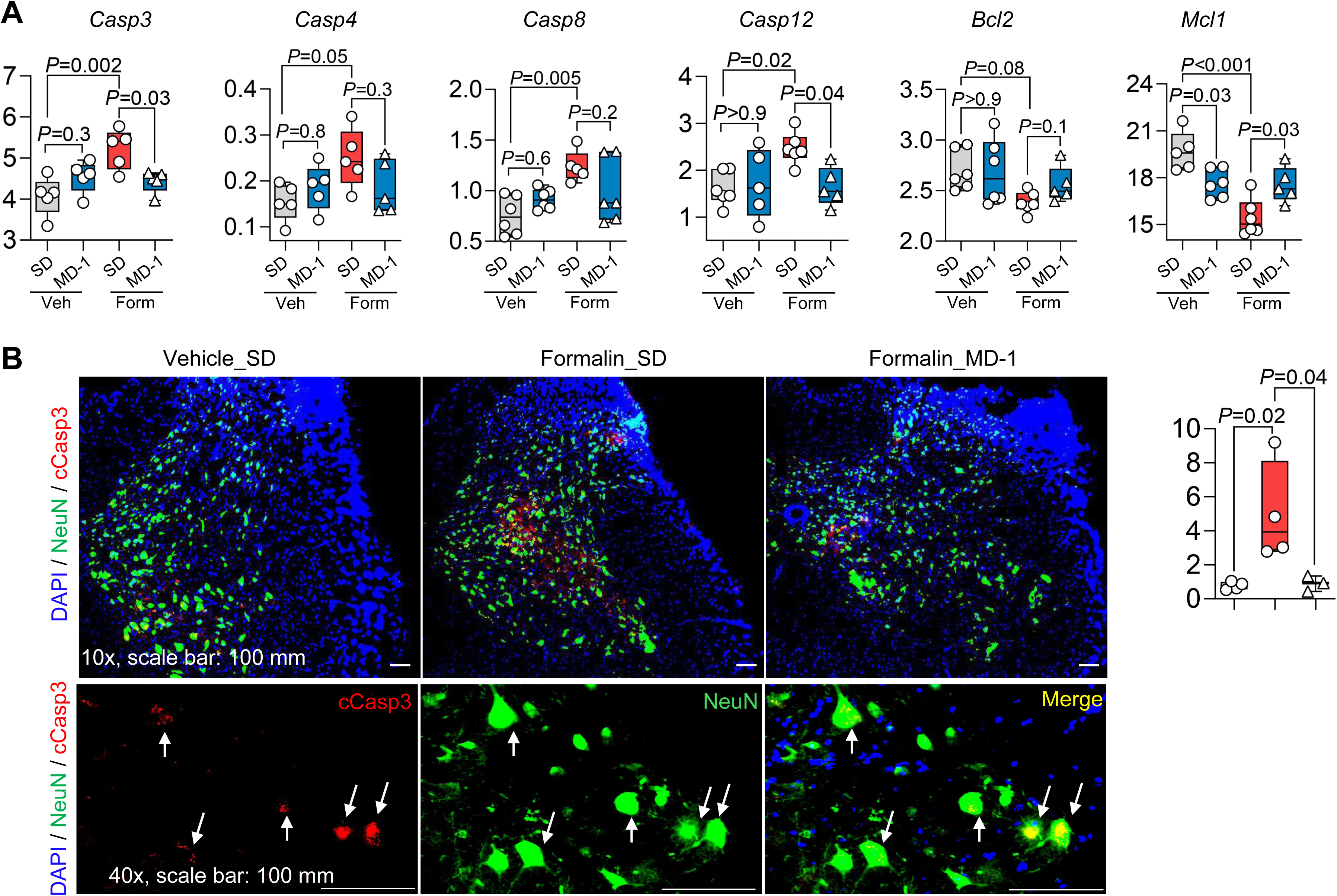
(Related to Figure 4) Formalin injection triggers apoptosis in L4-L6 spinal cord and MD-1 prevents this response. (A) Transcription of proapoptotic (*Casp3*, *Casp4*, *Casp8*, *Casp12*) and survival (*Bcl2*, *Mcl1*) genes in vehicle (Veh)- or formalin (Form)-injected mice fed SD or MD-1. Boxplots show individual and mean ± SEM data for vehicle-injected mice fed SD (gray boxes) and formalin-injected mice fed SD (red boxes) or MD-1 (blue boxes) (*n* = 5 per group); one-way ANOVA followed by Šídák’s test. (B) Immunofluorescence localization of caspase-3 in L4-L6 spinal cord. Top: representative images (10x magnification) showing caspase-3 (red) and NeuN (green) immunoreactivity. Nuclei are counterstained with DAPI (blue). Scale bar, 100 µm. Bottom row: images (40x magnification) highlighting the colocalization of activated caspase-3 with NeuN. (C) Quantification of activated caspase-3 immunofluorescence. Statistical significance was determined by one-way ANOVA with post hoc Šídák’s test.

**Supplemental Fig. S11.**
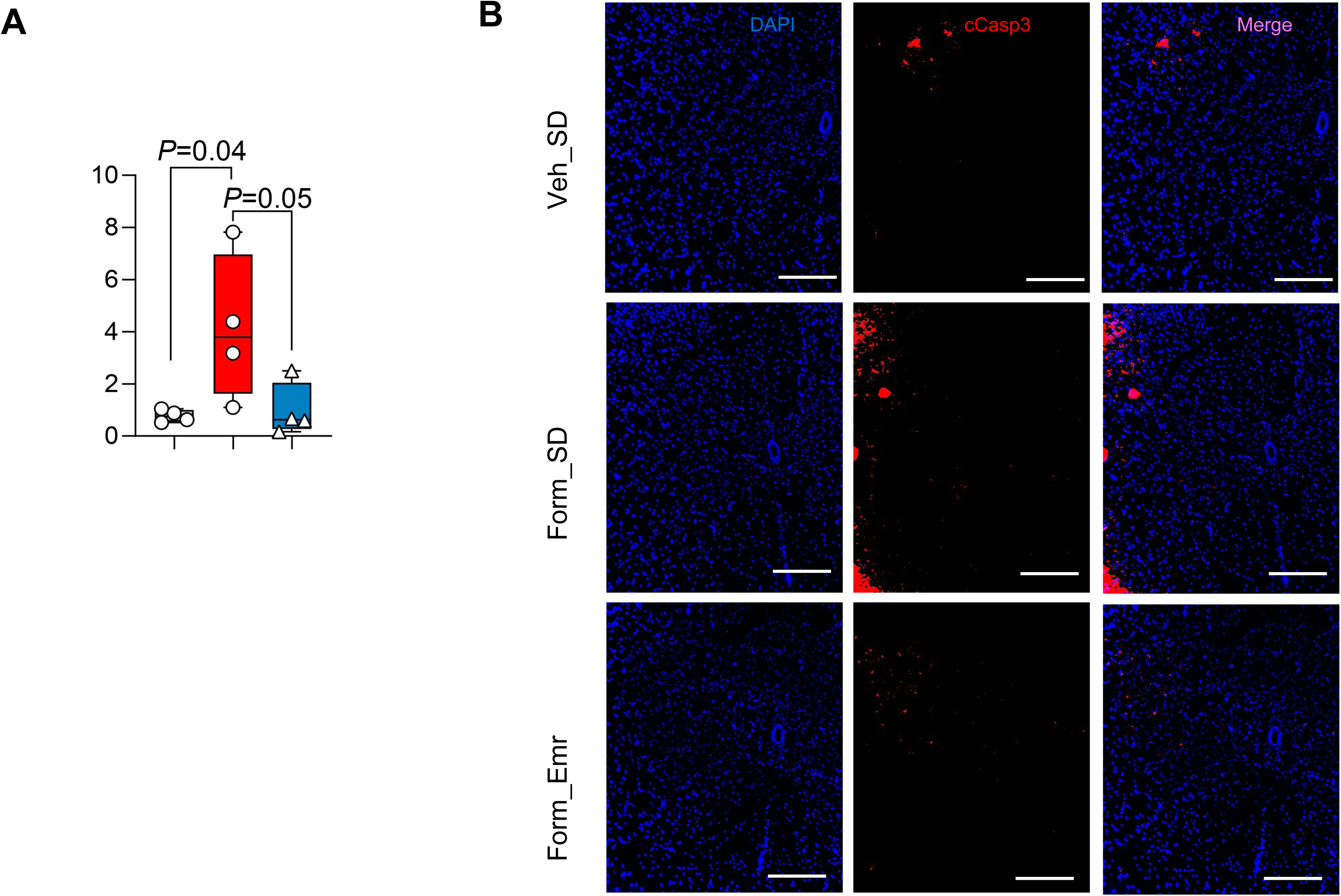
(Related to Figure 4) Caspase inhibitor emricasan (Emr, 3 mg/kg, IP) or its vehicle (Veh) was administered to SD-fed mice on days 2-4 following intraplantar injection of formalin (1%, v/v). (A) Quantification of cleaved caspase-3 (cCasp3) immunofluorescence in the L4-L6 spinal cord of vehicle (Veh)- or formalin (Form)-injected mice fed SD. (B) Representative immunofluorescent images for cCasp3 (red) in the L4-L6 spinal cord tissues. Nuclei are stained with DAPI. Magnification: 10x. Scale bar: 100 μm. Data are expressed as mean ± SEM and analyzed by two-way ANOVA followed by Dunnett’s test.

**Supplemental Fig. S12.**
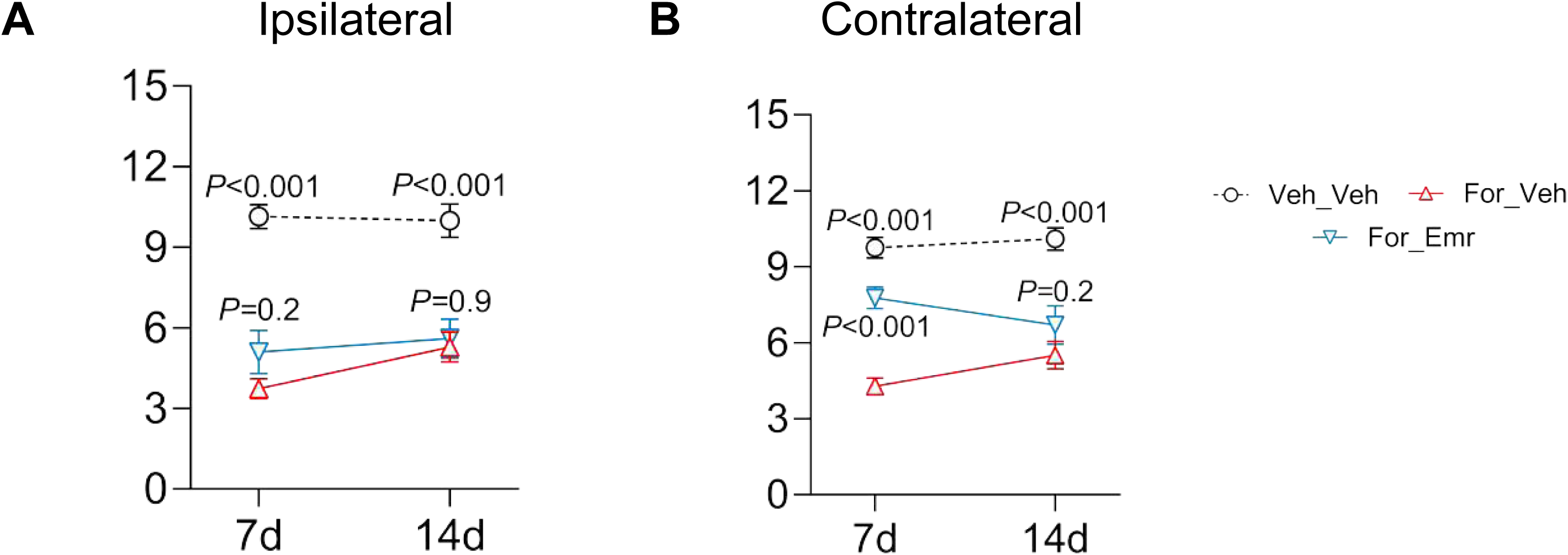
(Related to Figure 4) Caspase inhibitor emricasan (3 mg/kg, IP) or its vehicle (Veh) was administered to SD-fed mice on days 2-4 following intraplantar injection of formalin (1%, v/v). Thermal nociceptive thresholds were assessed in the ipsilateral (A) and contralateral (B) hind paws on days 7 and 14. Data are presented as mean ± SEM (n = 7–8 mice per group). Statistical significance was determined using two-way ANOVA followed by Dunnett’s multiple comparisons test. *P* < 0.05 compared to formalin-injected SD-fed mice (Form_SD).

**Supplemental Fig. S13.**
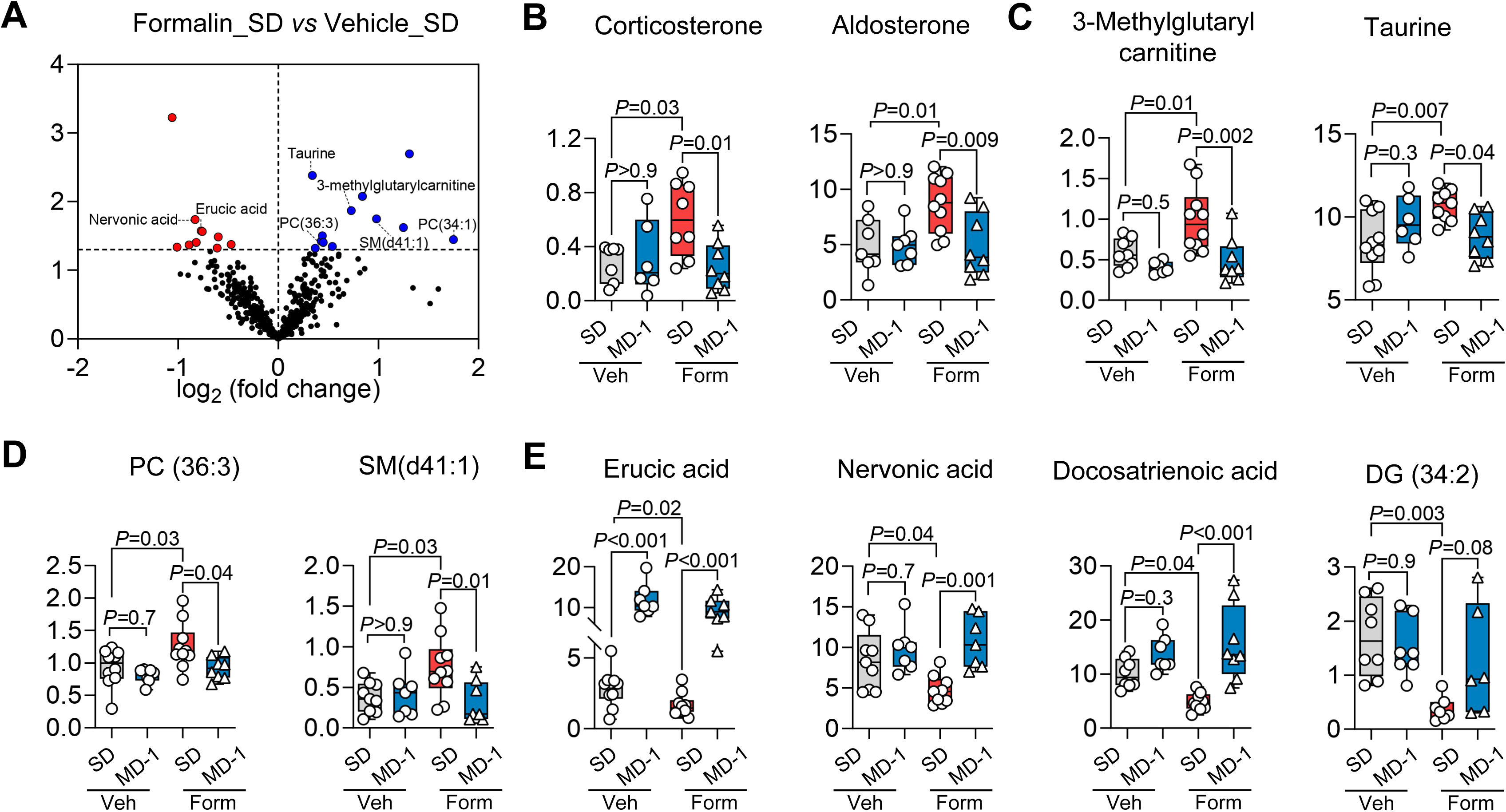
(Related to Figures 1, 2, and 3) Effects of hind-paw formalin injection and MD-1 exposure on the circulating metabolome. (A) Volcano plot showing metabolite changes in serum of SD-fed mice. Red dots: downregulated metabolites; blue dots: upregulated metabolites; black dots: metabolites with no significant change (*P* > 0.05) in formalin-injected SD-fed mice compared to vehicle-injected SD-fed controls. (B-E) Boxplots showing individual and mean ± SEM metabolite content (ion counts) in serum of formalin (Form)- or vehicle (Veh)-injected mice fed SD or MD-1 (*n* = 7-10 per group); one-way ANOVA followed by Šídák’s multiple comparisons test.

**Supplemental Fig. S14.**
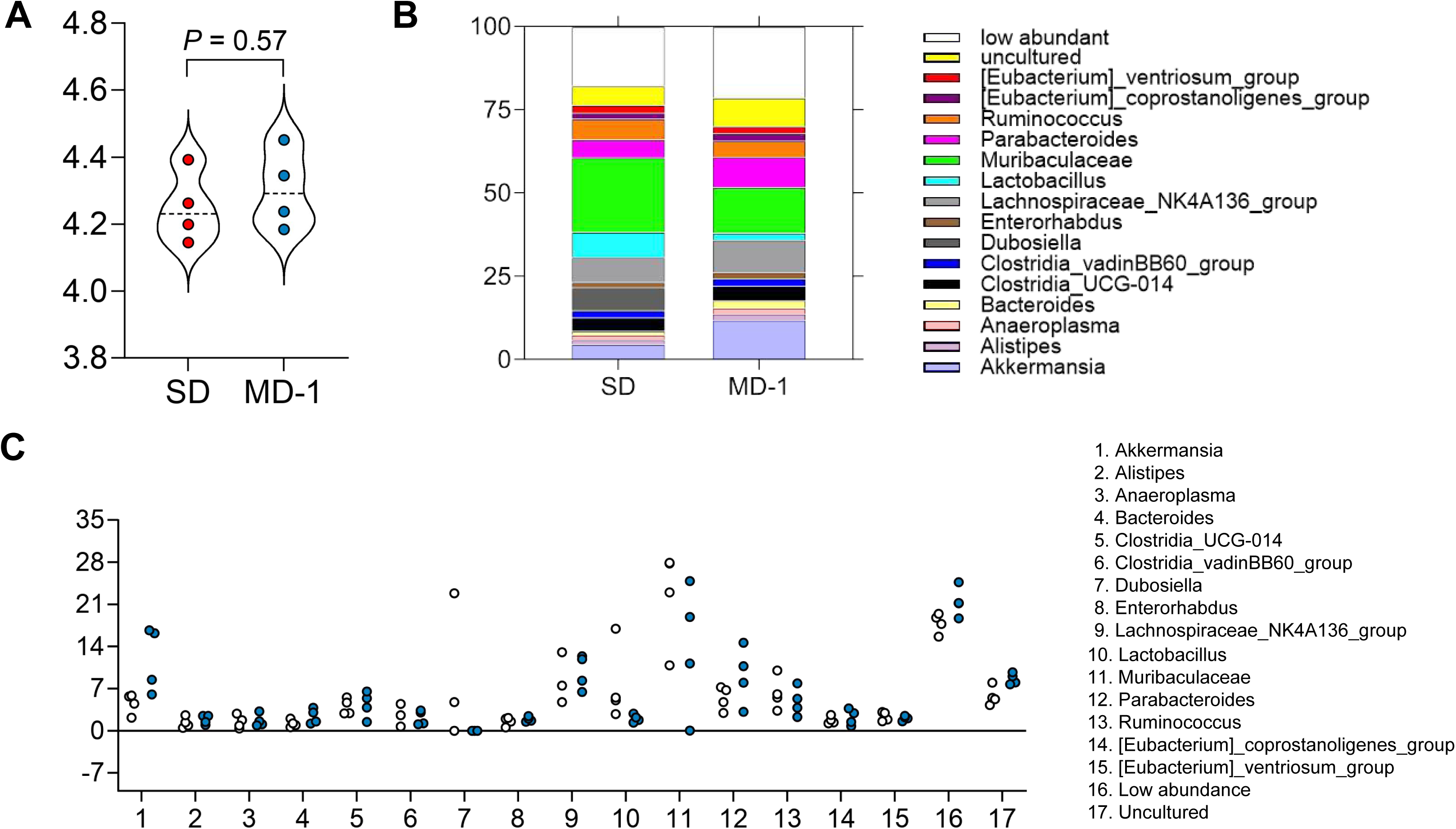
(Related to Figures 1, 2, and 3) MD-1 administration does not alter intestinal microbiome diversity or composition. (A) Shannon diversity index of fecal samples (*n* = 4 cages) from vehicle-injected mice fed SD or MD-1 for 25 days. (B) Relative abundance of predominant intestinal bacterial genera between groups. (C) Intestinal microbiome composition, showing bacterial genera that constitute >1% of total microbiome community.

**Supplemental Figure S15.**
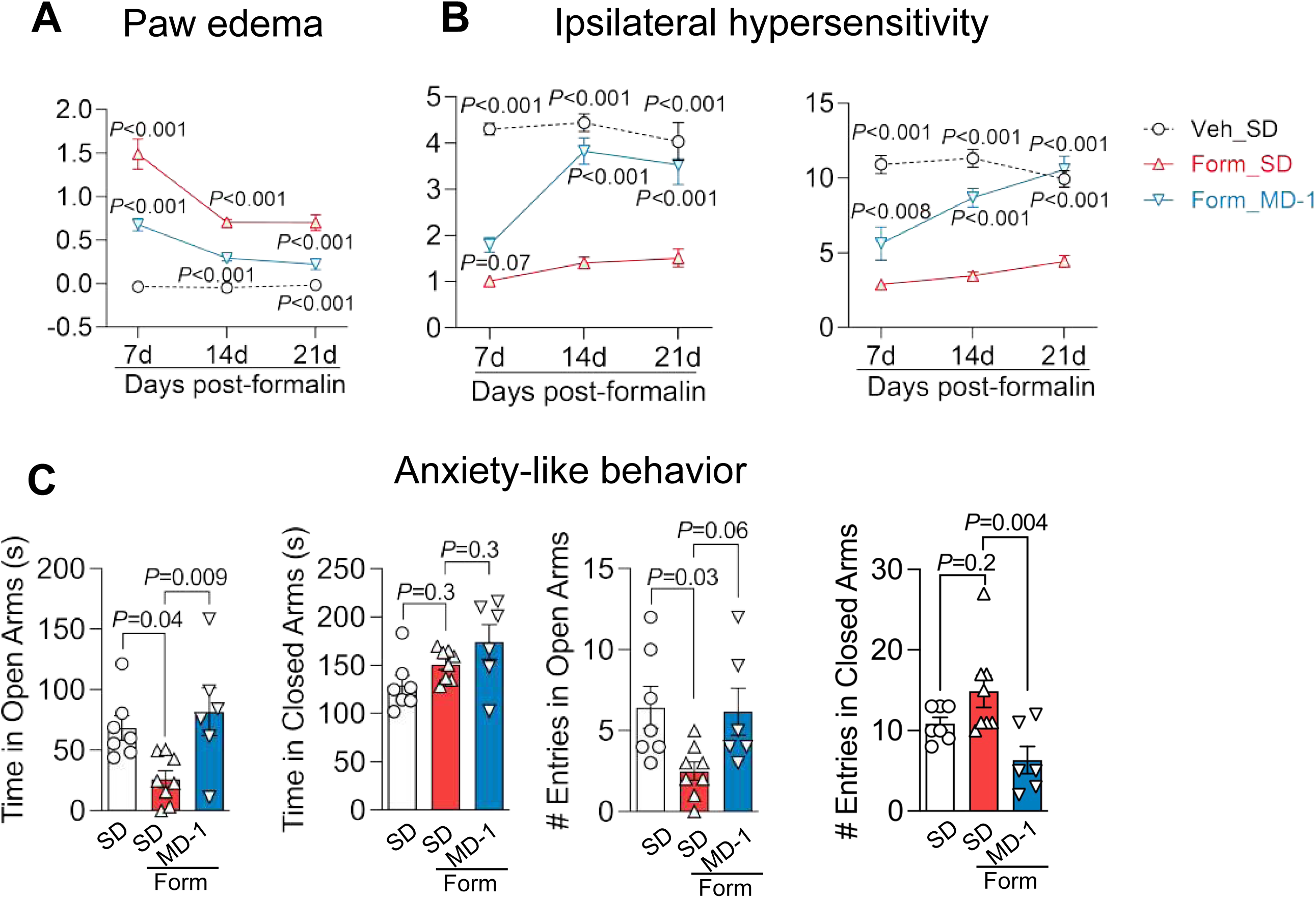
(Related to Figure 5A-E) Effects of MD-1 in formalin-injected mice fed SD (red symbols) or MD-1 (blue symbols). Open circles indicate vehicle-injected mice. (A, B) Time-course of (A) paw edema; and (B) ipsilateral hypersensitivity to mechanical (left) and thermal (right) stimuli. (C) Anxiety-like behavior (elevated plus maze). Data are expressed as mean ± SEM (*n* = 6-10 per group); one-way ANOVA with post hoc Šídák’s test; \**P*<0.05, \*\**P*<0.01, \*\*\**P*<0.001 compared to formalin/SD.

**Supplemental Figure S16.**
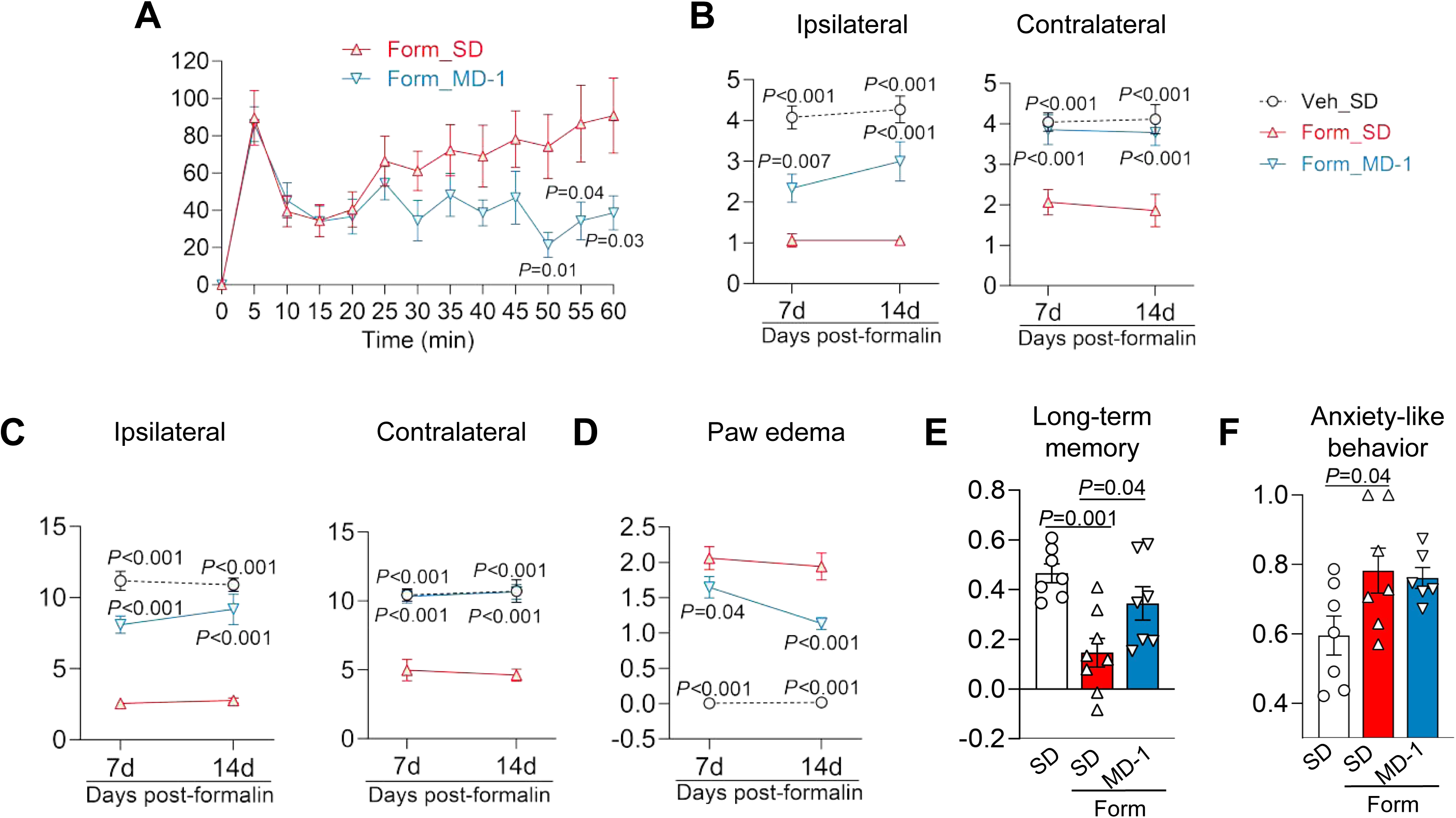
(Related to Figure 5A-E) Effects of MD-1 in female mice fed SD (red symbols) or MD-1 (blue symbols). (A) Acute nocifensive response to formalin. (B-D) Time-course of bilateral hypersensitivity to (B) mechanical and (C) thermal stimuli, and (D) paw edema. (E) Long-term memory (24-hour novel object recognition). (F) Anxiety-like behavior (elevated plus maze). Data are expressed as mean ± SEM (*n* = 5-10 per group); one-way ANOVA with post hoc Šídák’s test; \**P*<0.05, \*\**P*<0.01, \*\*\**P*<0.001 compared to formalin/SD.

**Supplemental Figure S17.**
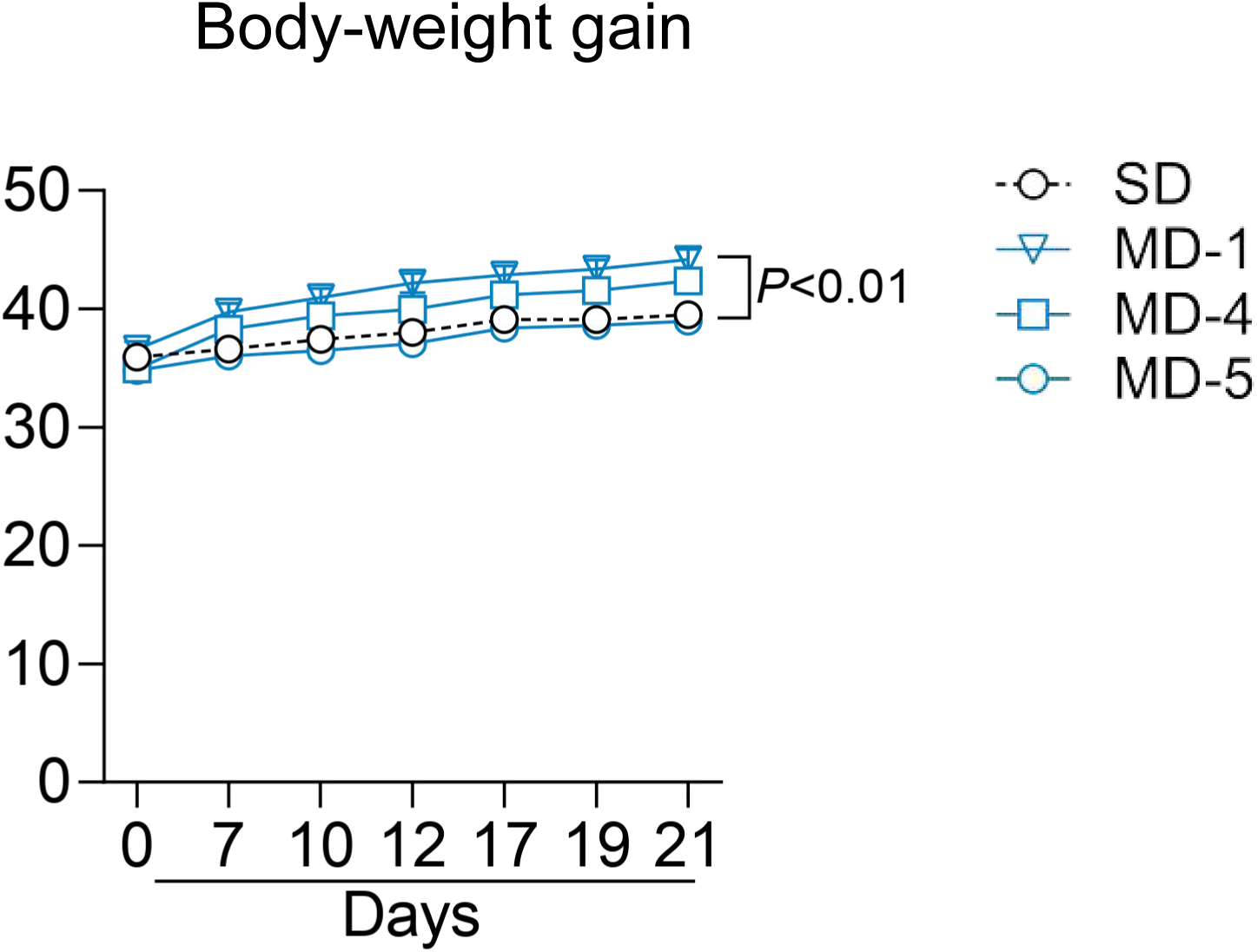
(Related to Figure 5A-E, J, and K) Body-weight trajectory in mice fed SD, MD-1, MD-4 or MD-5. Data are expressed as mean ± SEM (*n* = 8-10 per group); two-way ANOVA followed by post hoc Dunnett’s test. *P*<0.01 MD vs. SD.

**Supplemental Fig. S18.**
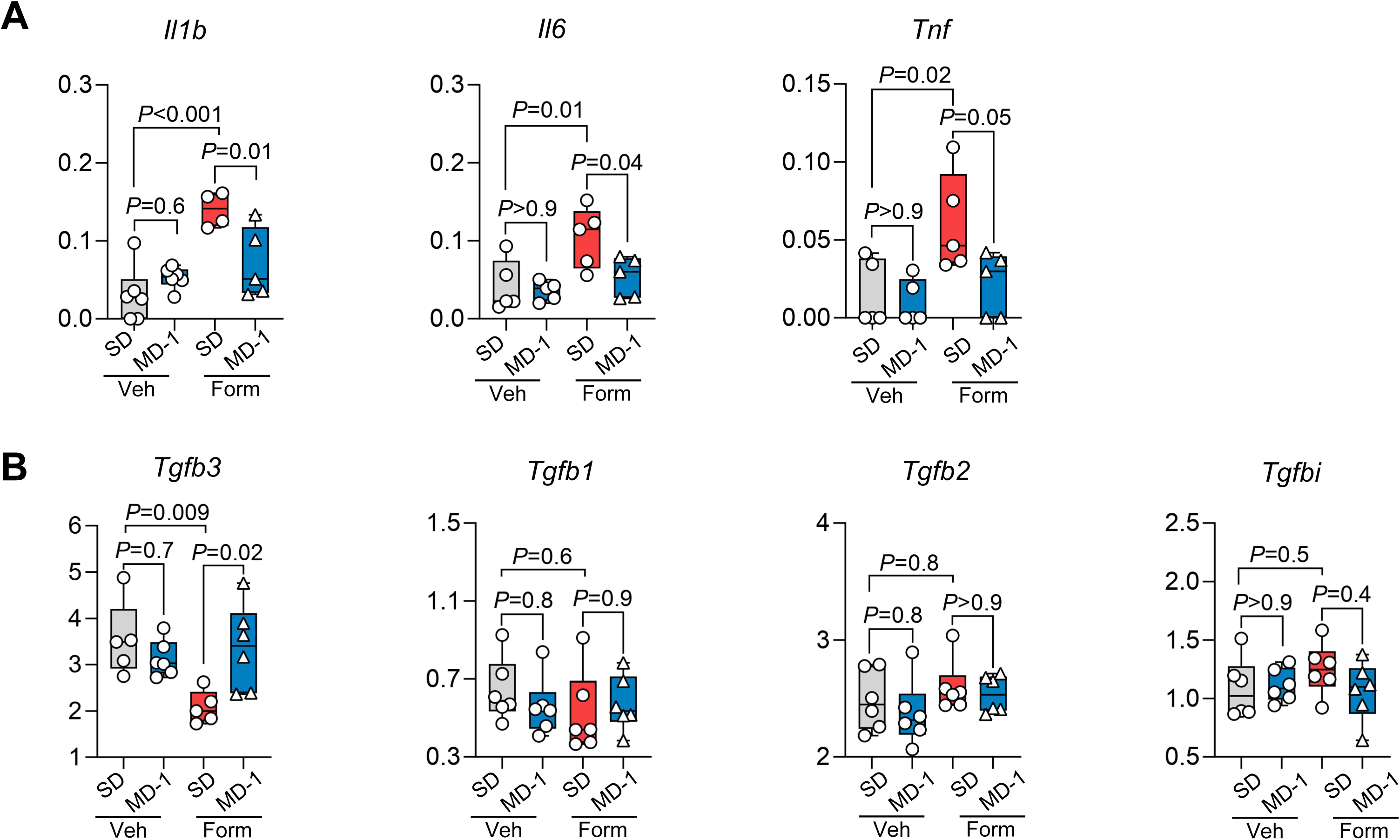
(Related to Figure 5A-E) Transcription of (A) proinflammatory and (B) tissue-reparative genes in ipsilateral L4-L6 spinal cord of formalin- or vehicle-injected mice fed SD or MD-1. Boxplots show individual and mean ± SEM data for vehicle-injected mice fed SD (gray boxes) and formalin-injected mice fed SD (red boxes) or MD-1 (blue boxes) (*n* = 4-6 per group); one-way ANOVA followed by Šídák’s test.

**Supplemental Fig. S19.**
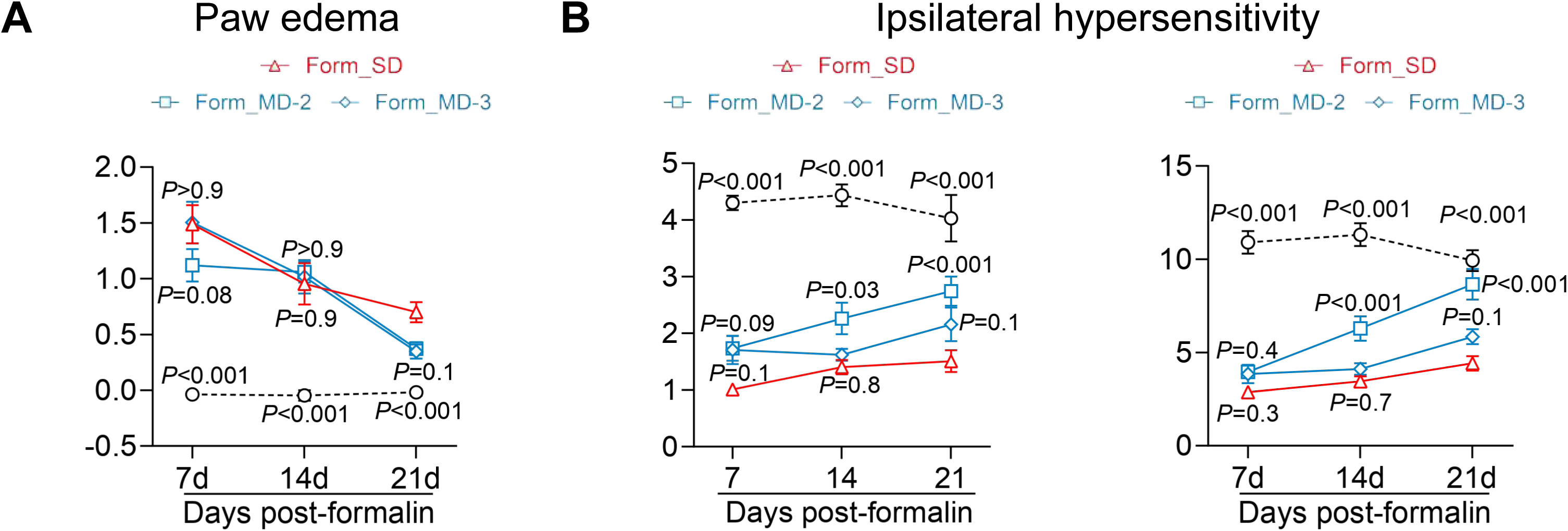
(Related to Figure 5G-I) Effects of MD-1 in formalin-injected mice fed SD (red triangles), MD-2 (blue squares), or MD-3 (blue diamonds). Open circles indicate vehicle-injected mice fed SD. Time-course of (A) paw edema and (B) ipsilateral hypersensitivity to mechanical (left) and thermal (right) stimuli. Data are expressed as mean ± SEM (*n* = 8-10 per group); one-way ANOVA followed by Šídák’s test. \**P*<0.05, \*\**P*<0.01, \*\*\**P*<0.001 compared to formalin/SD.

**Supplemental Fig. S20.**
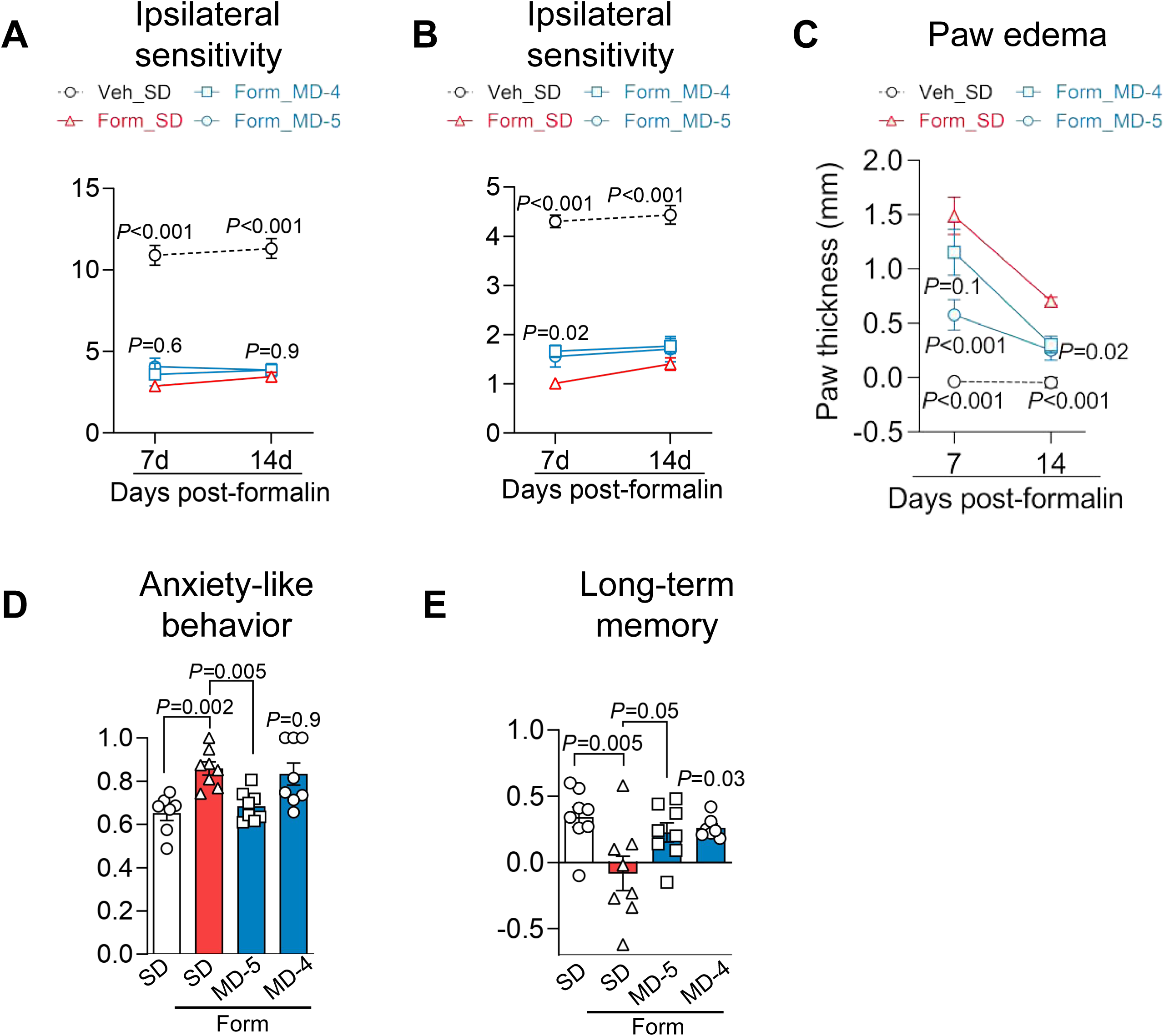
(Related to Figure 5J-K) Effects of MD-4 and MD-5 in formalin-injected mice fed SD (red triangles), MD-4 (blue squares), or MD-5 (blue circle). (A-C) Time-course of (A) paw edema and (B, C) ipsilateral hypersensitivity to (B) thermal and (C) mechanical stimuli. (D) Anxiety-like behavior (elevated plus maze). (E) Long-term memory (24-hour novel object recognition). Data are expressed as mean ± SEM (*n* = 8-10 per group); one- or two-way ANOVA followed by Šídák’s test. \**P*<0.05, \*\**P*<0.01, \*\*\**P*<0.001 compared to formalin/SD.

**Supplemental Fig. S21.**
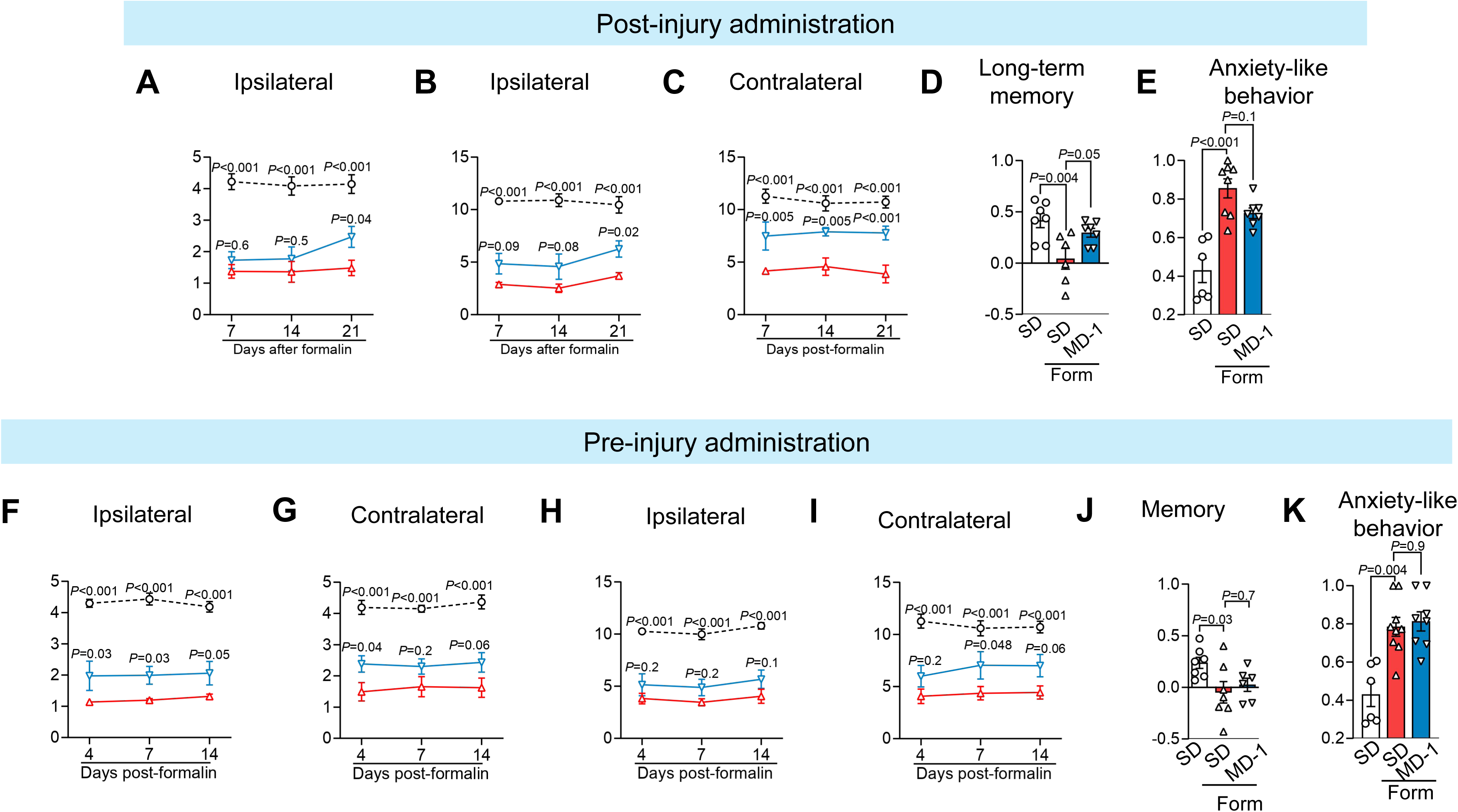
(Related to Figure 5L, M) Effects of timing of MD-1 exposure in formalin-injected mice fed SD (red symbols) or MD-1 (blue symbols). Open circles indicate vehicle-injected mice fed SD. (A-C) Time-course of the effects of post-injury MD-1 administration on (A, B) ipsilateral hypersensitivity to (A) mechanical and (B) thermal stimuli; and (C) contralateral thermal hypersensitivity. (D) Effects of post-injury MD-1 administration on long-term memory (24-hour novel object recognition). (E) Effects of post-injury MD-1 administration on anxiety-like behavior (elevated plus maze). (F-K) Time-course of the effects of pre-injury MD-1 administration on (F) ipsilateral mechanical hypersensitivity, (G) contralateral mechanical hypersensitivity, (H) ipsilateral thermal hypersensitivity, (I) contralateral thermal hypersensitivity. (J) Effects of pre-injury MD-1 administration on long-term memory (24-hour novel object recognition). (K) Effects of pre-injury MD-1 administration on anxiety-like behavior (elevated plus maze).. Data are expressed as mean ± SEM (*n* = 8-10 per group); one- or two-way ANOVA with Šídák’s test. \**P*<0.05, \*\**P*<0.01, \*\*\**P*<0.001 compared to formalin/SD.

**Supplemental Fig. S22.**
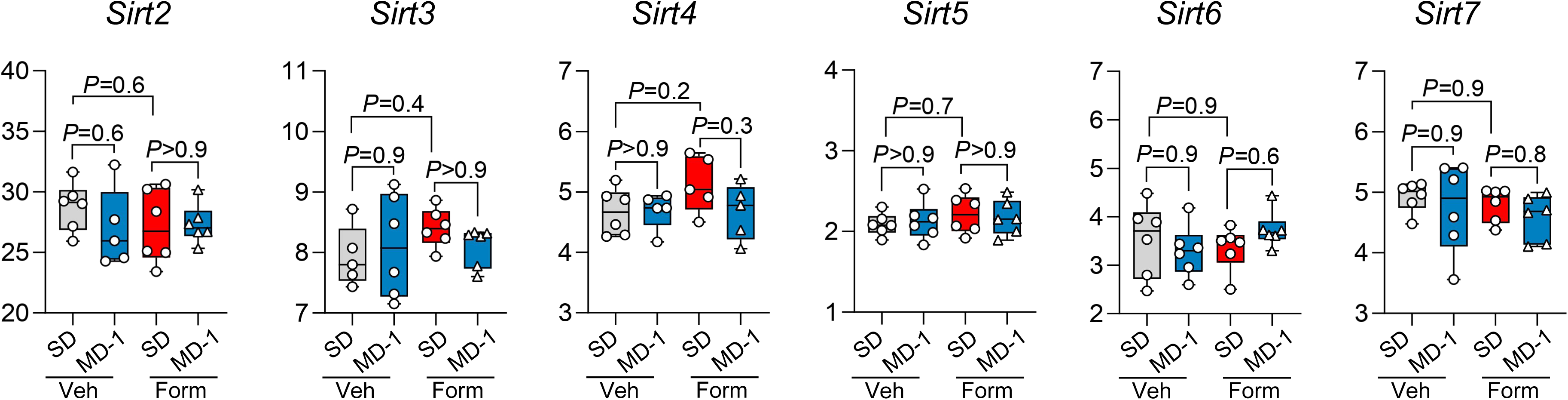
(Related to Figure 6A) Transcription of sirtuin family members (*Sirt2*-*Sirt7*) in ipsilateral L4-L6 spinal cord of vehicle (Veh)- or formalin (Form)-injected mice fed SD or MD-1. Boxplots show individual and mean ± SEM data for vehicle-injected mice fed SD (gray boxes) and formalin-injected mice fed SD (red boxes) or MD-1 (blue boxes). Data are expressed as mean ± SEM (*n* = 5-6 per group); one-way ANOVA with post hoc Šídák’s test.

**Supplemental Fig. S23.**
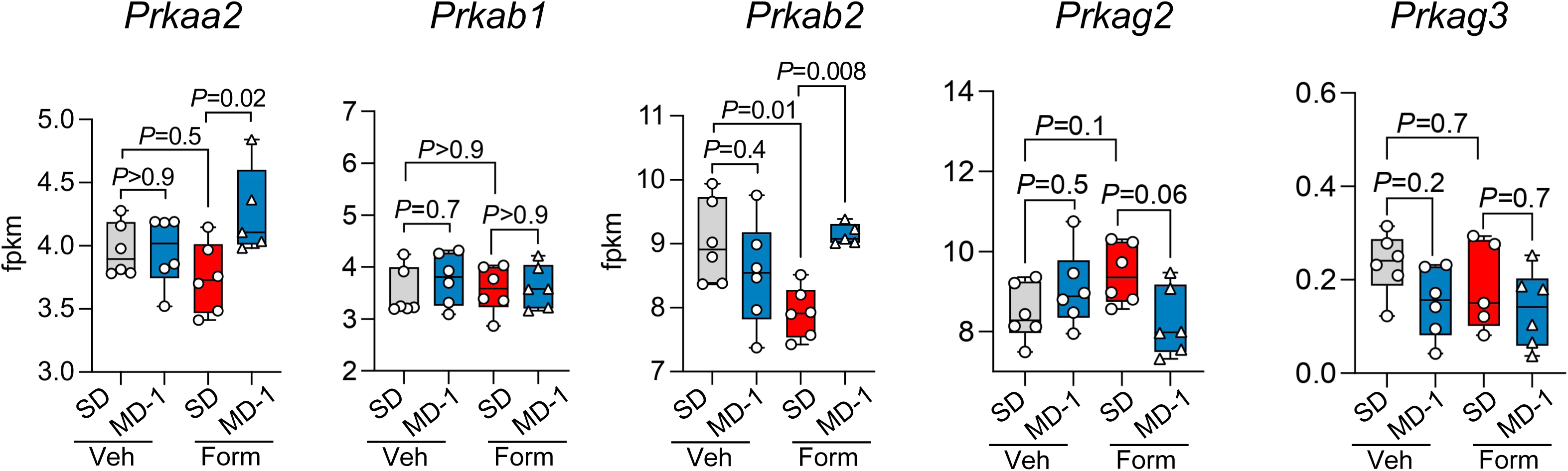
(Related to Figure 6F-H) Transcription of AMPK subunits in ipsilateral L4-L6 spinal cord of vehicle (Veh)- or formalin (Form)-injected mice fed SD or MD-1. Boxplots show individual and mean ± SEM data for vehicle-injected mice fed SD (gray boxes) and formalin-injected mice fed SD (red boxes) or MD-1 (blue boxes). Data are expressed as mean ± SEM (*n* = 5-6 per group); one-way ANOVA with post hoc Šídák’s test.

**Supplemental Fig. S24.**
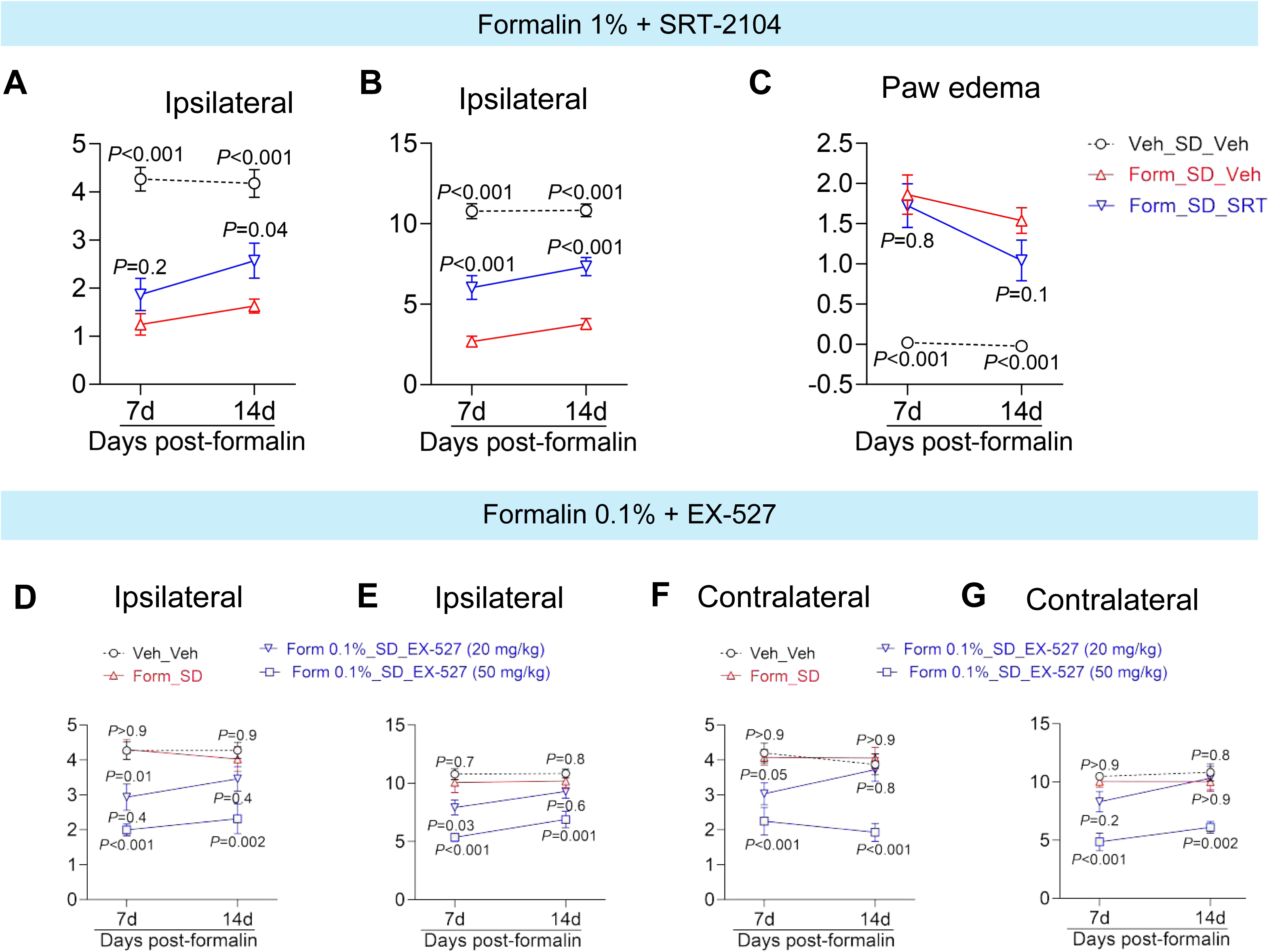
(Related to Figure 6I-K) (A-C) Effects of SIRT1 activator SRT-2104 (100 mg/kg, IP) on ipsilateral hypersensitivity to (A) mechanical and (B) thermal stimuli and (C) paw edema in formalin-injected mice fed SD (red symbols) or MD-1 (blue symbols). Open circles indicate vehicle-injected mice fed SD. (D-G) Effects of SIRT1 inhibitor EX-527. EX-527 (20 and 50 mg/kg, IP) or its vehicle (Veh) was administered to SD-fed mice on days 2-4 following intraplantar injection of formalin (0.1%, v/v). Hypersensitivity to mechanical (D, F) and thermal (E, G) stimuli were assessed in the ipsilateral and contralateral hind paws on days 7 and 14. Data are presented as mean ± SEM (*n* = 7–8 mice per group). Statistical significance was determined using two-way ANOVA followed by Šídák’s test. *P* < 0.05 compared to formalin-injected SD-fed mice (Form_SD).

**Supplemental Fig. S25.**
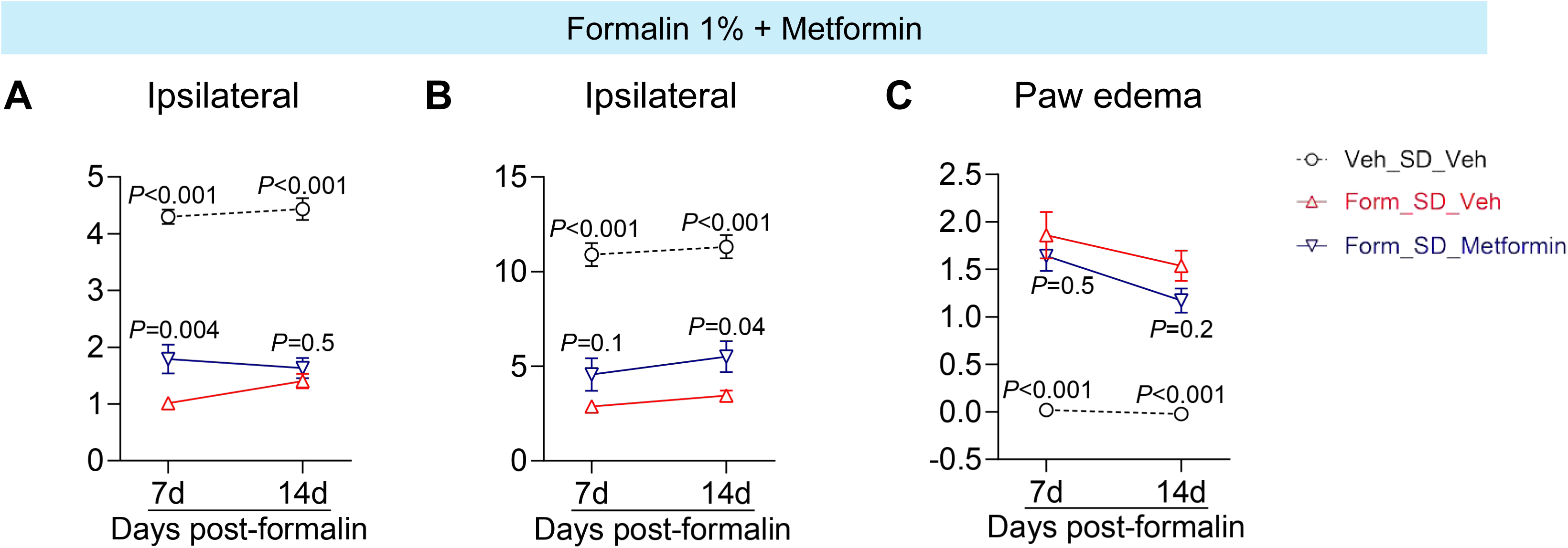
(Related to Figure 6) Effects of AMPK activator metformin (200 mg/kg, IP) on ipsilateral hypersensitivity to (A) mechanical and (B) thermal stimuli and (C) paw edema. Data are expressed as mean ± SEM (*n* = 7-10 per group); two-way ANOVA with Šídák’s test; \**P*<0.05, \*\**P*<0.01, \*\*\**P*<0.001 vs. formalin/SD.

**Supplemental Fig. S26.**
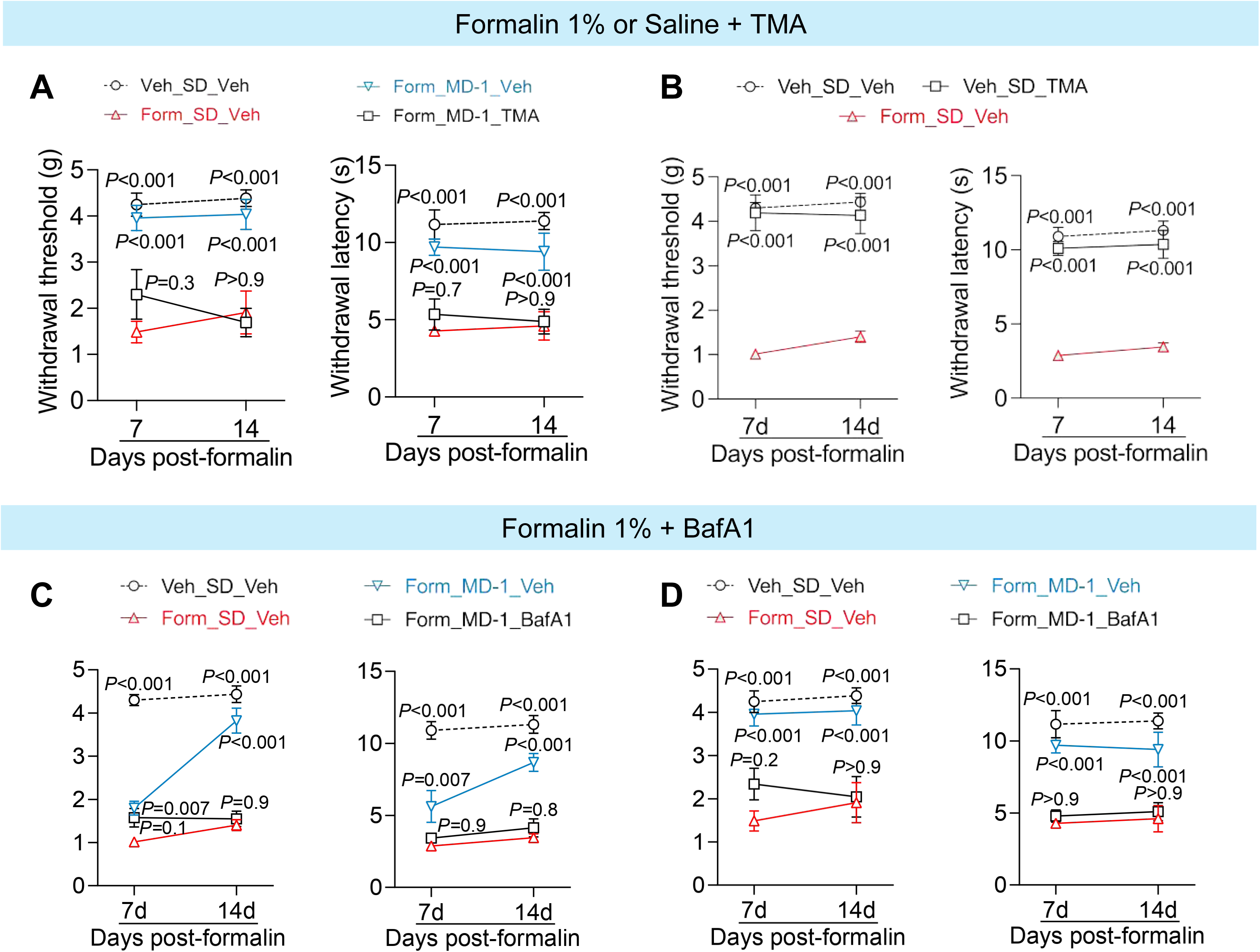
(Related to Figure 6) Effects of autophagy inhibition on the response to MD-1. (A) Treatment with autophagy inhibitor TMA (30 mg/kg, IP) on days 2-4 after formalin (1% vol/vol) negated the protective effects of MD-1 on formalin-induced mechanical (left) and thermal (right) hypersensitivity in the contralateral paws. (B) Effects of TMA on ipsilateral mechanical (left) and thermal (right) hypersensitivity in saline-injected mice. (C-D) Post-injury administration of autophagy inhibitor bafilomycin A1 (BafA1, 1 mg/kg, IP) negated the protective effects of MD-1 on formalin-induced ipsilateral (C) and contralateral (D) hypersensitivity to mechanical (left) and thermal (right) stimuli. Open circles: vehicle (veh)/SD/veh; red triangles: formalin/SD/veh; blue triangles: formalin/MD-1/veh; open squares, formalin/MD-1/TMA or BafA1. Statistical significance was determined by one- or two-way ANOVA followed by Dunnett or Šídák multiple comparisons test. \**P*<0.05, \*\**P*<0.01, \*\*\**P*<0.001 compared to formalin-SD (*n* = 7-10).

**Supplemental Fig. S27.**
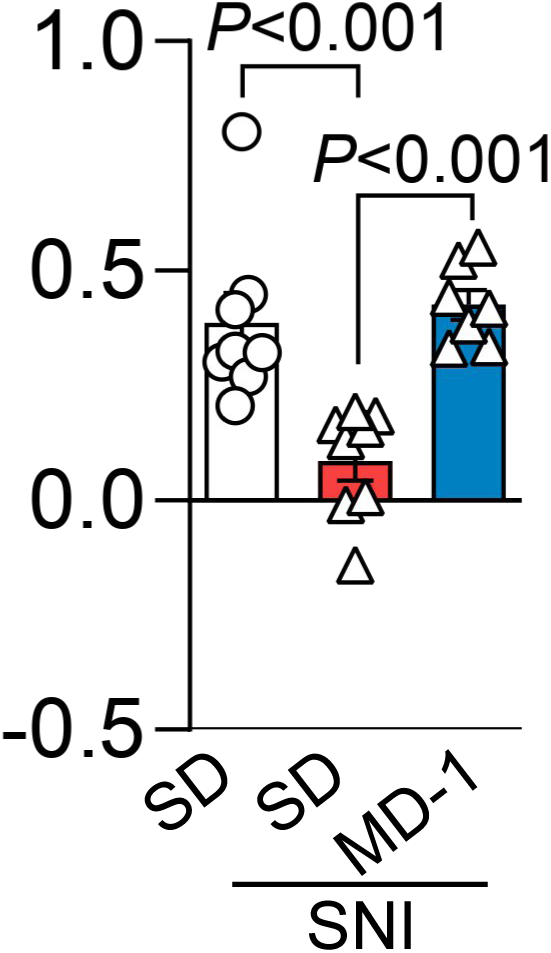
(Related to Figure 7A-E) Effects of MD-1 administration on SNI-induced long-term memory deficits (24-hour novel-object recognition). The test was performed on post-SNI day 21. Data are expressed as mean ± SEM (n = 7-8 per group); one-way ANOVA with Šídák’s test; \**P*<0.05, \*\**P*<0.01, \*\*\**P*<0.001 vs. SNI-SD group.

**Supplemental Fig. S28.**
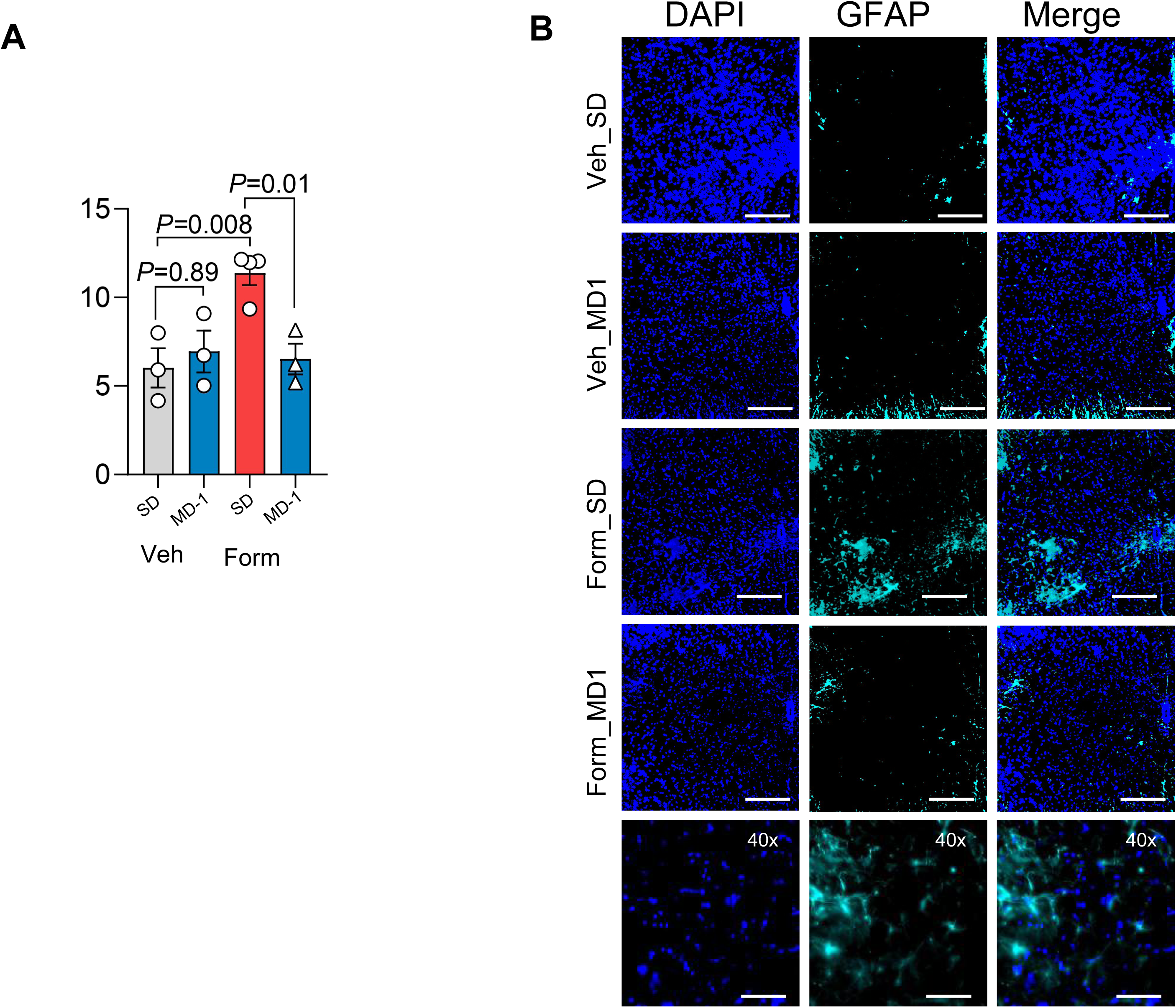
(Related to Figure 7A-E) (A) Quantification of GFAP immunofluorescence in the L4-L6 spinal cord of vehicle (Veh)- or formalin (Form)-injected mice fed SD or MD-1. (B) Representative immunofluorescent images for GFAP (cyan) in the L4-L6 spinal cord tissues. Nuclei are stained with DAPI. Magnification: 10x and 40x. Scale bar: 100 μm. Data are mean ± SEM and analyzed by two-way ANOVA followed by Dunnett’s test.

**Supplemental Fig. S29.**
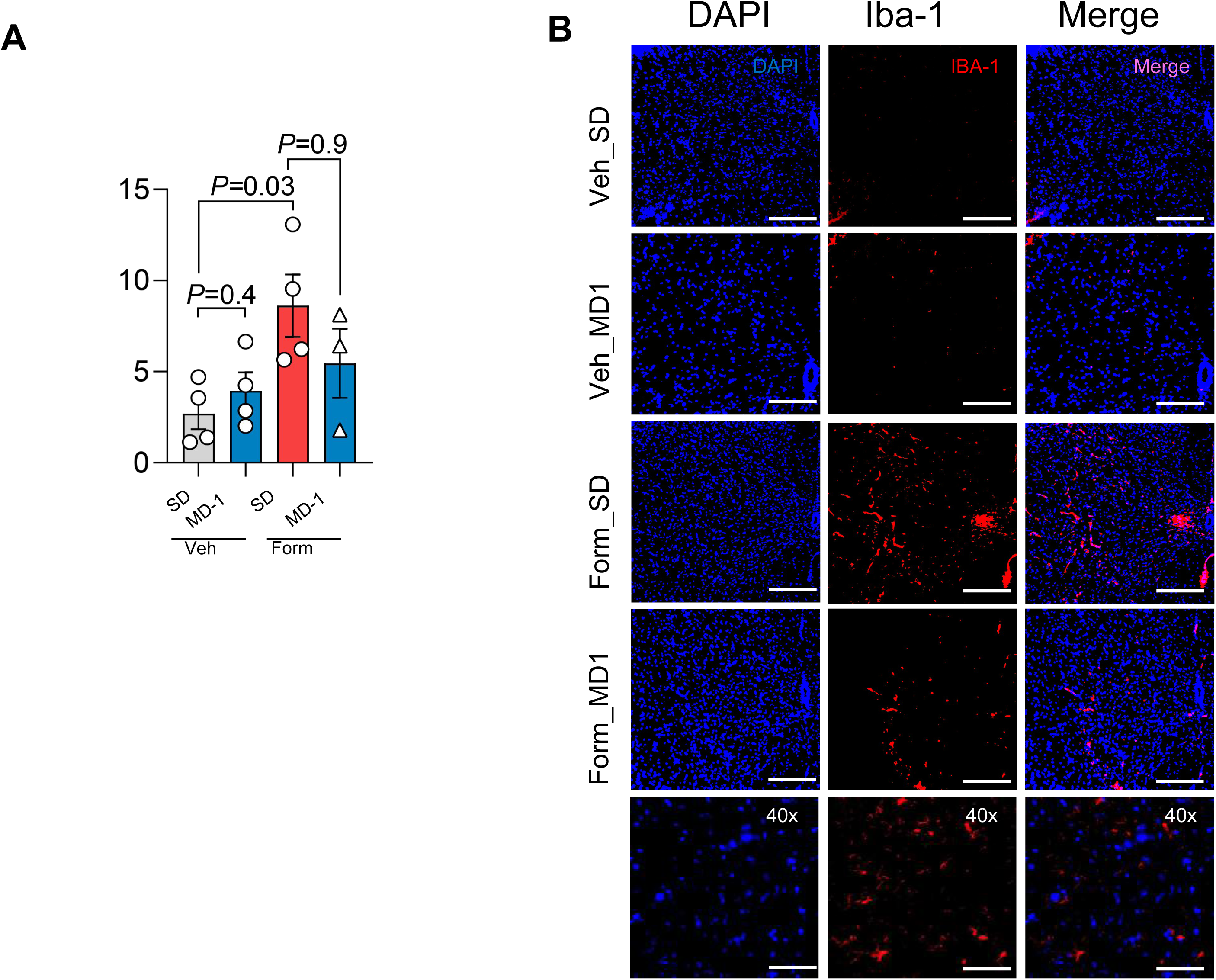
(Related to Figure 7A-E) (A) Quantification of IBA-1 immunofluorescence in the L4-L6 spinal cord of vehicle (Veh)- or formalin (Form)- injected mice fed SD or MD-1. (B) Representative immunofluorescent images for IBA-1 (red) in the L4-L6 spinal cord tissues. Nuclei are stained with DAPI. Magnification: 10x and 40x. Scale bar: 100 μm. Data are mean ± SEM and analyzed by two-way ANOVA followed by Dunnett’s test.

**Supplemental Fig. S30.**
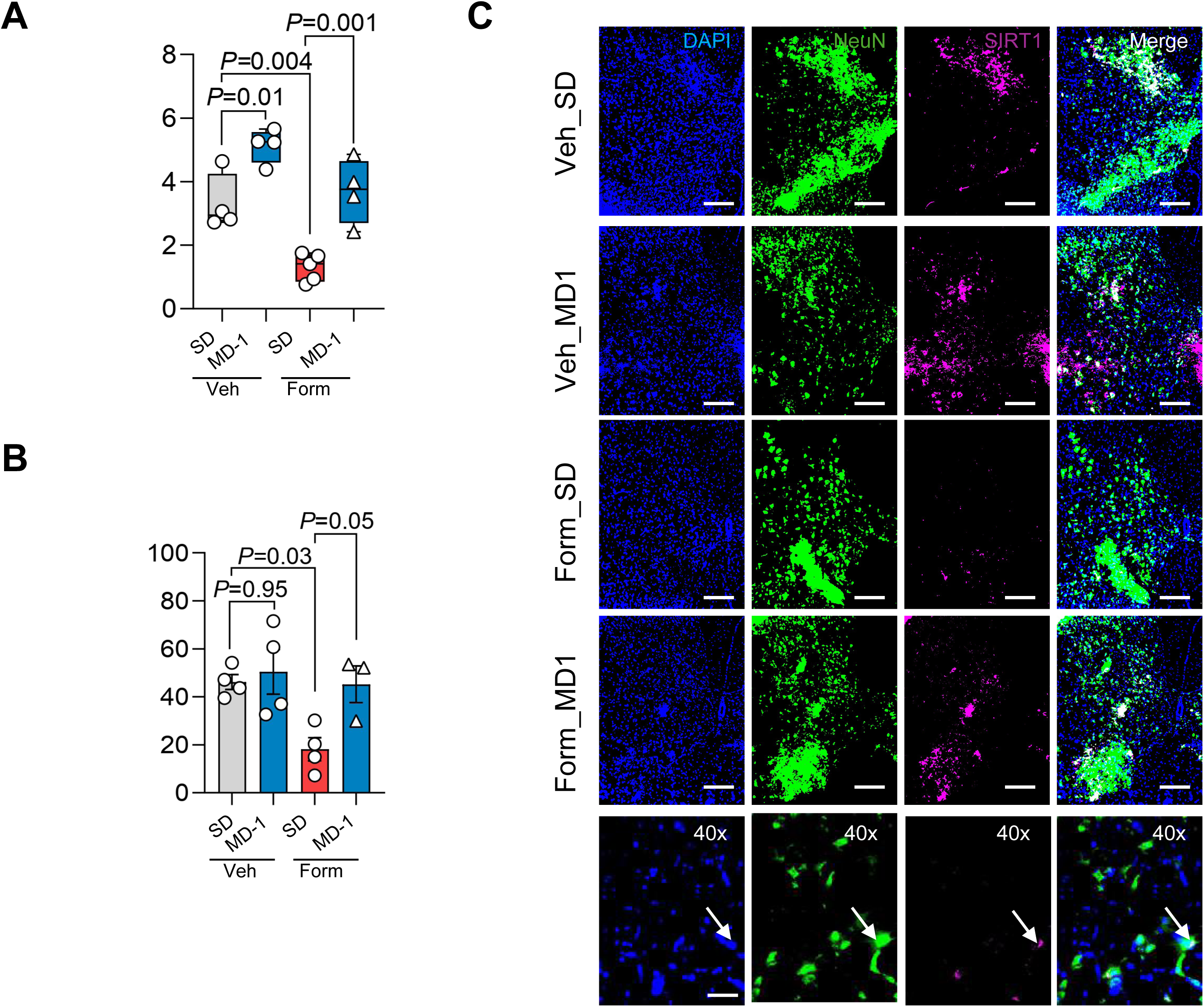
(Related to Figure 7A-E) (A-B) Quantification of SIRT1 immunofluorescence in the L4-L6 spinal cord of vehicle (Veh)- or formalin (Form)-injected mice fed SD or MD-1. (C) Representative immunofluorescent images for SIRT1 (magenta) and neuronal marker NeuN (green) in the L4-L6 spinal cord tissues. Nuclei are stained with DAPI. Magnification: 10x. Scale bar: 100 μm. Arrows indicate SIRT1 and NeuN colocalization at 40x. Data are mean ± SEM and analyzed by two-way ANOVA followed by Dunnett’s test.

**Supplemental Fig. S31.**
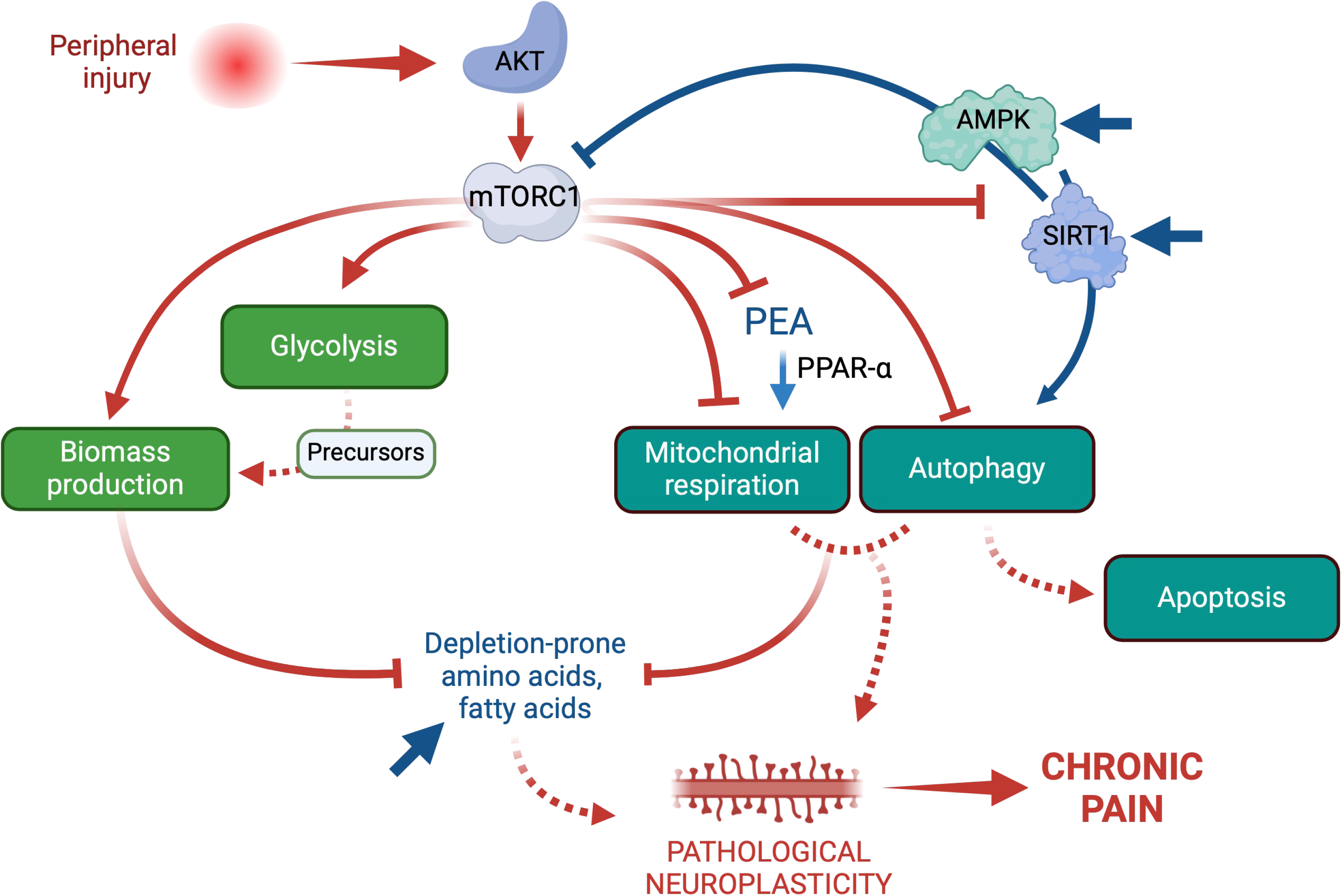
(Related to Figure 1-7) Metabolic reprogramming and autophagy suppression in spinal cord drive chronic pain development after injury. This hypothetical model illustrates how injury-induced alterations in spinal cord metabolism may contribute to chronic pain development. Peripheral tissue damage activates AKT/mTORC1 in afferent segments of the spinal cord, initiating a molecular cascade (red arrows) that simultaneously enhances aerobic glycolysis and biomass production while suppressing mitochondrial respiration, SIRT1/AMPK activity, and autophagy. These metabolic changes create a resource allocation crisis, depleting a local pool of amino acids and fatty acids essential for the maintenance of normal synaptic plasticity. At the same time, increased expression and activity of the enzyme N-acylethanolamine acid amidase lower PEA levels, impairing PPAR-α signaling, exacerbating mitochondrial dysfunction, and promoting neuroinflammation. These events cooperate to foster the progression to pain chronicity. MD-1 administration (blue arrows) blocks this molecular cascade and prevents the transition to chronic pain by restoring SIRT1/AMPK signaling and reactivating autophagy. Whether neuronal apoptosis contributes to pain chronification is an open question for future investigation.

**Supplemental Table S1.** (A) Composition of standard diet (SD) and modified diets (MD) used in the present study. Data are expressed as the percentage of each component’s weight relative to the total weight of the diet (% g/g total). (B) Estimated caloric content for each diet, expressed as a percentage of the diet’s total calories, including the contribution of carbohydrates, proteins, and lipids to the total caloric content. The daily intake for SD and MD-1(g/100 g body weight) is also provided (*n* = 4). Abbreviations: ND, Not Determined.

| Ingredient | SD | MD-1 | MD-2 | MD-3 | MD-4 | MD-5 |
| --- | --- | --- | --- | --- | --- | --- |
| Ala | 1.20 | 2.80 | 2.00 | 1.60 | 2.80 | 1.20 |
| Thr | 0.60 | 1.76 | 1.18 | 0.89 | 1.76 | 0.60 |
| Pro | 1.90 | 3.84 | 2.87 | 2.39 | 3.84 | 1.90 |
| Ser | 0.90 | 1.68 | 1.29 | 1.10 | 1.68 | 0.90 |
| Leu | 2.30 | 4.85 | 3.58 | 2.94 | 4.85 | 2.30 |
| Ile | 0.70 | 2.98 | 1.84 | 1.27 | 2.98 | 0.70 |
| Val | 0.90 | 2.98 | 1.94 | 1.42 | 2.98 | 0.90 |
| Phe | 1.00 | 2.26 | 1.63 | 1.32 | 2.26 | 1.00 |
| Tyr | 0.50 | 1.28 | 0.89 | 0.70 | 1.28 | 0.50 |
| Met | 0.50 | 0.98 | 0.74 | 0.62 | 0.98 | 0.50 |
| Cys | 0.30 | 0.68 | 0.49 | 0.40 | 0.68 | 0.30 |
| Trp | 0.20 | 0.76 | 0.48 | 0.34 | 0.76 | 0.20 |
| Oleic acid | 1.10 | 6.06 | 3.58 | 2.34 | 6.06 | 1.10 |
| Erucic acid | 0.00 | 0.25 | 0.13 | 0.06 | 0.25 | 0.00 |
| PEA | 0.00 | 0.08 | 0.04 | 0.02 | 0.00 | 0.08 |

**B.**
|  | Carbohydrate | Fat | Protein | Energy density | Daily intake (g/100g) |
| --- | --- | --- | --- | --- | --- |
| SD | 47.0 | 6.5 | 19.1 | 3.2 | 13.5 ± 0.8 |
| MD-1 | 35.7 | 10.2 | 33.0 | 3.7 | 11.8 ± 1.9 |
| MD-2 | 41.4 | 9.1 | 28.4 | 3.6 | ND |
| MD-3 | 44.2 | 7.8 | 23.7 | 3.4 | ND |
| MD-4 | 35.7 | 10.2 | 33.0 | 3.7 | ND |
| MD-5 | 47.0 | 6.6 | 19.1 | 3.2 | ND |

**Supplemental Table S2.** Serum metabolites influenced by MD-1 in vehicle-injected mice, ranked in ascending order of Log2 fold change (FC). Data are expressed as mean±SEM values (ion counts). Abbreviations: DG, diacylglycerol; PI, phosphatidylinositol; PG, phosphatidylglycerol; PE, phosphatidylethanolamine; PC, phosphatidylcholine; Leu, Leucine; Asp, aspartic acid. FC: fold changes ([Veh_MD-1]/[Veh_SD]).

| Compound Id | Veh_SD |  | Veh_MD-1 |  | Log<sub>2</sub> (FC) | P-value |
| --- | --- | --- | --- | --- | --- | --- |
|  | Mean | SEM | Mean | SEM |  |  |
| DG(18:1(11Z)/20:5(5Z,8Z,11Z,14Z,17Z)/0:0) | 2.50E+04 | 6.79E+03 | 8.09E+03 | 1.65E+03 | -1.63 | 3.63E-02 |
| 5-Nonadecyl-1,3-benzenediol | 2.56E+05 | 4.86E+04 | 9.10E+04 | 1.82E+04 | -1.49 | 8.48E-03 |
| Isocaucalol isomer | 1.17E+06 | 2.95E+05 | 4.27E+05 | 7.37E+04 | -1.45 | 3.46E-02 |
| (S)-Homostachydrine | 1.29E+06 | 1.10E+05 | 4.87E+05 | 4.26E+04 | -1.41 | 2.29E-05 |
| Perillic acid | 1.14E+06 | 1.61E+05 | 4.45E+05 | 7.99E+04 | -1.35 | 2.06E-03 |
| Palmitoylcarnitine | 1.29E+06 | 1.50E+05 | 5.10E+05 | 5.39E+04 | -1.34 | 4.40E-04 |
| 9-Decenoylcarnitine | 1.18E+04 | 1.85E+03 | 5.13E+03 | 3.62E+02 | -1.20 | 5.62E-03 |
| Indole-3-methyl acetate | 3.36E+05 | 7.15E+04 | 1.51E+05 | 1.63E+04 | -1.16 | 3.03E-02 |
| (5Z,8Z)-Tetradecadienoylcarnitine | 1.06E+05 | 1.67E+04 | 4.76E+04 | 4.65E+03 | -1.15 | 7.16E-03 |
| 3',4',5'-Trimethoxycinnamyl alcohol acetate | 6.00E+05 | 6.64E+04 | 2.78E+05 | 3.94E+04 | -1.11 | 9.66E-04 |
| Myristoylcarnitine | 2.39E+05 | 3.02E+04 | 1.13E+05 | 1.75E+04 | -1.08 | 2.90E-03 |
| Isolinderanolide | 4.65E+05 | 7.74E+04 | 2.19E+05 | 1.67E+04 | -1.08 | 1.16E-02 |
| DL-2-Aminooctanoic acid | 1.17E+05 | 1.43E+04 | 5.55E+04 | 5.35E+03 | -1.07 | 1.95E-03 |
| PI(16:0/18:2(9Z,12Z)) | 1.55E+05 | 2.50E+04 | 7.36E+04 | 1.05E+04 | -1.07 | 1.15E-02 |
| 3-Oxododecanoic acid | 1.66E+06 | 2.75E+05 | 7.98E+05 | 1.11E+05 | -1.05 | 1.37E-02 |
| Linoleic acid | 6.64E+08 | 1.28E+08 | 3.26E+08 | 3.54E+07 | -1.03 | 2.84E-02 |
| Tetradecanedioic acid | 4.21E+04 | 6.59E+03 | 2.08E+04 | 2.62E+03 | -1.02 | 1.13E-02 |
| (2R,3R,4R)-2-Amino-4-hydroxy-3-methylpentanoic acid | 3.79E+05 | 5.56E+04 | 1.91E+05 | 2.52E+04 | -0.99 | 9.27E-03 |
| (10Z,12Z)-octadeca-10,12-dienoylcarnitine | 2.90E+05 | 4.10E+04 | 1.47E+05 | 1.45E+04 | -0.98 | 7.20E-03 |
| Podecdysone b isomer | 4.62E+05 | 6.58E+04 | 2.38E+05 | 3.77E+04 | -0.95 | 1.08E-02 |
| Stearoylcarnitine | 1.45E+05 | 1.25E+04 | 7.68E+04 | 9.41E+03 | -0.92 | 5.50E-04 |
| SM(d16:1/24:1(15Z)) | 7.62E+05 | 1.13E+05 | 4.05E+05 | 6.65E+04 | -0.91 | 1.68E-02 |
| 2-Cyclohexen-1-one, 3-(15-hydroxypentadecyl)-2,4,4-trimethyl- | 3.23E+04 | 5.56E+03 | 1.72E+04 | 2.44E+03 | -0.91 | 2.81E-02 |
| PG(16:0/18:2(9Z,12Z)) | 3.87E+04 | 5.38E+03 | 2.06E+04 | 2.38E+03 | -0.91 | 9.65E-03 |
| N2-Acetylornithine | 6.95E+03 | 1.13E+03 | 3.73E+03 | 4.20E+02 | -0.90 | 2.16E-02 |
| DL-Stachydrine | 2.02E+07 | 1.89E+06 | 1.09E+07 | 7.67E+05 | -0.89 | 6.85E-04 |
| Proline betaine | 2.20E+06 | 1.46E+05 | 1.23E+06 | 6.22E+04 | -0.84 | 5.31E-05 |
| 5-Dodecenoylcarnitine | 4.75E+04 | 7.27E+03 | 2.68E+04 | 4.01E+03 | -0.83 | 2.59E-02 |
| Octadeca-2,4,6,8-tetraenoic acid | 2.05E+05 | 3.71E+04 | 1.19E+05 | 1.00E+04 | -0.79 | 4.80E-02 |
| PE(P-18:0/22:4(7Z,10Z,13Z,16Z)) | 3.35E+04 | 5.74E+03 | 1.94E+04 | 3.07E+03 | -0.79 | 4.94E-02 |
| PE(20:2(11Z,14Z)/15:0) | 3.88E+05 | 3.22E+04 | 2.31E+05 | 1.93E+04 | -0.75 | 9.10E-04 |
| Adrenic acid | 9.05E+06 | 1.51E+06 | 5.46E+06 | 3.98E+05 | -0.73 | 4.39E-02 |
| Tetracosahexaenoic acid | 1.66E+06 | 2.17E+05 | 1.03E+06 | 1.37E+05 | -0.69 | 2.74E-02 |
| 4-Guanidinobutanoic acid | 2.32E+05 | 2.88E+04 | 1.46E+05 | 2.04E+04 | -0.68 | 2.68E-02 |
| 9-Hexadecenoylcarnitine | 1.85E+05 | 2.89E+04 | 1.16E+05 | 1.29E+04 | -0.67 | 4.96E-02 |
| Ferulic acid 4-O-sulfate | 1.83E+06 | 1.75E+05 | 1.18E+06 | 1.57E+05 | -0.63 | 1.51E-02 |
| 2-Methoxyestradiol-3-methylether | 2.31E+04 | 2.47E+03 | 1.54E+04 | 1.80E+03 | -0.58 | 2.44E-02 |
| PG(18:0/16:0) | 4.60E+04 | 5.31E+03 | 3.14E+04 | 3.48E+03 | -0.55 | 3.71E-02 |
| PC(18:2(9Z,12Z)/14:0) | 1.98E+06 | 2.10E+05 | 1.41E+06 | 8.64E+04 | -0.49 | 2.83E-02 |
| Argininosuccinic acid | 4.79E+03 | 5.09E+02 | 3.47E+03 | 3.42E+02 | -0.46 | 4.96E-02 |
| L-Carnitine | 2.22E+08 | 1.40E+07 | 1.61E+08 | 1.06E+07 | -0.46 | 3.52E-03 |
| PC(18:2(9Z,12Z)/16:1(9Z)) | 4.59E+06 | 4.18E+05 | 3.54E+06 | 2.36E+05 | -0.37 | 4.74E-02 |
| Indoleacrylic acid | 3.01E+07 | 2.46E+06 | 3.69E+07 | 1.98E+06 | 0.29 | 4.91E-02 |
| 8-Hydroxyquinoline | 2.74E+06 | 2.64E+05 | 3.44E+06 | 1.99E+05 | 0.33 | 4.84E-02 |
| 1,5-Naphthalenediamine | 2.02E+06 | 1.84E+05 | 2.61E+06 | 1.30E+05 | 0.37 | 1.89E-02 |
| Methylindole isomer | 5.97E+05 | 5.26E+04 | 8.06E+05 | 5.22E+04 | 0.43 | 1.34E-02 |
| 3-Indoleacetonitrile | 6.23E+05 | 6.05E+04 | 8.62E+05 | 5.16E+04 | 0.47 | 8.84E-03 |
| PC(22:5(4Z,7Z,10Z,13Z,16Z)/16:0) | 1.26E+07 | 1.67E+06 | 1.77E+07 | 9.69E+05 | 0.48 | 2.11E-02 |
| D-(+)-Proline | 2.47E+07 | 1.60E+06 | 3.46E+07 | 2.53E+06 | 0.49 | 7.33E-03 |
| PE(22:2(13Z,16Z)/18:1(11Z)) | 3.62E+05 | 3.49E+04 | 5.22E+05 | 4.13E+04 | 0.53 | 1.09E-02 |
| LysoPC(24:1(15Z)/0:0) | 1.55E+05 | 2.13E+04 | 2.27E+05 | 2.11E+04 | 0.56 | 2.87E-02 |
| PE(18:2(9Z,12Z)/18:1(9Z)) | 1.61E+05 | 2.21E+04 | 2.40E+05 | 2.47E+04 | 0.58 | 3.07E-02 |
| Galactitol | 8.91E+05 | 6.94E+04 | 1.36E+06 | 1.62E+05 | 0.62 | 2.67E-02 |
| Cer(d18:1/24:1(15Z)) | 1.12E+05 | 1.54E+04 | 1.74E+05 | 1.47E+04 | 0.64 | 1.06E-02 |
| Elaidic acid | 5.63E+08 | 9.07E+07 | 8.87E+08 | 1.09E+08 | 0.66 | 3.93E-02 |
| Deoxyadenosine monophosphate | 2.89E+03 | 3.87E+02 | 4.60E+03 | 4.51E+02 | 0.67 | 1.25E-02 |
| 3-Methylcrotonylglycine | 5.48E+04 | 8.34E+03 | 8.80E+04 | 8.73E+03 | 0.68 | 1.54E-02 |
| Docosatrienoic acid | 8.51E+05 | 1.31E+05 | 1.37E+06 | 1.22E+05 | 0.69 | 1.10E-02 |
| PC(18:2(9Z,12Z)/18:1(11Z)) | 3.13E+07 | 4.65E+06 | 5.08E+07 | 2.63E+06 | 0.70 | 2.66E-03 |
| Indole-3-carboxylic acid-sulphate isomer | 7.13E+05 | 1.08E+05 | 1.19E+06 | 1.62E+05 | 0.74 | 3.07E-02 |
| L-Allothreonine | 6.40E+06 | 4.48E+05 | 1.09E+07 | 7.04E+05 | 0.76 | 2.61E-04 |
| LysoPC(P-18:0/0:0) | 3.06E+05 | 3.74E+04 | 5.39E+05 | 4.85E+04 | 0.82 | 2.40E-03 |
| lysops(22:0) | 5.58E+06 | 8.13E+05 | 9.83E+06 | 7.56E+05 | 0.82 | 1.68E-03 |
| LysoPC(18:1(9Z)/0:0) | 5.45E+07 | 7.92E+06 | 9.64E+07 | 1.01E+07 | 0.82 | 6.45E-03 |
| Annoglabasin F | 1.22E+06 | 1.82E+05 | 2.15E+06 | 2.31E+05 | 0.82 | 7.46E-03 |
| 3-Hydroxybutyrylcarnitine | 2.18E+05 | 1.70E+04 | 3.95E+05 | 4.51E+04 | 0.86 | 6.60E-03 |
| PC(18:1(9Z)/15:0) | 2.47E+05 | 2.87E+04 | 4.53E+05 | 4.49E+04 | 0.87 | 2.75E-03 |
| DG(18:2(9Z,12Z)/18:3(6Z,9Z,12Z)/0:0) | 4.21E+04 | 8.44E+03 | 7.72E+04 | 1.29E+04 | 0.88 | 4.36E-02 |
| Indoxyl sulfate | 6.87E+06 | 8.78E+05 | 1.26E+07 | 1.39E+06 | 0.88 | 5.39E-03 |
| LysoPC(22:1(13Z)/0:0) | 3.00E+05 | 4.43E+04 | 5.62E+05 | 5.74E+04 | 0.91 | 3.39E-03 |
| 5-Aminovaleric acid | 5.37E+07 | 4.68E+06 | 1.01E+08 | 1.17E+07 | 0.91 | 5.58E-03 |
| PI(20:4(8Z,11Z,14Z,17Z)/18:1(9Z)) | 8.97E+04 | 1.72E+04 | 1.71E+05 | 2.43E+04 | 0.93 | 1.90E-02 |
| PE(18:0/18:1(9Z)) | 3.06E+05 | 2.98E+04 | 5.95E+05 | 3.31E+04 | 0.96 | 1.62E-05 |
| LysoPE(0:0/18:1(11Z)) | 1.01E+06 | 1.29E+05 | 2.02E+06 | 1.99E+05 | 1.01 | 1.32E-03 |
| 5-Hydroxyindole | 1.15E+05 | 1.32E+04 | 2.31E+05 | 2.58E+04 | 1.01 | 2.90E-03 |
| N-Acetyltryptophan | 3.22E+05 | 1.61E+04 | 6.60E+05 | 6.54E+04 | 1.04 | 1.71E-03 |
| Leu asp isomer | 1.23E+05 | 1.05E+04 | 2.73E+05 | 3.14E+04 | 1.15 | 2.34E-03 |
| Pemoline | 5.26E+03 | 5.88E+02 | 1.29E+04 | 1.15E+03 | 1.29 | 2.19E-04 |
| Tiglylcarnitine | 6.70E+03 | 7.17E+02 | 1.75E+04 | 1.44E+03 | 1.39 | 8.75E-05 |
| PE(18:2(9Z,12Z)/24:1(15Z)) | 4.39E+04 | 8.23E+03 | 1.19E+05 | 2.54E+04 | 1.44 | 2.53E-02 |
| PE(16:0/18:1(9Z)) | 2.64E+04 | 3.78E+03 | 7.42E+04 | 1.44E+04 | 1.49 | 1.54E-02 |
| PC(18:1(9Z)/20:2(11Z,14Z)) | 1.75E+06 | 4.36E+05 | 5.63E+06 | 4.73E+05 | 1.69 | 3.15E-05 |
| PE(16:1(9Z)/22:0) | 2.59E+05 | 2.69E+04 | 8.78E+05 | 7.72E+04 | 1.76 | 9.25E-05 |
| Cresol sulfate isomer | 3.84E+06 | 1.15E+06 | 1.41E+07 | 1.56E+06 | 1.87 | 2.01E-04 |
| Guanosine | 2.44E+04 | 7.51E+03 | 1.07E+05 | 2.73E+04 | 2.14 | 2.26E-02 |
| 1-Pentanesulfenothioic acid | 1.77E+06 | 4.19E+05 | 1.39E+07 | 2.77E+06 | 2.98 | 4.45E-03 |
| Inosine | 1.31E+06 | 3.22E+05 | 1.16E+07 | 2.32E+06 | 3.15 | 4.24E-03 |
| 2-(3,4-Dihydroxybenzoyloxy)-4,6-dihydroxybenzoate | 3.22E+04 | 8.81E+03 | 3.96E+05 | 9.37E+04 | 3.62 | 8.00E-03 |
| PC(18:1(9Z)/16:0) | 7.02E+05 | 2.85E+05 | 1.59E+07 | 2.17E+06 | 4.50 | 3.76E-04 |
| Xanthine | 1.07E+06 | 4.62E+05 | 3.17E+07 | 1.17E+07 | 4.89 | 4.04E-02 |
| PE(18:0/22:1(13Z)) | 4.41E+03 | 9.38E+02 | 2.48E+05 | 3.72E+04 | 5.81 | 6.14E-04 |

**Supplemental Table S3.**
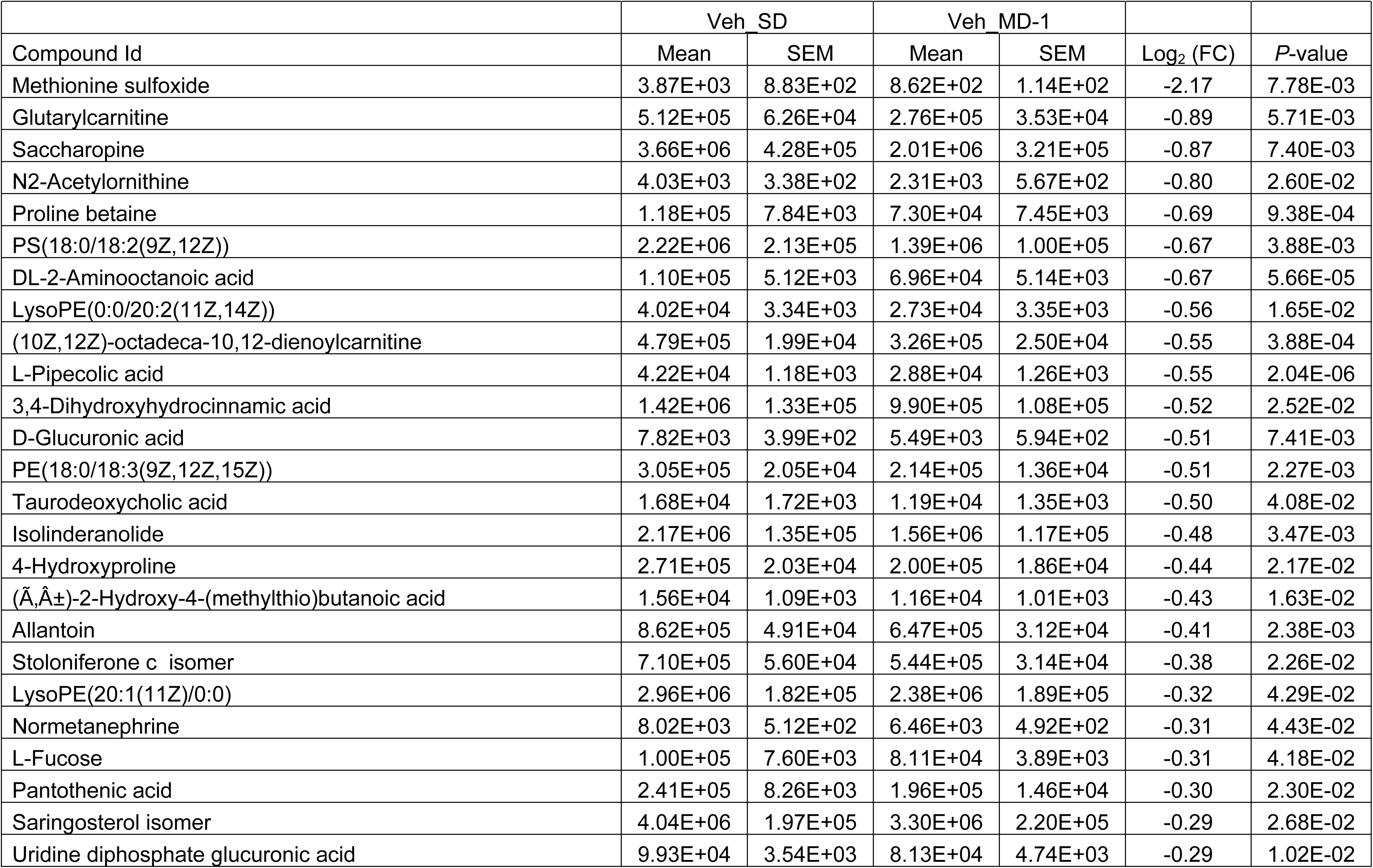
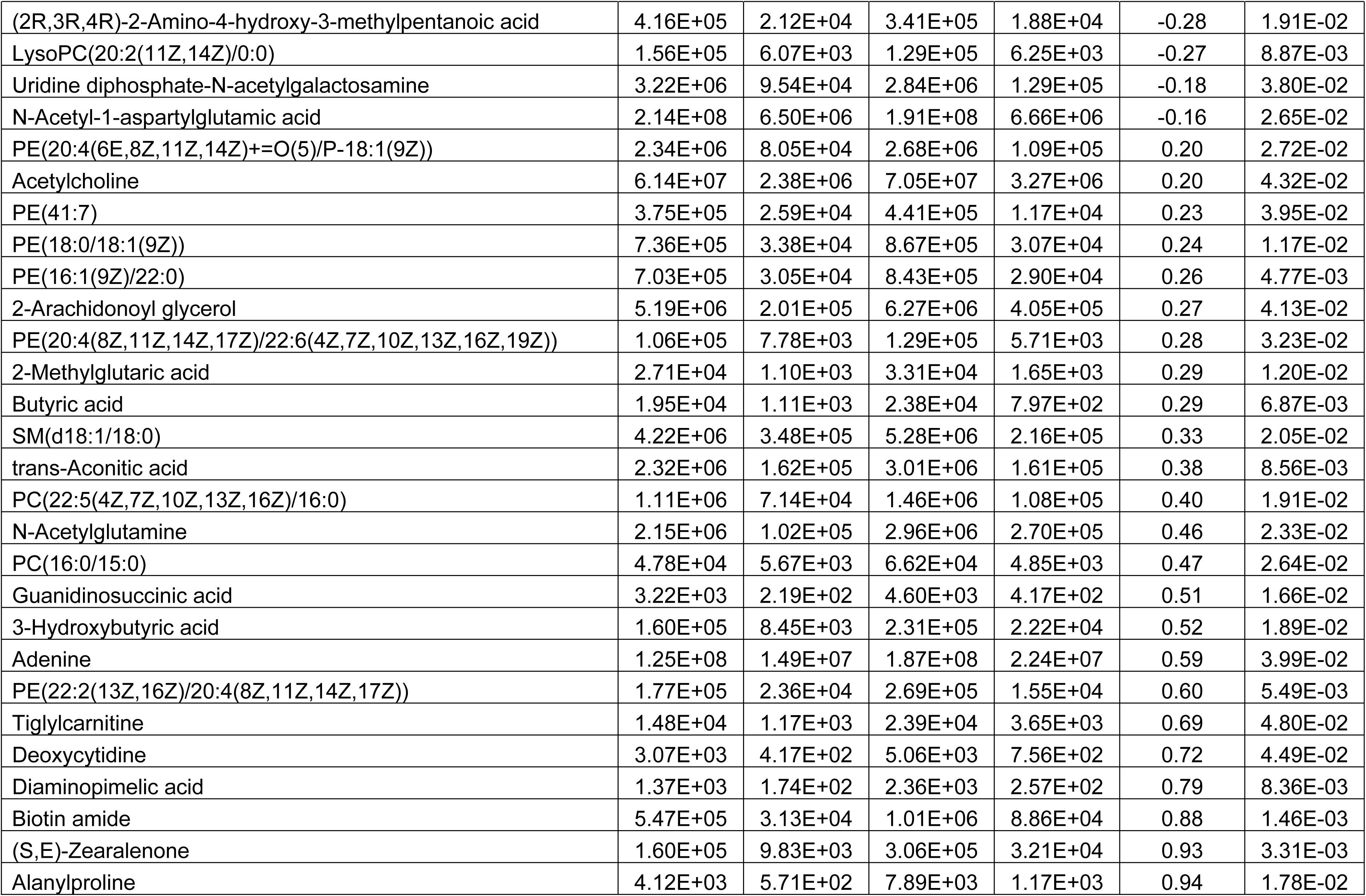
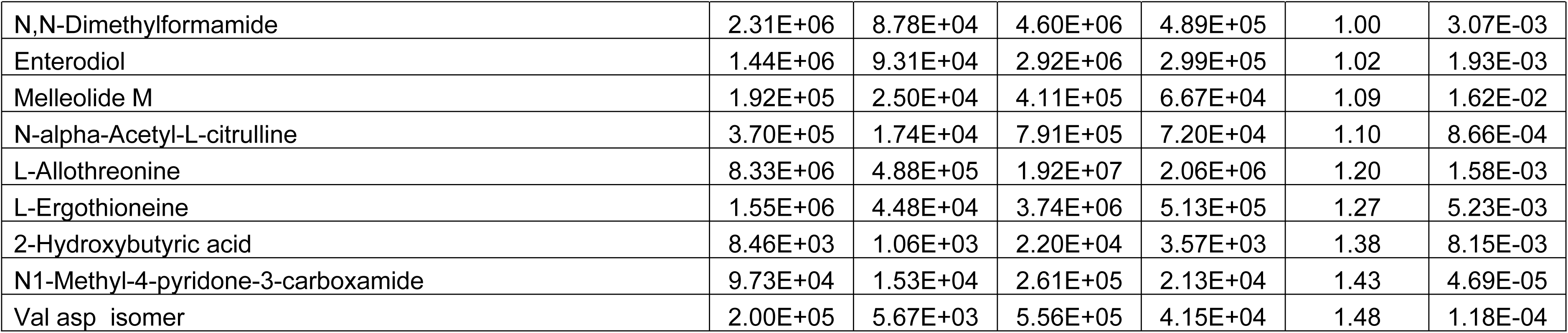
Spinal cord metabolites affected by MD-1 in vehicle-injected mice, ranked in ascending order of Log_2_ fold change (FC). Data are expressed as mean±SEM values (ion counts). Abbreviations: PS, phosphatidylserine; Val, valine; Asp, aspartate; FC: fold changes ([Veh_MD-1]/[Veh_SD]).

**Supplemental Table S4.**
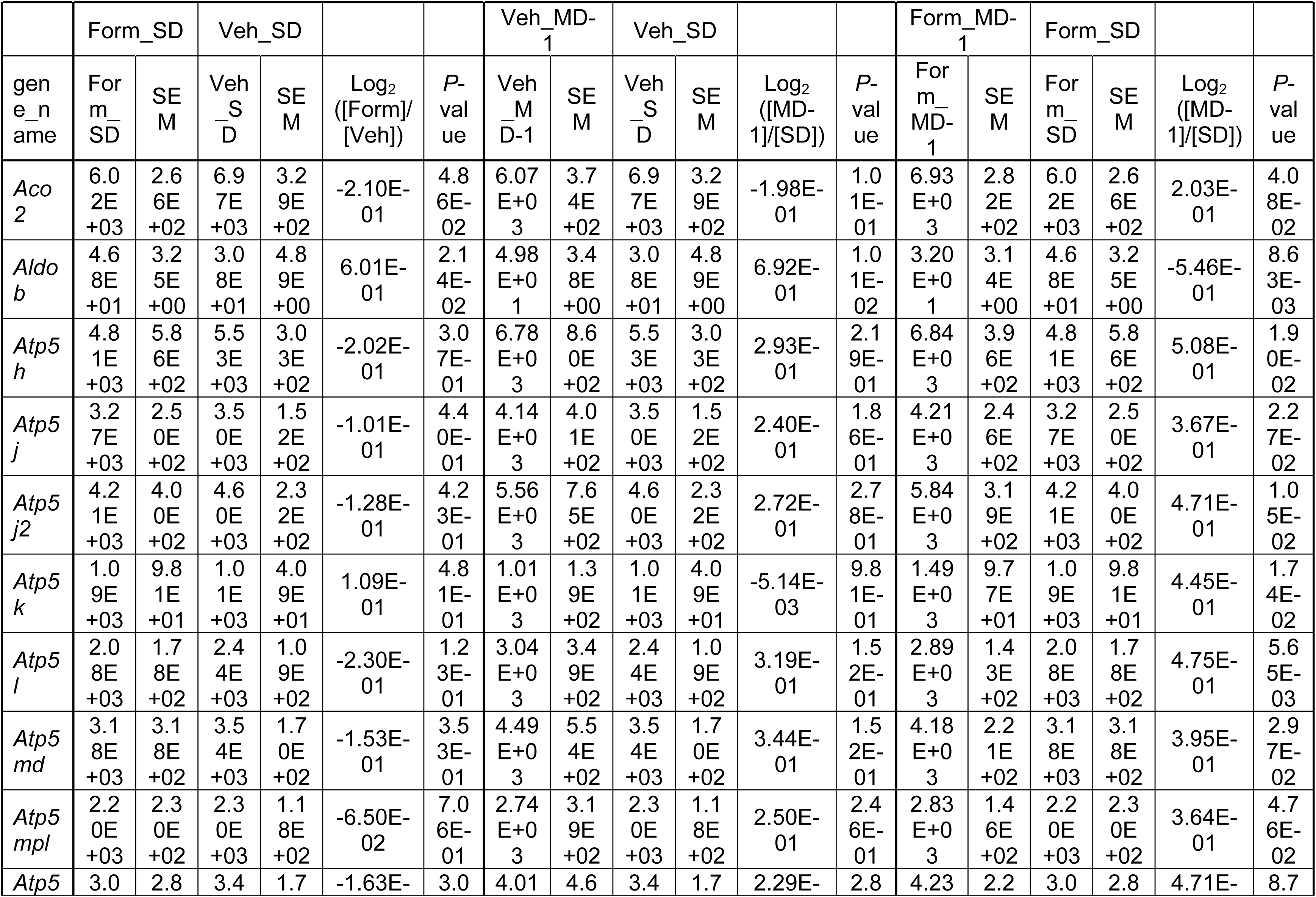
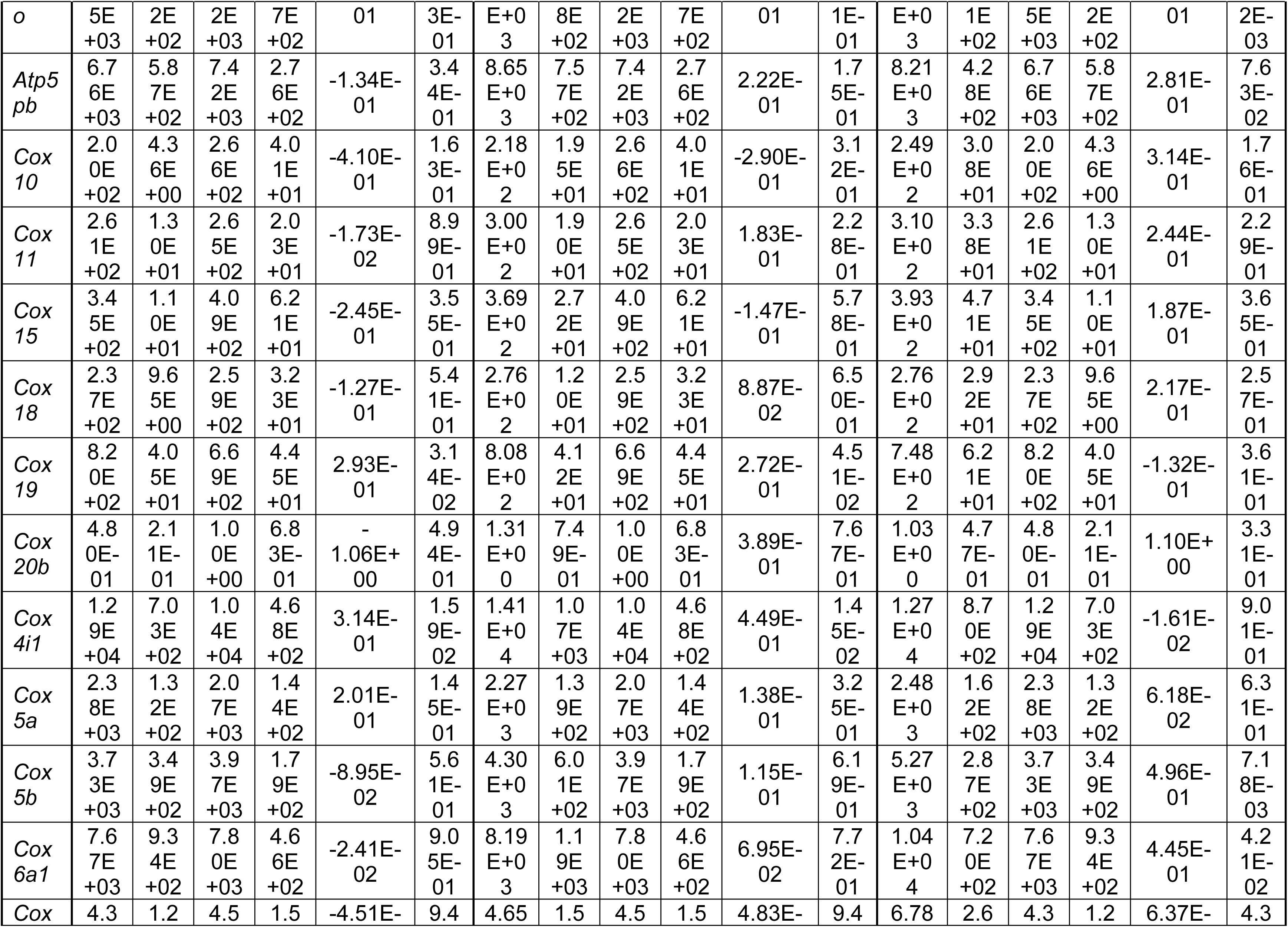
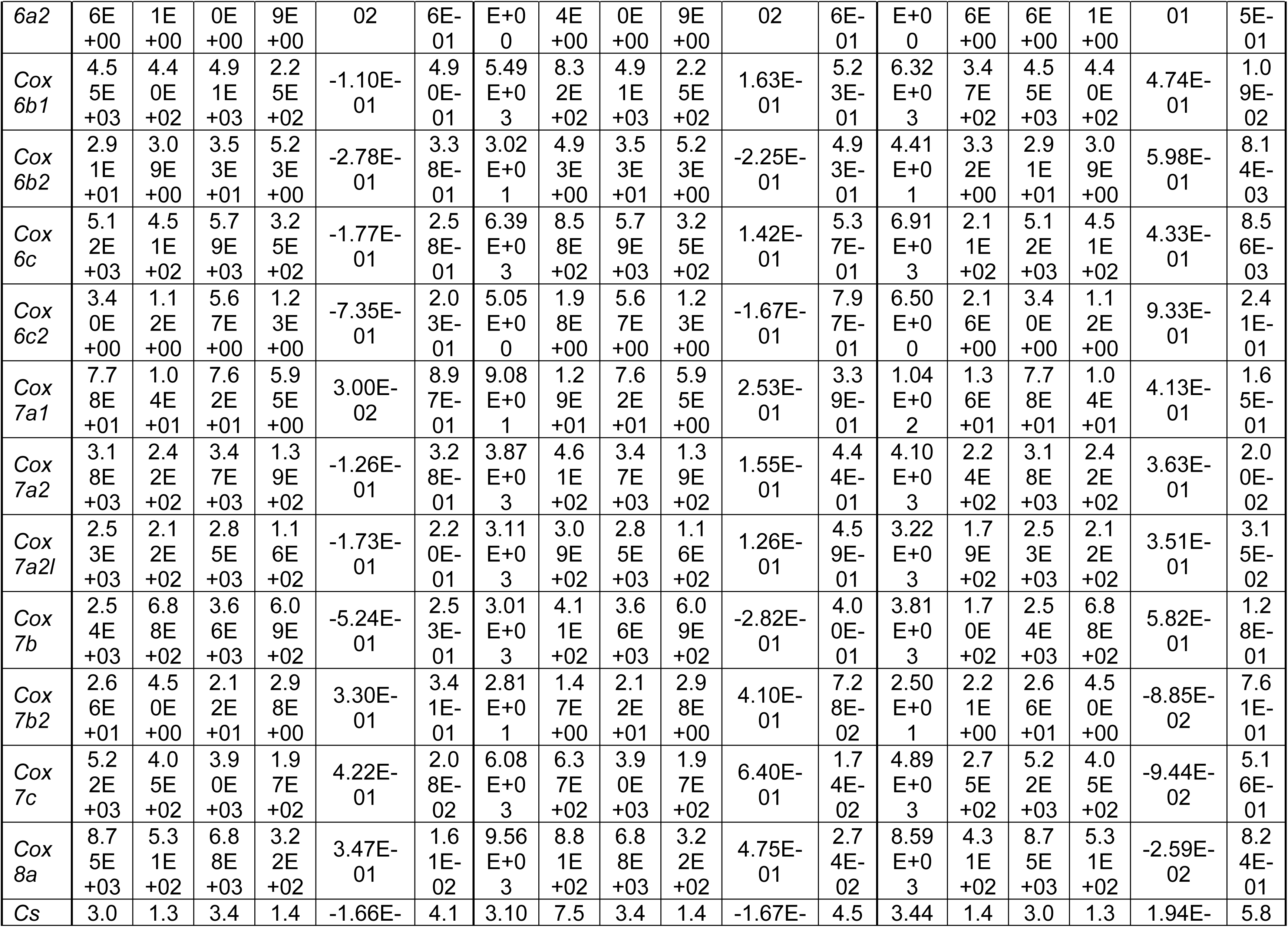
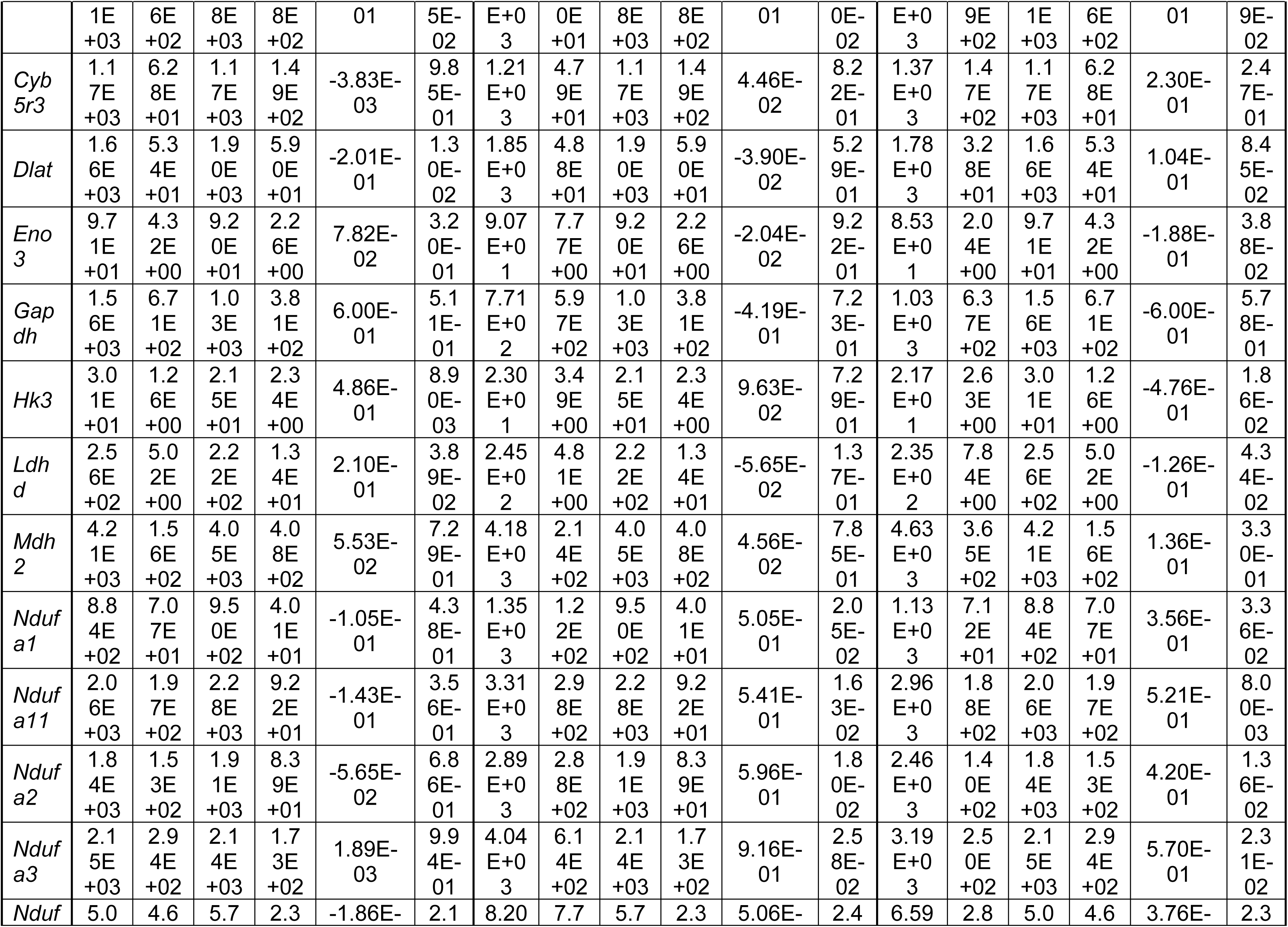
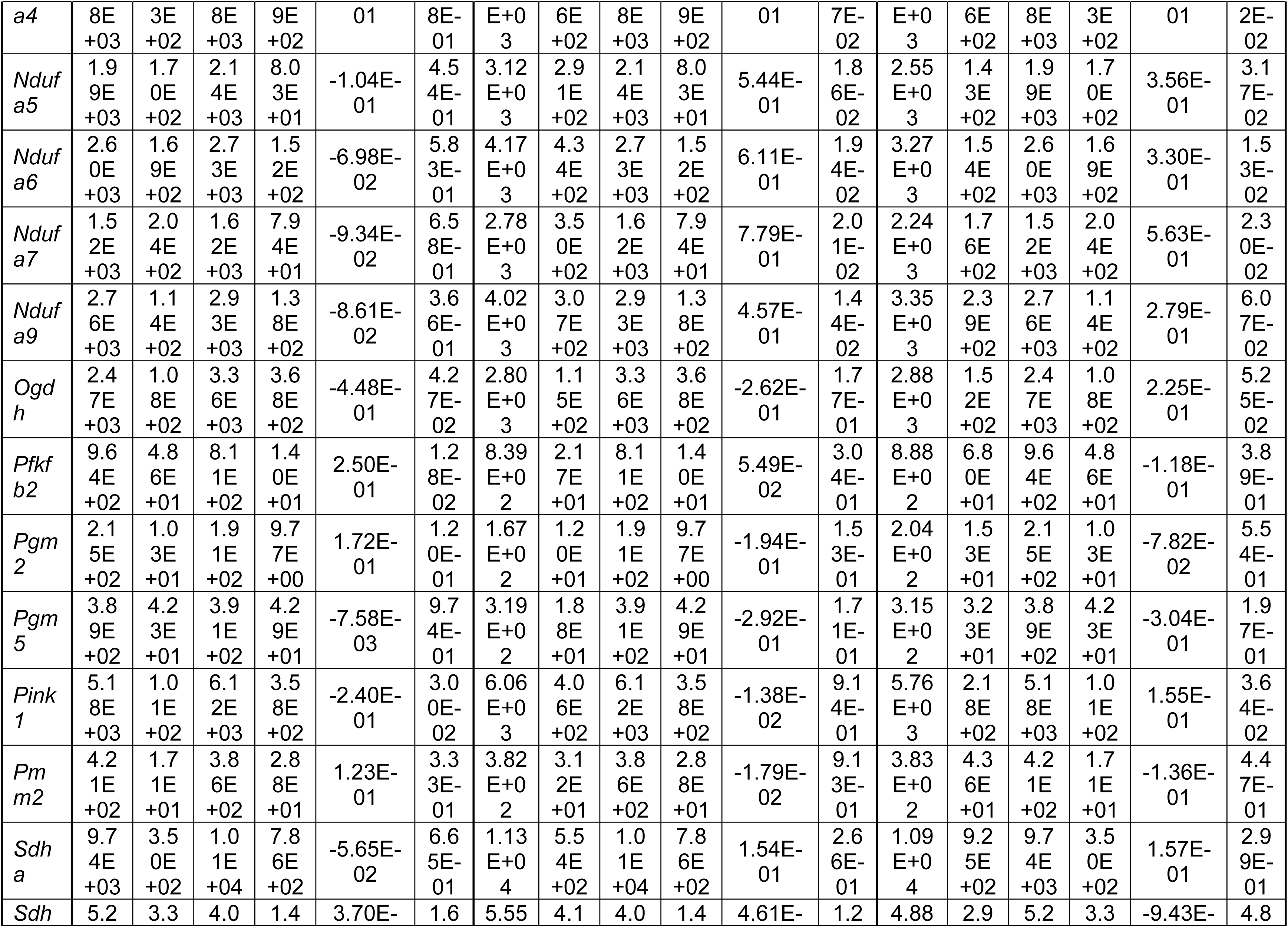
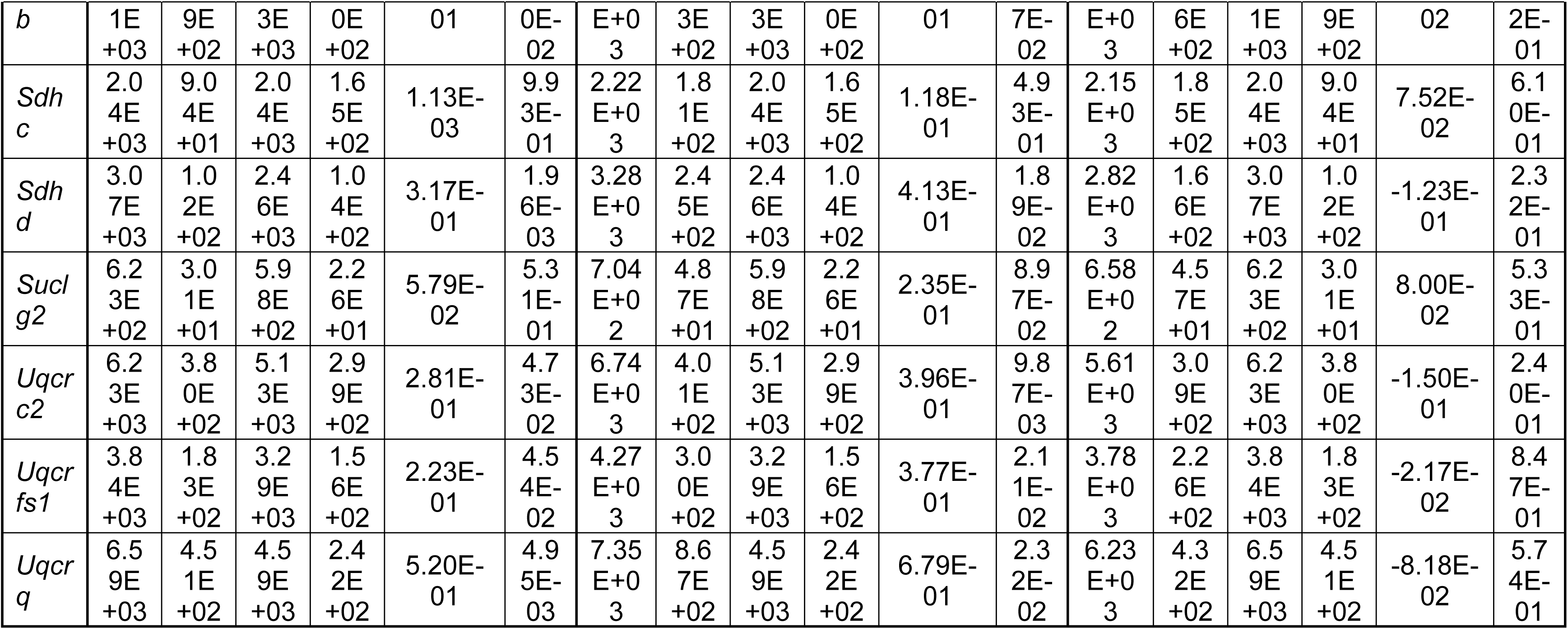
Gene transcription levels associated with glycolysis, Krebs’ cycle, and oxidative phosphorylation in ipsilateral L4-L6 spinal cord of vehicle (Veh)- or formalin (Form)-injected mice fed either SD or MD-1. Genes are listed in alphabetical order. Data are expressed as mean±SEM values (counts). FC: fold change.

**Supplemental Table S5.**
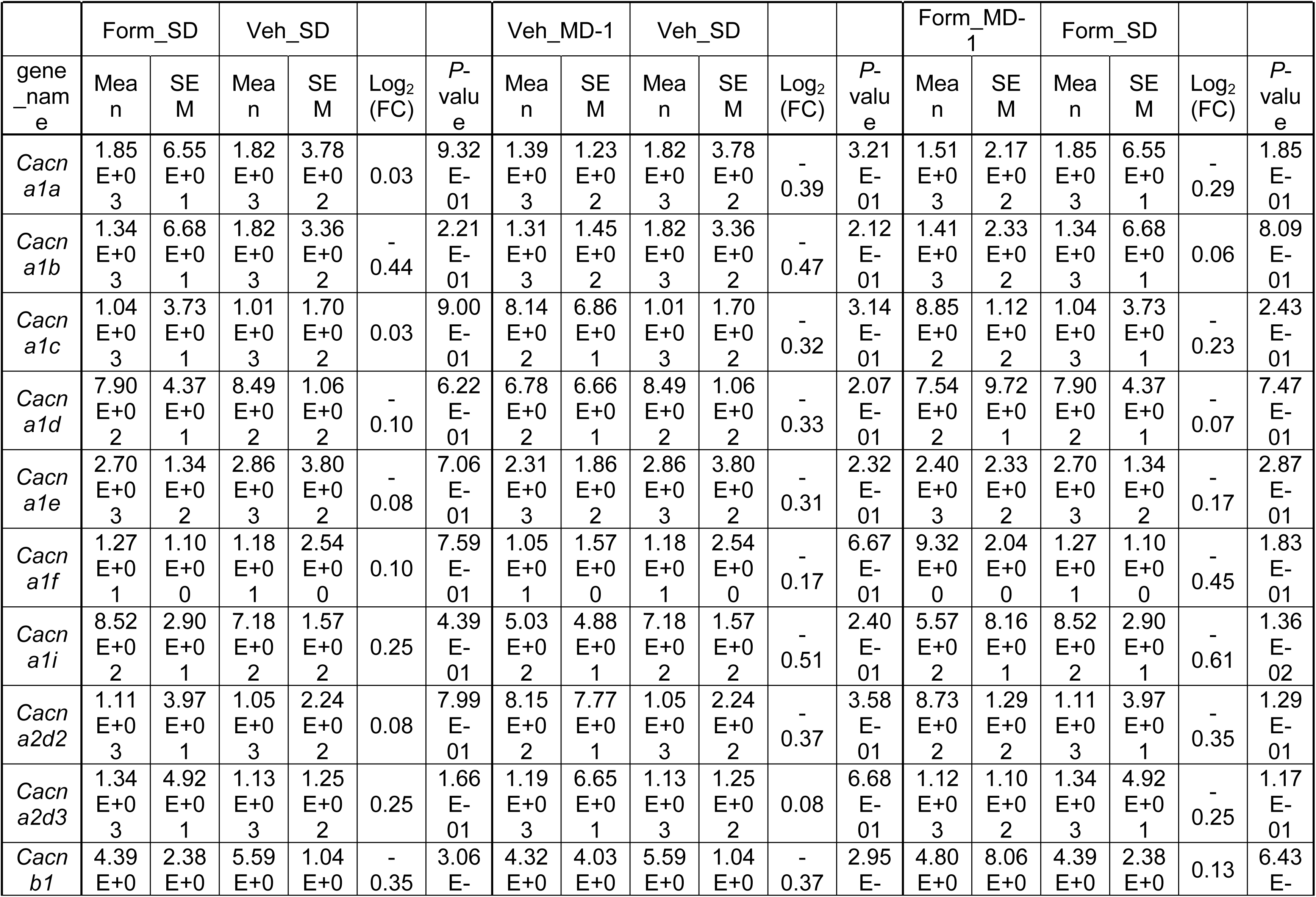
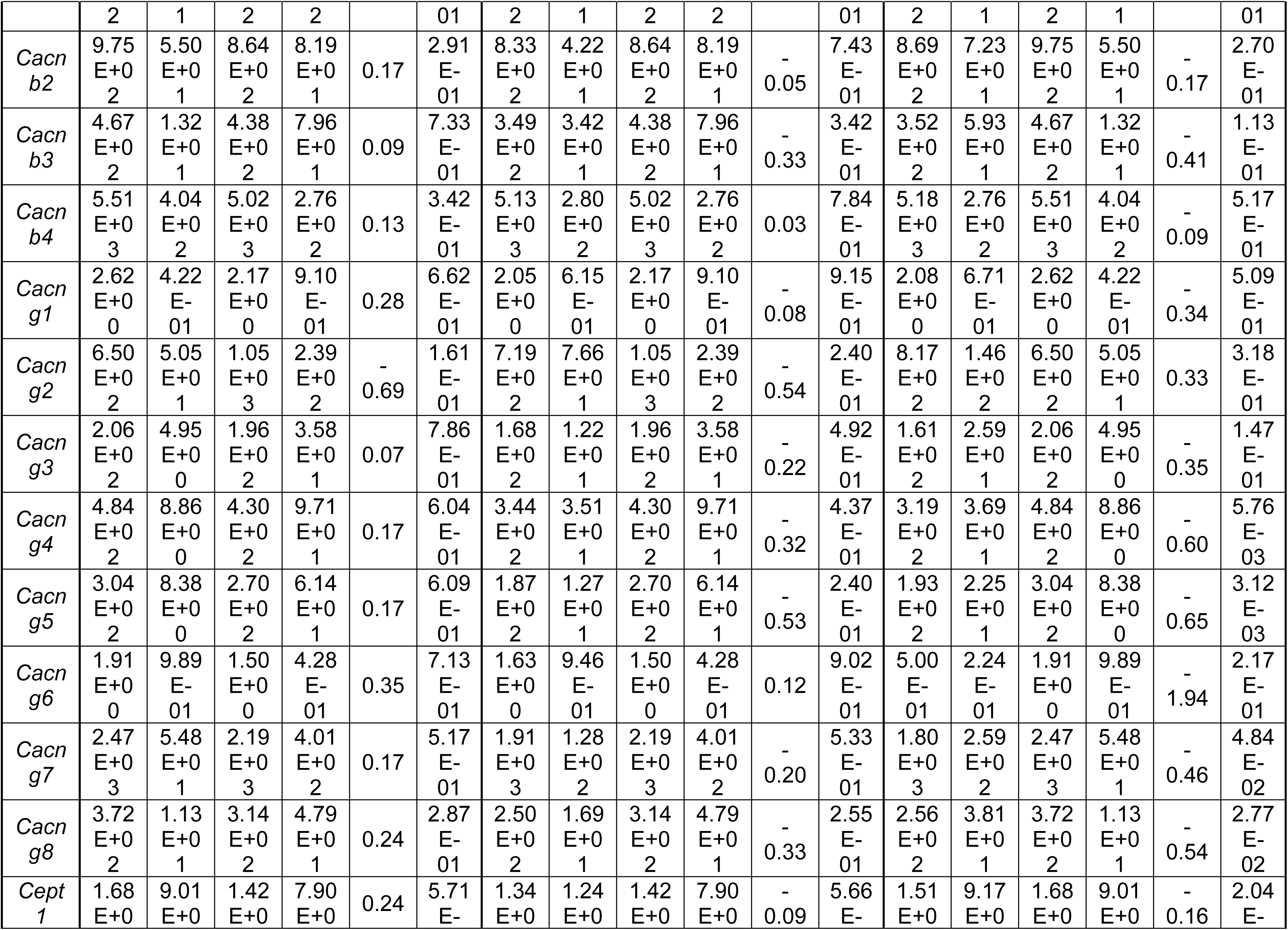
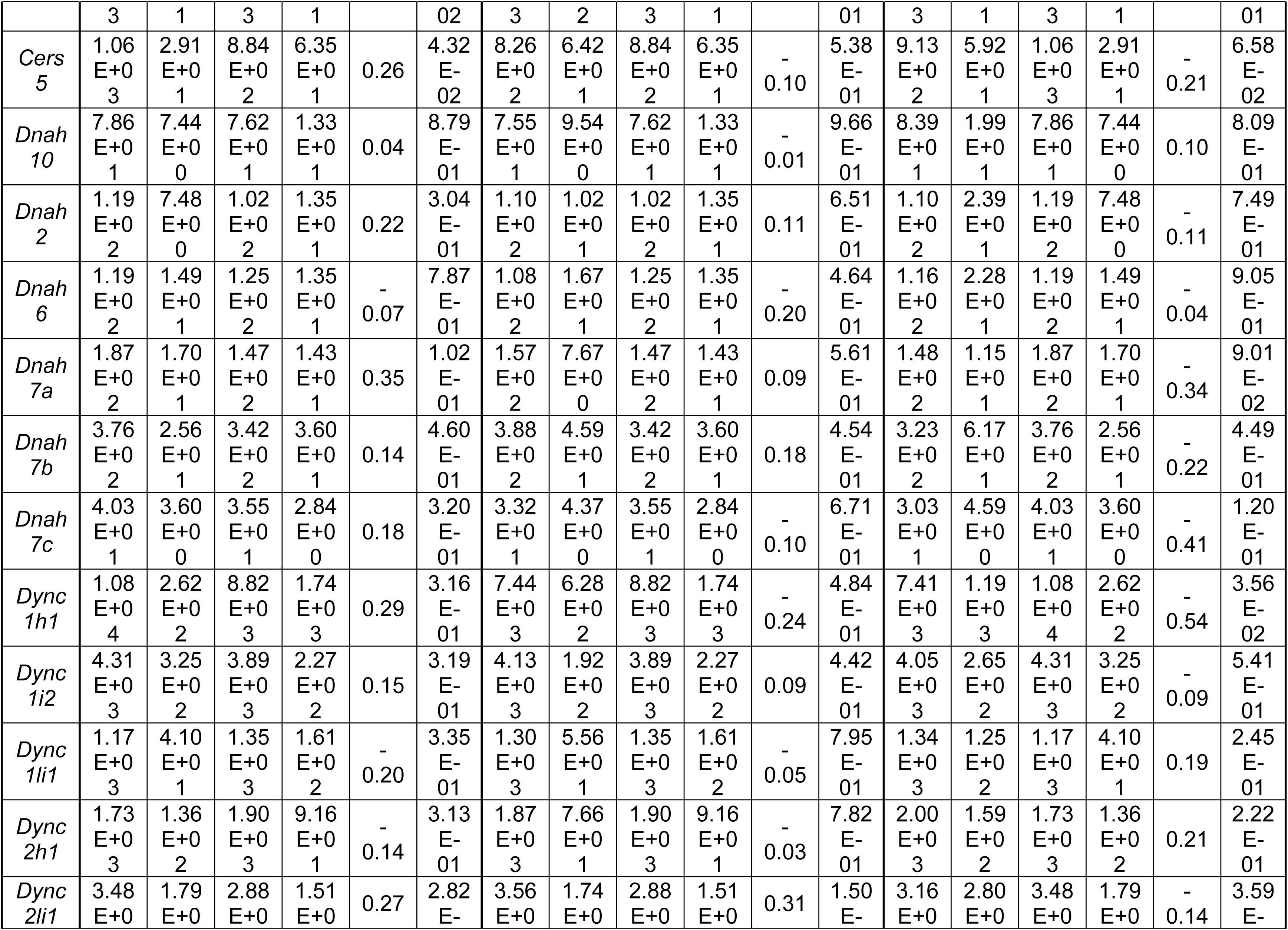
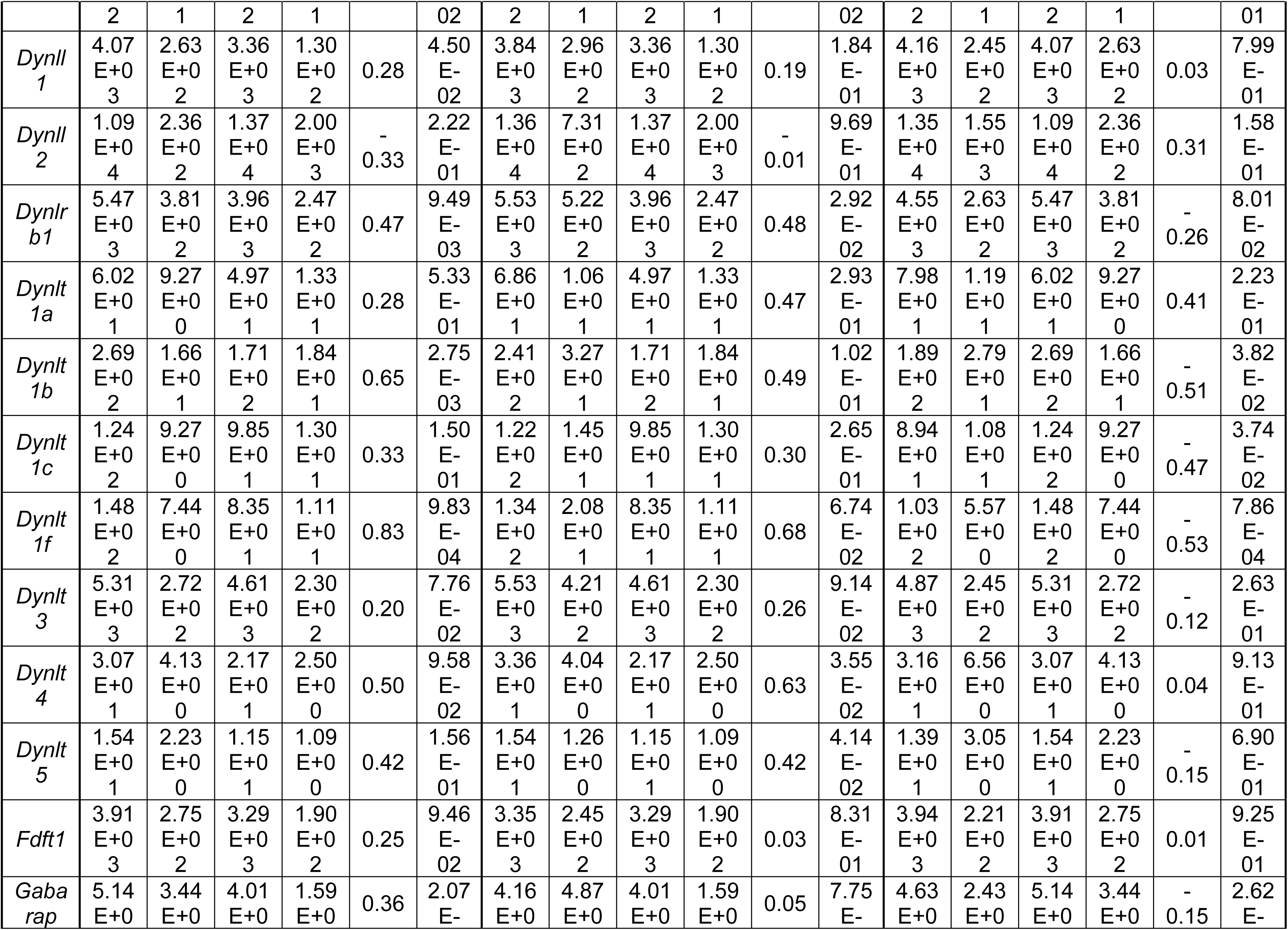
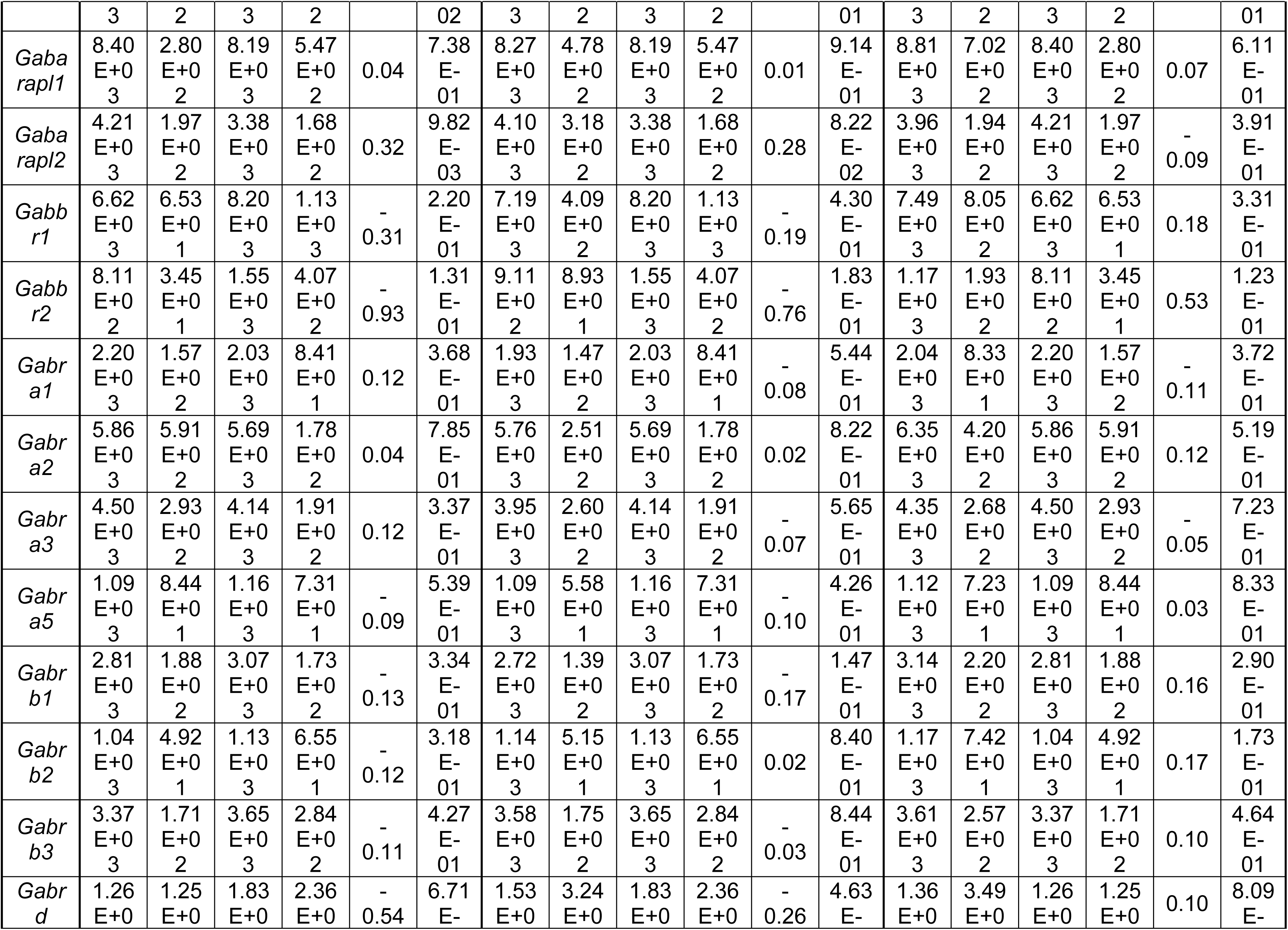
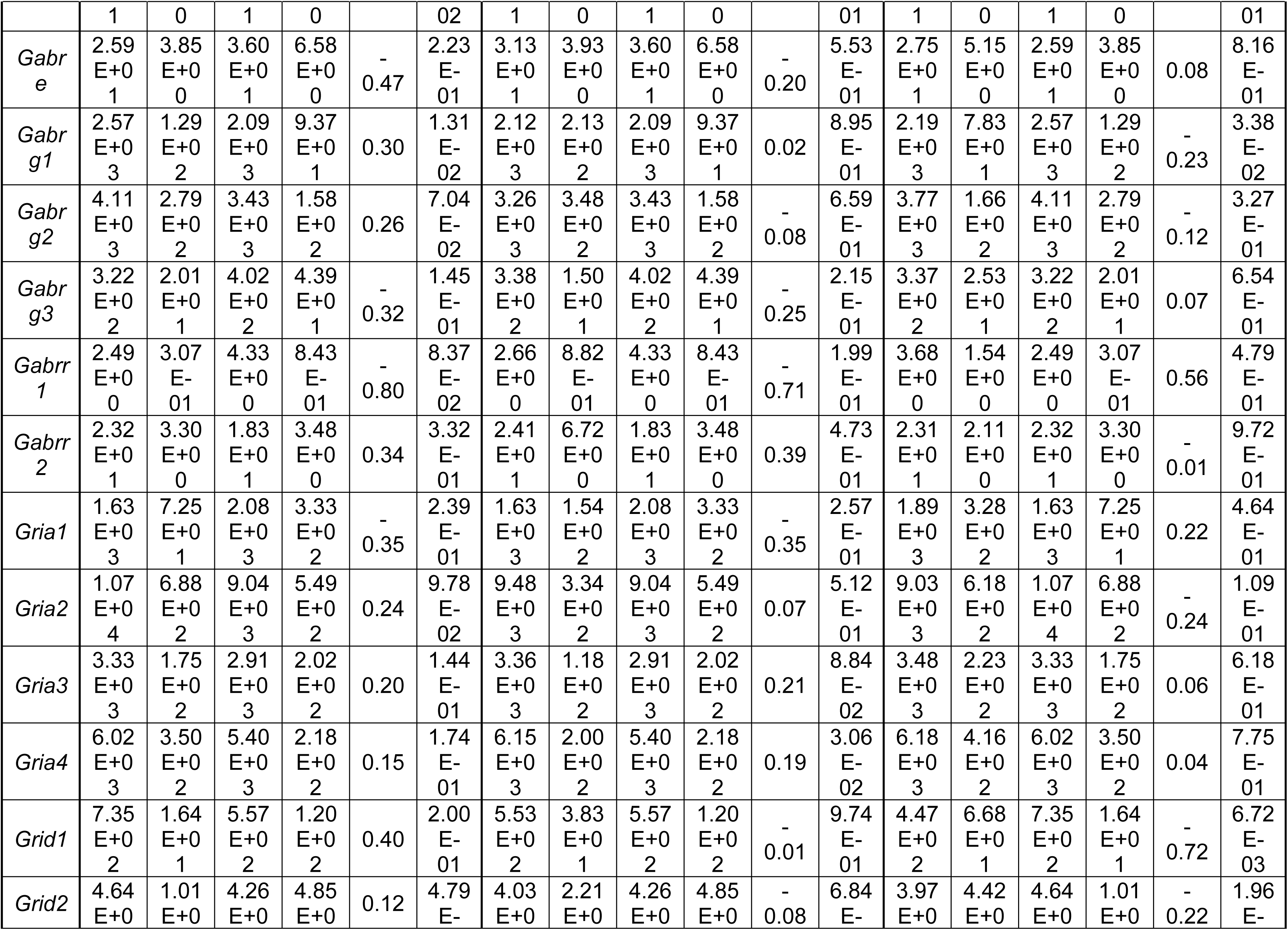
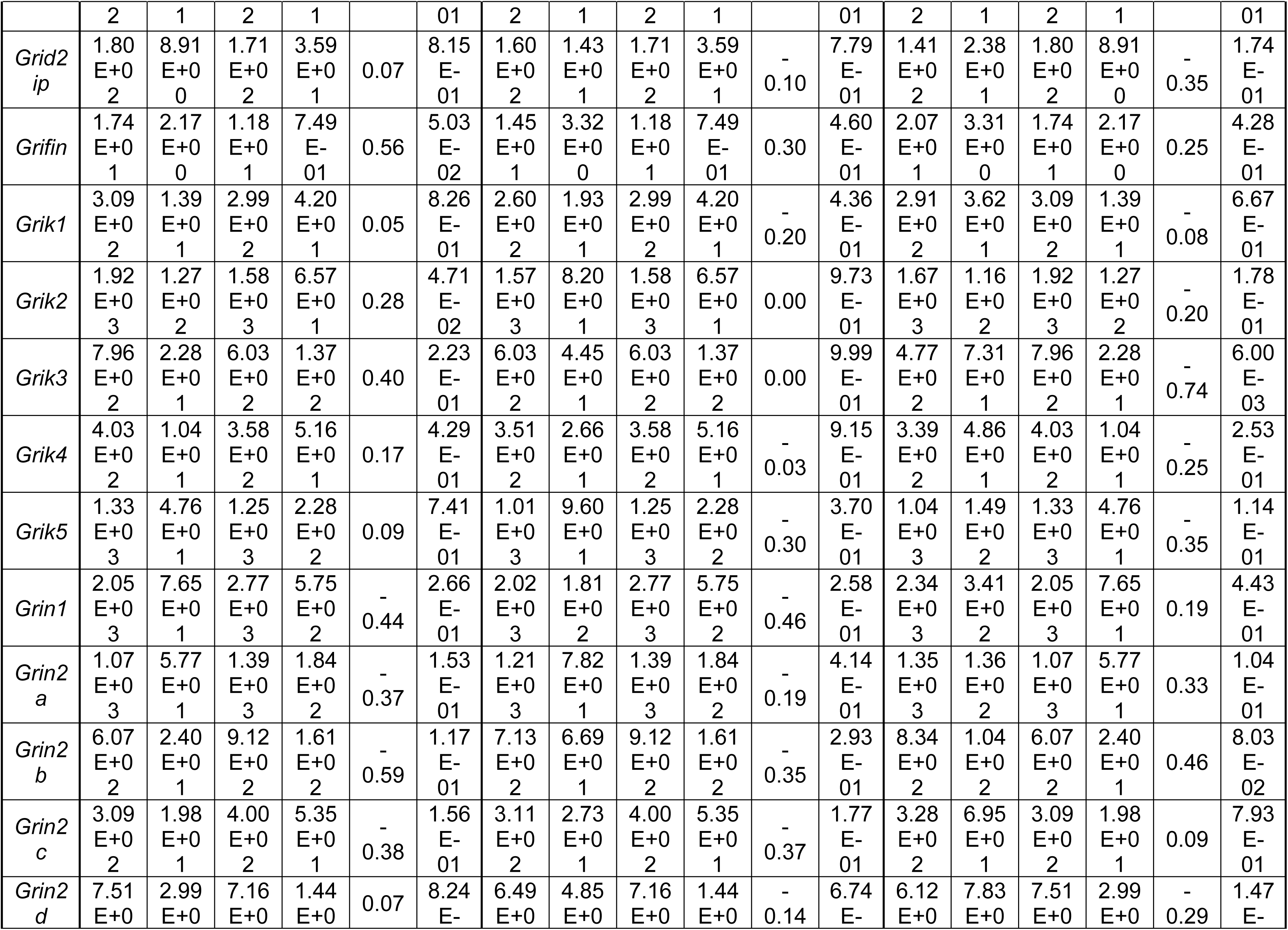
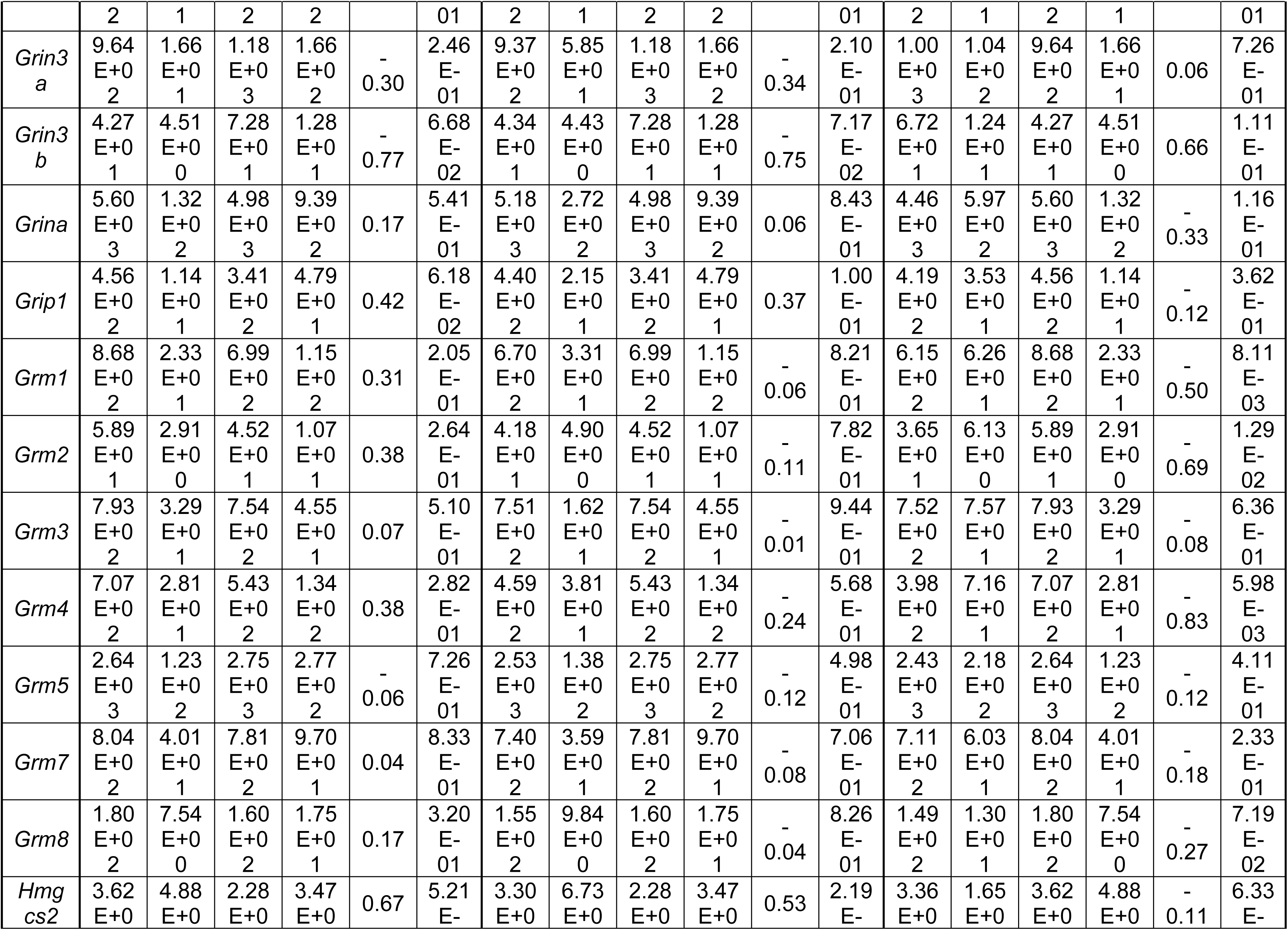
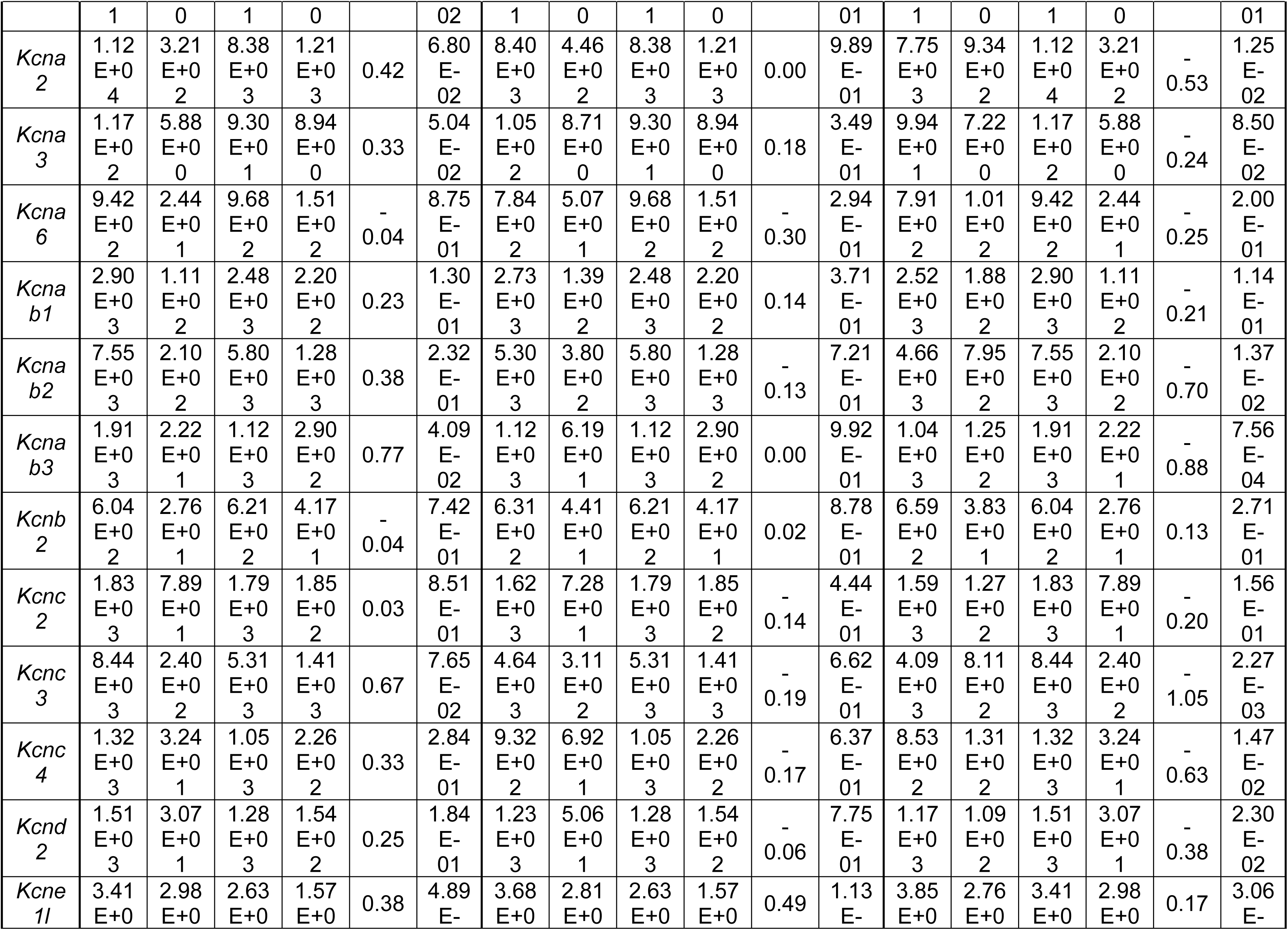
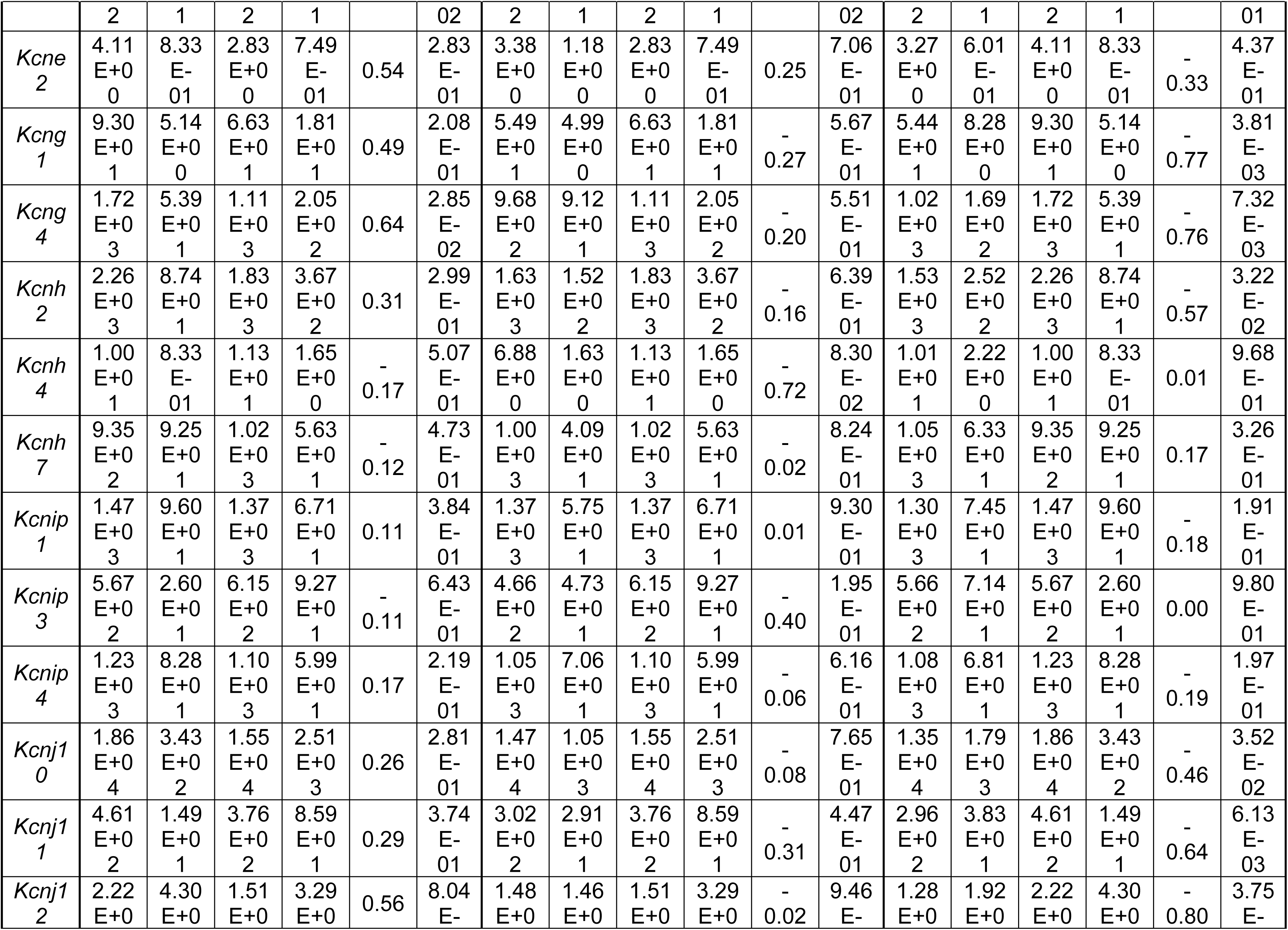
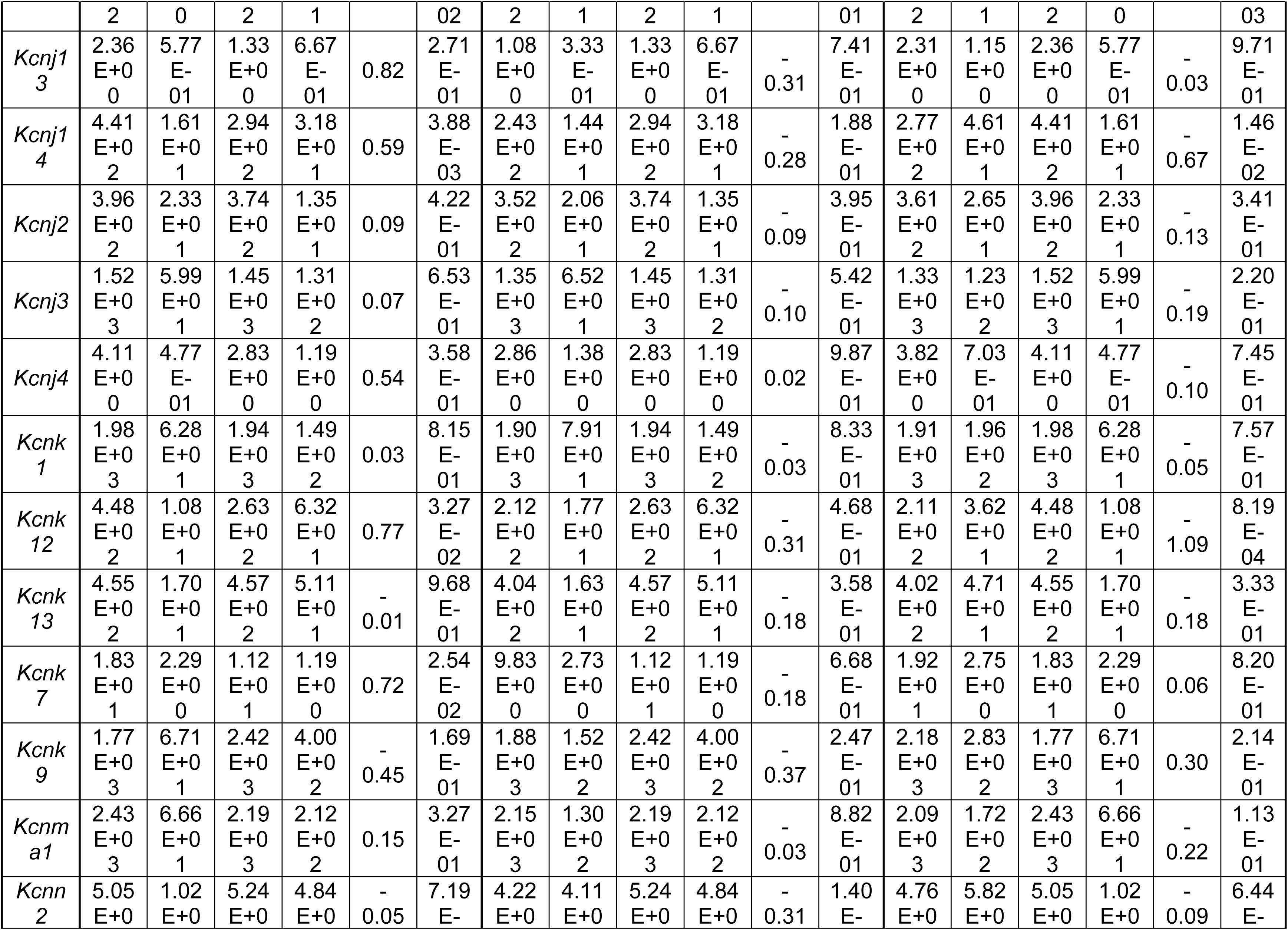
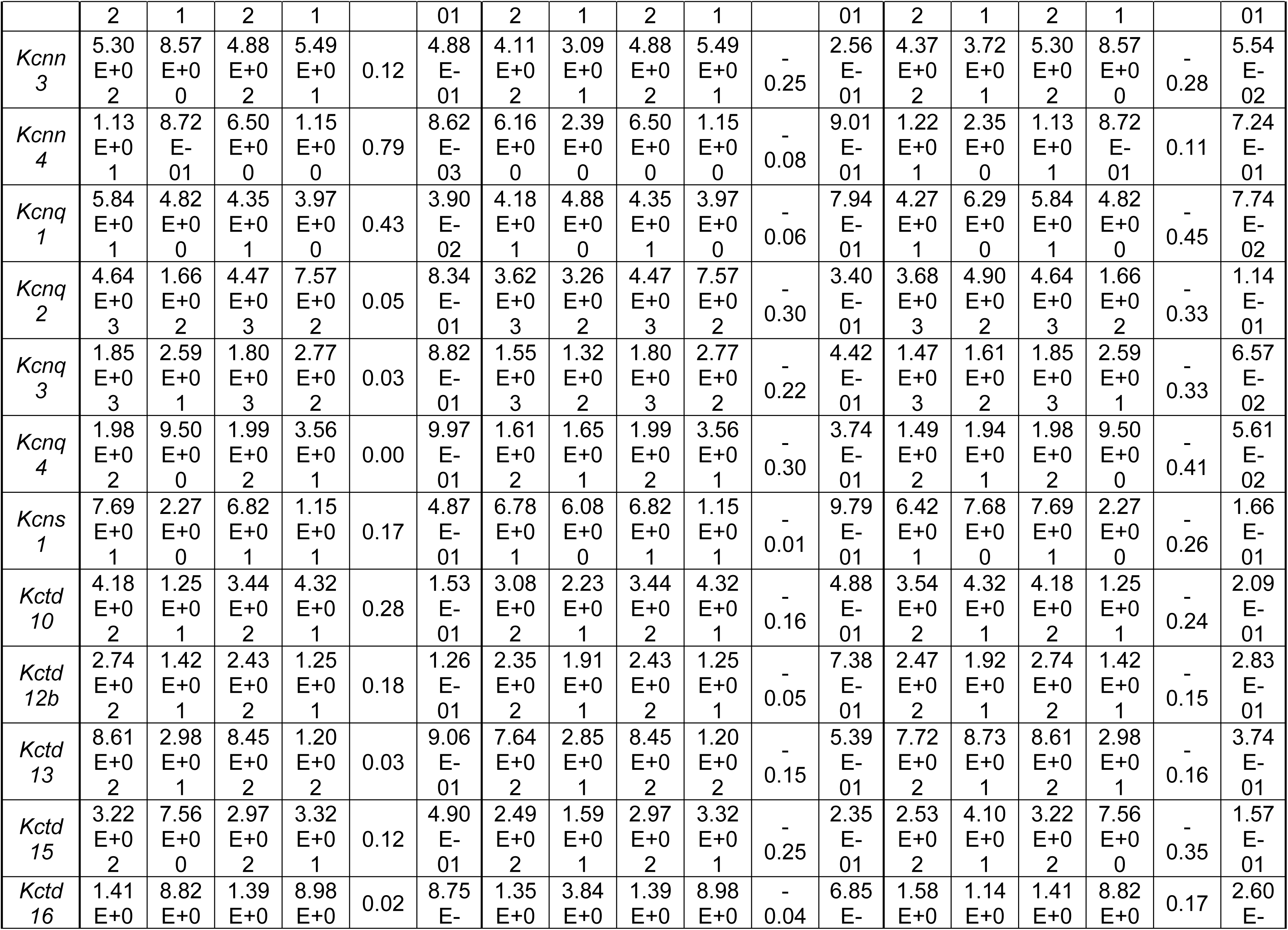
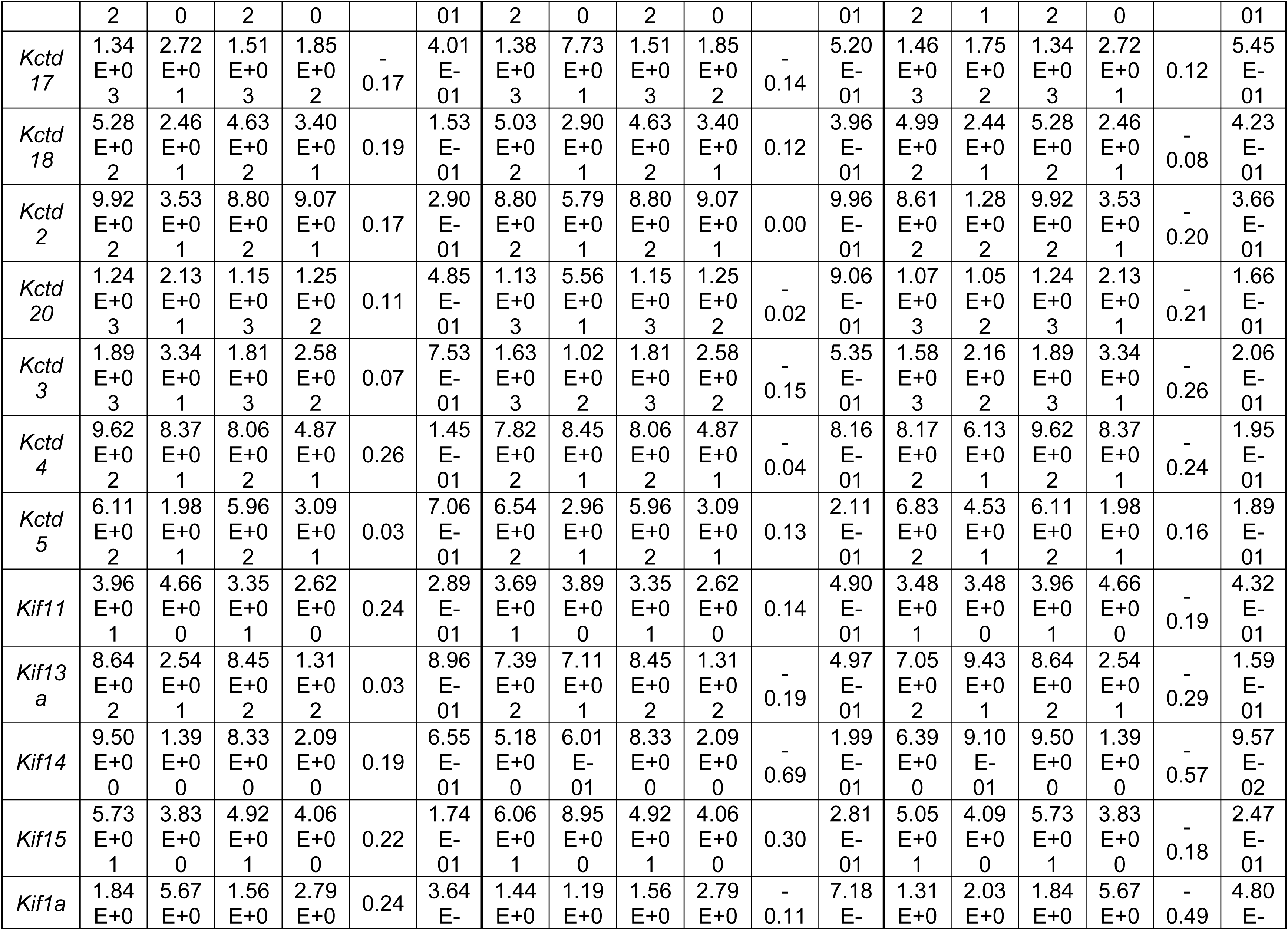
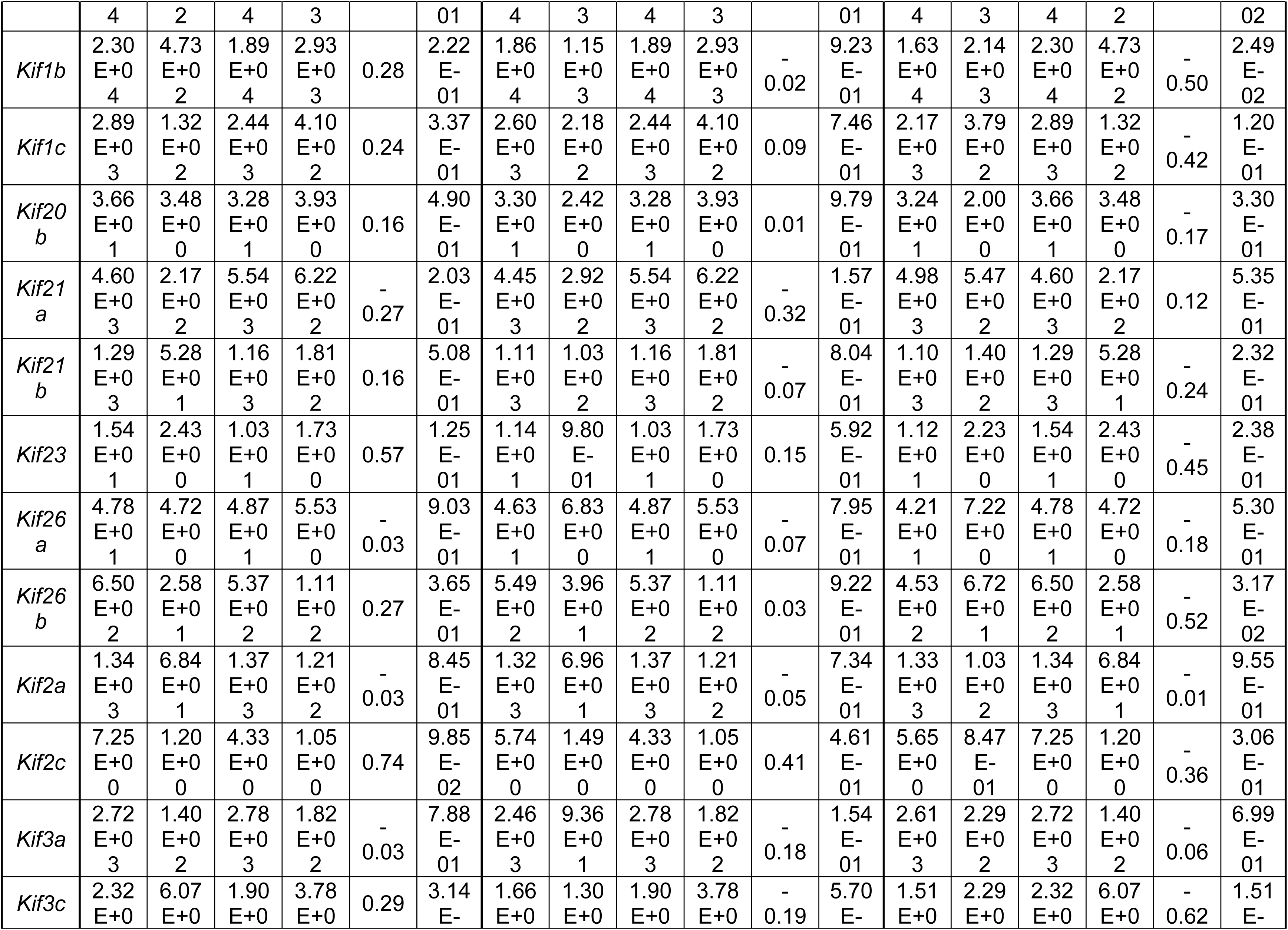
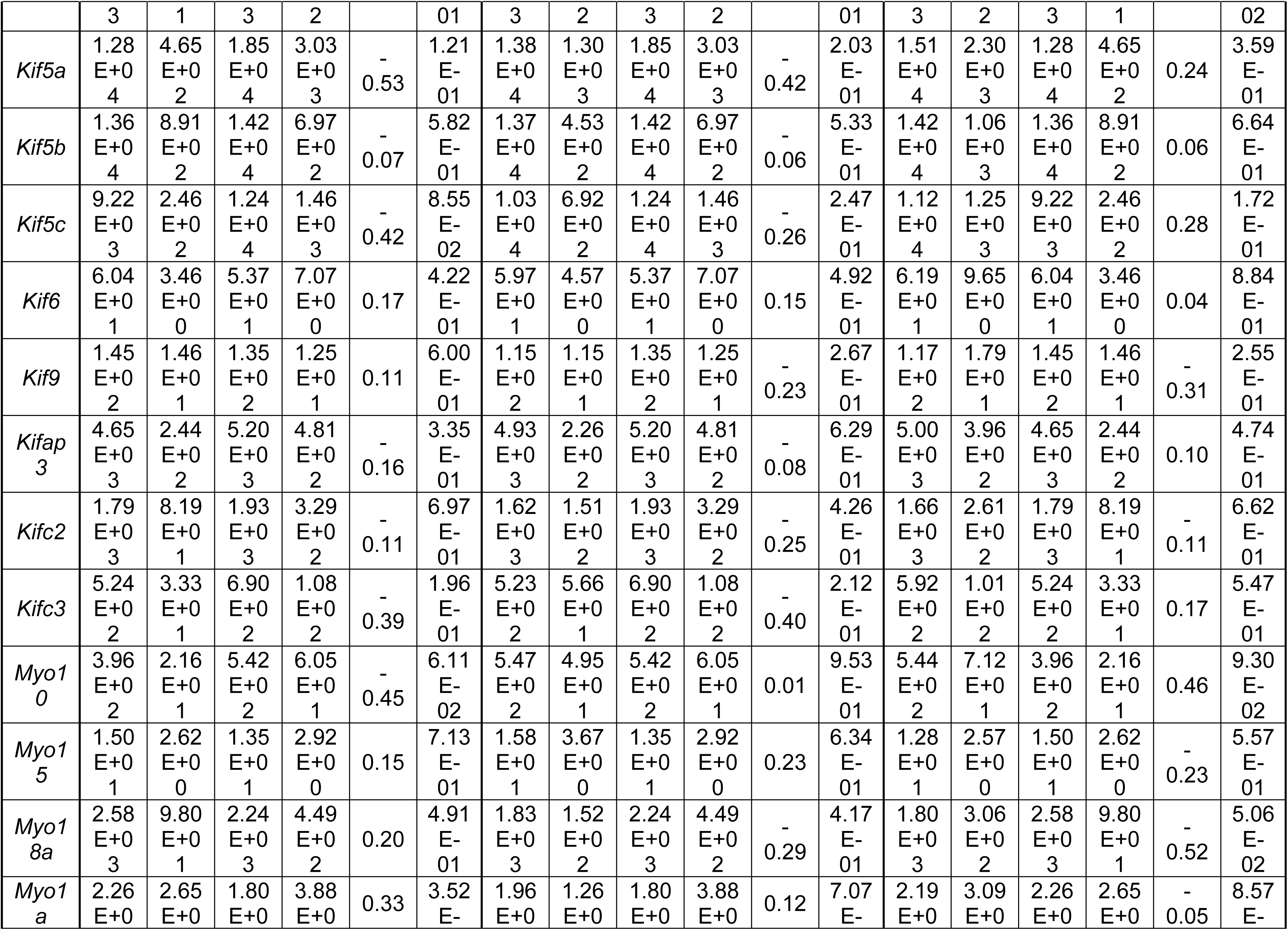
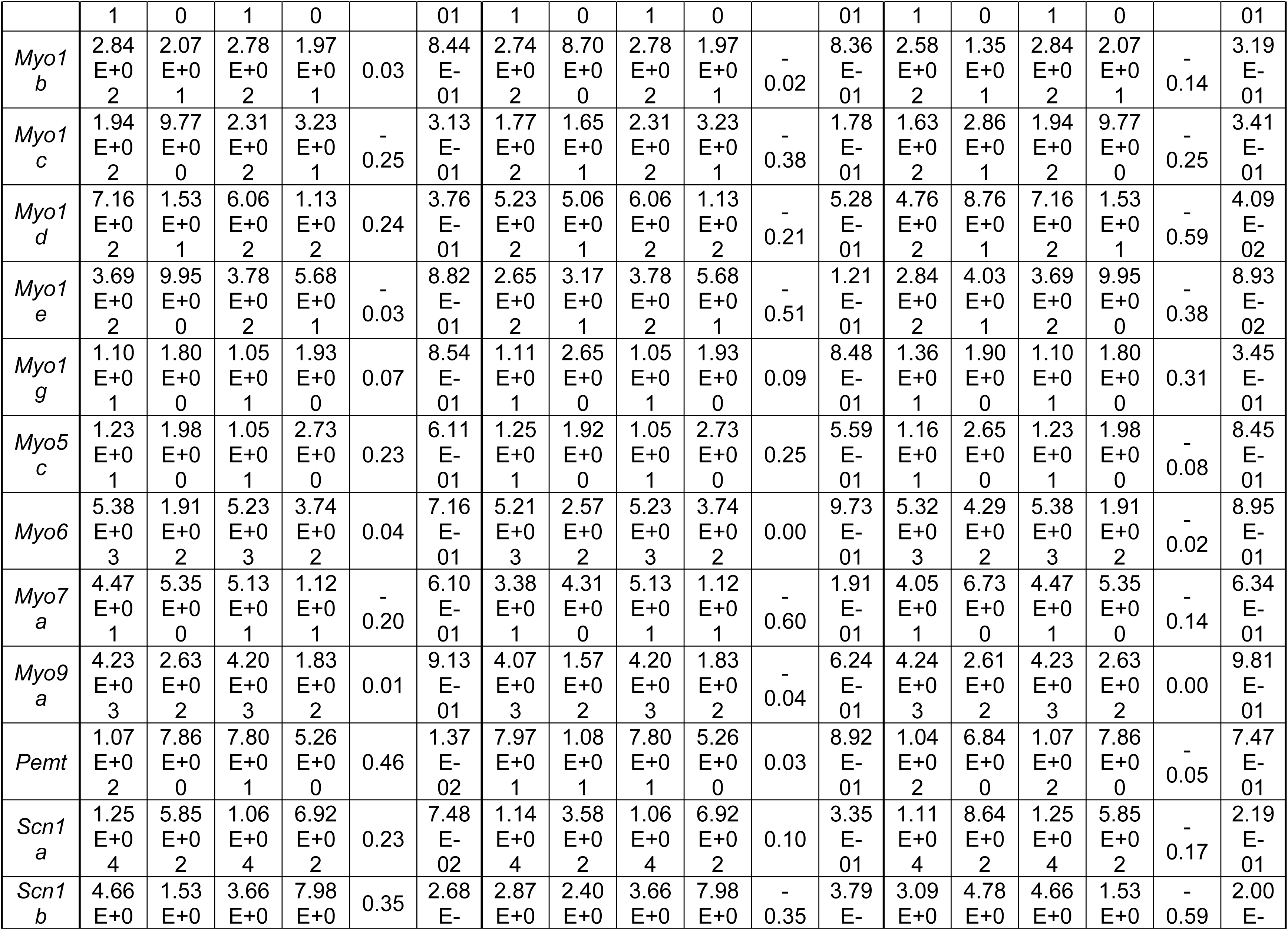
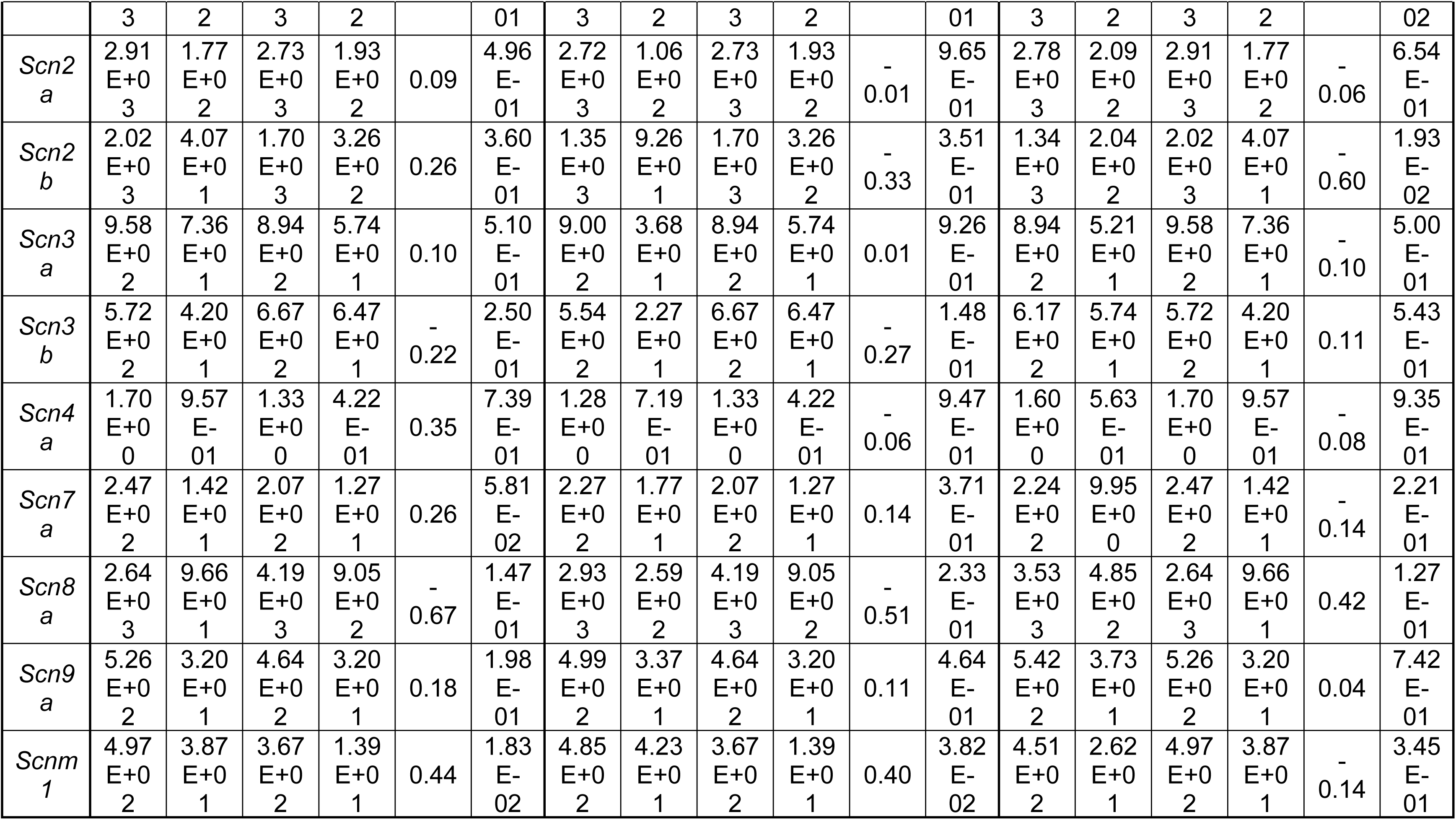
Transcription levels of genes associated with biomass production in ipsilateral L4- L6 spinal cord of vehicle (Veh)- or formalin (Form)-injected mice fed either SD or MD-1. Genes are listed in alphabetical order. Data are expressed as mean±SEM values (counts). FC: fold change.

**Supplemental Table S6.** Serum metabolites influenced by formalin injection, ranked in ascending order of Log_2_ fold change (FC). Data are presented as mean±SEM values (ion counts) for each metabolite. DG: diacylglycerol; PG: phosphatidylglycerol; PE: phosphatidylethanolamine; PC: phosphatidylcholine. FC: fold changes ([Form_SD]/[Veh_SD]).

| Compound Id | Veh_SD |  | Form_SD |  | Log<sub>2</sub> FC | P-value |
| --- | --- | --- | --- | --- | --- | --- |
|  | Mean | SEM | Mean | SEM |  |  |
| DG(16:0/18:2(9Z,12Z)/0:0) | 1.70E+04 | 2.56E+03 | 3.75E+03 | 8.34E+02 | -2.18 | 9.79E-04 |
| Docosatrienoic acid | 1.01E+06 | 9.66E+04 | 4.85E+05 | 5.40E+04 | -1.06 | 5.94E-04 |
| Indole-3-methyl acetate | 3.36E+05 | 7.15E+04 | 1.67E+05 | 2.28E+04 | -1.01 | 4.62E-02 |
| 5-Nonadecyl-1,3-benzenediol | 2.56E+05 | 4.86E+04 | 1.38E+05 | 1.80E+04 | -0.89 | 4.28E-02 |
| Nervonic acid | 8.40E+05 | 1.20E+05 | 4.72E+05 | 5.96E+04 | -0.83 | 1.82E-02 |
| Docosadienoate (22:2n6) | 6.16E+05 | 1.06E+05 | 3.48E+05 | 4.68E+04 | -0.82 | 3.93E-02 |
| Erucic acid | 2.88E+06 | 4.13E+05 | 1.69E+06 | 2.53E+05 | -0.77 | 2.68E-02 |
| Arachidic acid | 1.39E+06 | 2.03E+05 | 8.19E+05 | 1.13E+05 | -0.76 | 2.73E-02 |
| 10Z-Pentadecenoic acid | 1.41E+05 | 2.00E+04 | 9.23E+04 | 9.83E+03 | -0.61 | 4.77E-02 |
| Tetracosahexaenoic acid | 1.66E+06 | 2.17E+05 | 1.10E+06 | 8.29E+04 | -0.6 | 3.28E-02 |
| PG(18:0/16:0) | 4.60E+04 | 5.31E+03 | 3.33E+04 | 1.46E+03 | -0.47 | 4.22E-02 |
| Taurine | 8.47E+07 | 5.72E+06 | 1.07E+08 | 3.13E+06 | 0.34 | 4.15E-03 |
| Butyric acid | 4.34E+04 | 4.60E+03 | 5.61E+04 | 3.83E+03 | 0.37 | 4.79E-02 |
| PC(18:0/18:3(9Z,12Z,15Z)) | 9.29E+05 | 9.06E+04 | 1.25E+06 | 1.13E+05 | 0.43 | 3.77E-02 |
| Pseudouridine | 3.61E+06 | 3.52E+05 | 4.90E+06 | 4.28E+05 | 0.44 | 3.16E-02 |
| Glutaryl carnitine | 2.53E+04 | 2.08E+03 | 3.46E+04 | 3.52E+03 | 0.45 | 3.93E-02 |
| Capryloylglycine | 1.89E+05 | 2.31E+04 | 2.74E+05 | 3.18E+04 | 0.54 | 4.50E-02 |
| 3-methylglutaryl carnitine | 5.94E+04 | 5.87E+03 | 9.89E+04 | 1.25E+04 | 0.73 | 1.35E-02 |
| Aldosterone | 4.87E+05 | 8.77E+04 | 8.71E+05 | 8.22E+04 | 0.84 | 8.42E-03 |
| SM(d18:1/23:0) | 3.78E+04 | 6.33E+03 | 7.48E+04 | 1.22E+04 | 0.98 | 1.78E-02 |
| Corticosterone | 2.81E+04 | 5.11E+03 | 6.68E+04 | 1.39E+04 | 1.25 | 2.39E-02 |
| Indole-3-carboxylic acid-sulphate isomer | 7.13E+05 | 1.08E+05 | 1.77E+06 | 2.47E+05 | 1.31 | 2.01E-03 |
| PC(18:1(9Z)/16:0) | 7.02E+05 | 2.85E+05 | 2.36E+06 | 6.46E+05 | 1.75 | 3.59E-02 |

**Supplemental Table S7.** ANCOVA analysis of the effects of MD-1 on ipsilateral and contralateral thermal and mechanical hypersensitivity, adjusted for body weight (BW) variations. Results show the source of variation, Type III Sum of Squares, degrees of freedom (df), Mean Square, F-value, significance level (Sig.), and Partial Eta Squared. Statistical significance is set at *P*<0.05.

| Dependent Variable | Source | Type III Sum of Squares | df | Mean Square | F | Sig. | Partial Eta Squared |
| --- | --- | --- | --- | --- | --- | --- | --- |
| Thermal (Ipsilateral) | Corrected Model | 554.178 | 6 | 92.363 | 29.654 | <.001 | 0.813 |
|  | Intercept | 2358.363 | 1 | 2358.363 | 757.175 | <.001 | 0.949 |
|  | BW | 10.658 | 1 | 10.658 | 3.422 | 0.072 | 0.077 |
|  | MD-1 | 481.944 | 2 | 240.972 | 77.366 | <.001 | 0.791 |
|  | PFD | 22.214 | 1 | 22.214 | 7.132 | 0.011 | 0.148 |
|  | MD-1 * PFD | 17.788 | 2 | 8.894 | 2.856 | 0.069 | 0.122 |
|  | Error | 127.702 | 41 | 3.115 |  |  |  |
|  | Total | 3129.472 | 48 |  |  |  |  |
|  | Corrected Total | 681.88 | 47 |  |  |  |  |
| Thermal (Contralateral) | Corrected Model | 410.877 | 6 | 68.479 | 13.637 | <.001 | 0.666 |
|  | Intercept | 3381.238 | 1 | 3381.238 | 673.337 | <.001 | 0.943 |
|  | BW | 5.594 | 1 | 5.594 | 1.114 | 0.297 | 0.026 |
|  | MD-1 | 336.72 | 2 | 168.36 | 33.527 | <.001 | 0.621 |
|  | PFD | 0.068 | 1 | 0.068 | 0.014 | 0.908 | 0 |
|  | MD-1 * PFD | 0.919 | 2 | 0.459 | 0.091 | 0.913 | 0.004 |
|  | Error | 205.886 | 41 | 5.022 |  |  |  |
|  | Total | 4024.143 | 48 |  |  |  |  |
|  | Corrected Total | 616.763 | 47 |  |  |  |  |
| Mechanical (Ipsilateral) | Corrected Model | 96.994 | 6 | 16.166 | 65.436 | <.001 | 0.905 |
|  | Intercept | 367.227 | 1 | 367.227 | 1486.476 | <.001 | 0.973 |
|  | BW | 0.027 | 1 | 0.027 | 0.109 | 0.744 | 0.003 |
|  | MD-1 | 69.051 | 2 | 34.526 | 139.754 | <.001 | 0.872 |
|  | PFD | 8.695 | 1 | 8.695 | 35.198 | <.001 | 0.462 |
|  | MD-1 * PFD | 8.428 | 2 | 4.214 | 17.058 | <.001 | 0.454 |
|  | Error | 10.129 | 41 | 0.247 |  |  |  |
|  | Total | 482.211 | 48 |  |  |  |  |
|  | Corrected Total | 107.122 | 47 |  |  |  |  |
| Mechanical (Contralateral) | Corrected Model | 67.359 | 6 | 11.227 | 15.798 | <.001 | 0.698 |
|  | Intercept | 517.257 | 1 | 517.257 | 727.874 | <.001 | 0.947 |
|  | BW | 1.272 | 1 | 1.272 | 1.79 | 0.188 | 0.042 |
|  | MD-1 | 65.028 | 2 | 32.514 | 45.753 | <.001 | 0.691 |
|  | PFD | 0.562 | 1 | 0.562 | 0.791 | 0.379 | 0.019 |
|  | MD-1 * PFD | 0.261 | 2 | 0.13 | 0.184 | 0.833 | 0.009 |
|  | Error | 29.136 | 41 | 0.711 |  |  |  |
|  | Total | 630.762 | 48 |  |  |  |  |
|  | Corrected Total | 96.495 | 47 |  |  |  |  |

**Supplemental Table S8.**
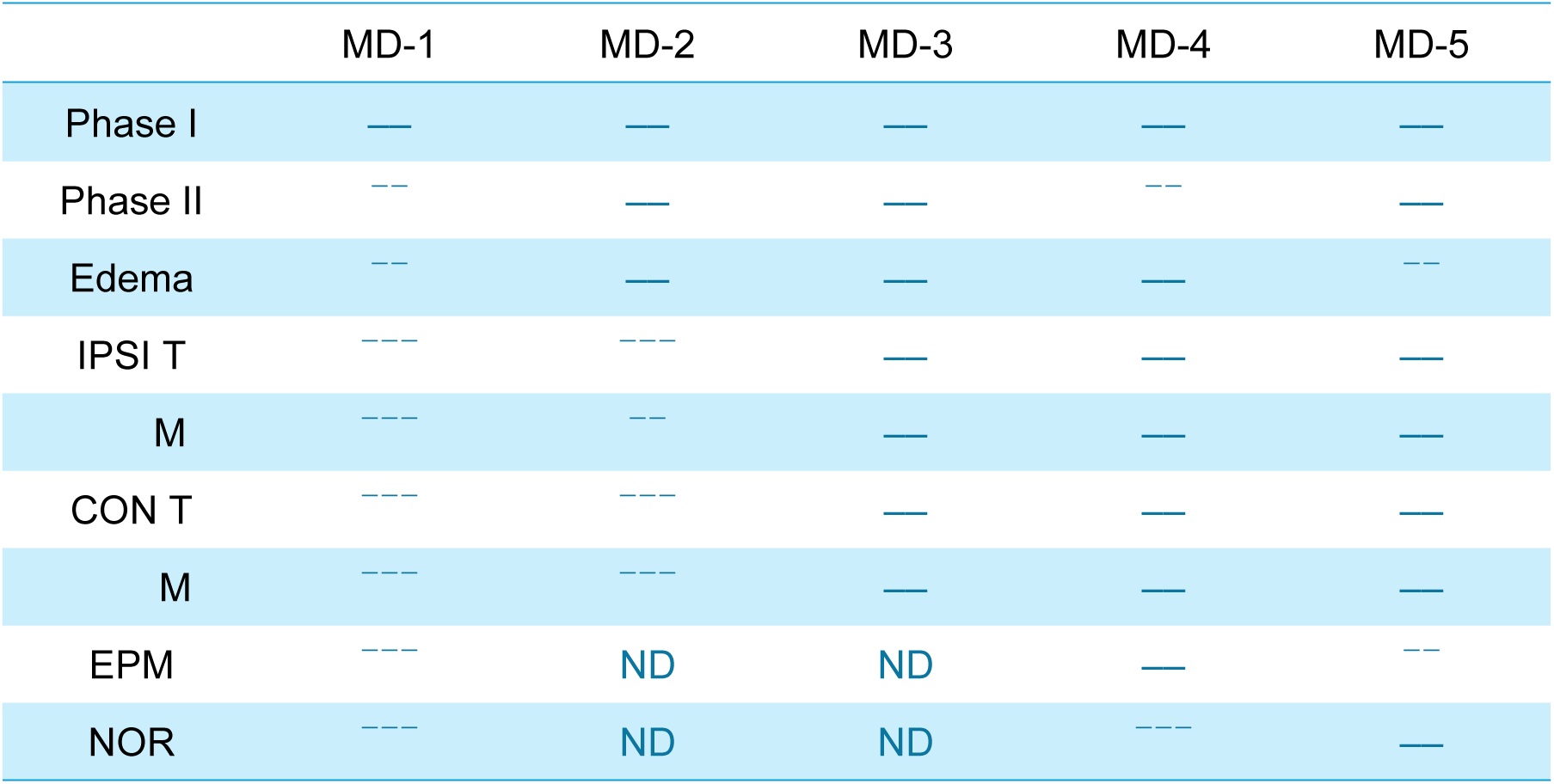
Impact of modified diets on formalin-induced acute nociception (phase I and II), paw edema, contralateral (CON) and ipsilateral (IPSI) hypersensitivity to thermal (T) or mechanical (M) stimuli (assessed on days 14 and 21 post-injury), anxiety-like behavior (elevated-plus maze, EPM), and long-term memory deficits (24-hour novel-object recognition, NOR). Arrows symbolize percent of maximum possible effect (MPE) produced by the modified diet on the formalin response (for effects with *P* < 0.05): ^-^ <20% MPE; ^--^ 20-80% MPE; ^---^ >80% MPE. Horizontal lines indicate no change or change with P ³ 0.05.

